# Extensive exploration of structure activity relationships for the SARS-CoV-2 macrodomain from shape-based fragment merging and active learning

**DOI:** 10.1101/2024.08.25.609621

**Authors:** Galen J. Correy, Moira Rachman, Takaya Togo, Stefan Gahbauer, Yagmur U. Doruk, Maisie Stevens, Priyadarshini Jaishankar, Brian Kelley, Brian Goldman, Molly Schmidt, Trevor Kramer, Alan Ashworth, Patrick Riley, Brian K. Shoichet, Adam R. Renslo, W. Patrick Walters, James S. Fraser

## Abstract

The macrodomain contained in the SARS-CoV-2 non-structural protein 3 (NSP3) is required for viral pathogenesis and lethality. Inhibitors that block the macrodomain could be a new therapeutic strategy for viral suppression. We previously performed a large-scale X-ray crystallography-based fragment screen and discovered a sub-micromolar inhibitor by fragment linking. However, this carboxylic acid-containing lead had poor membrane permeability and other liabilities that made optimization difficult. Here, we developed a shape-based virtual screening pipeline - FrankenROCS - to identify new macrodomain inhibitors using fragment X-ray crystal structures. We used FrankenROCS to exhaustively screen the Enamine high-throughput screening (HTS) collection of 2.1 million compounds and selected 39 compounds for testing, with the most potent compound having an IC_50_ value equal to 130 μM. We then paired FrankenROCS with an active learning algorithm (Thompson sampling) to efficiently search the Enamine REAL database of 22 billion molecules, testing 32 compounds with the most potent having an IC_50_ equal to 220 μM. Further optimization led to analogs with IC_50_ values better than 10 μM, with X-ray crystal structures revealing diverse binding modes despite conserved chemical features. These analogs represent a new lead series with improved membrane permeability that is poised for optimization. In addition, the collection of 137 X-ray crystal structures with associated binding data will serve as a resource for the development of structure-based drug discovery methods. FrankenROCS may be a scalable method for fragment linking to exploit ever-growing synthesis-on-demand libraries.

## Introduction

Since 2020, our combined team of structural biologists and medicinal and computational chemists has been working to advance hits emerging from a large-scale X-ray-based fragment screen targeting the SARS-CoV-2 NSP3 macrodomain (Mac1, **Fig. 1A**) (*1*, *2*). One fruitful computational avenue towards this end has been our use of the fragment-linking method Fragmenstein (*2*, *3*). This approach links fragments *in silico* and uses the resulting larger molecules to search for related commercial or tangible (readily synthesized) analogs (*3*).

**Figure 1.**
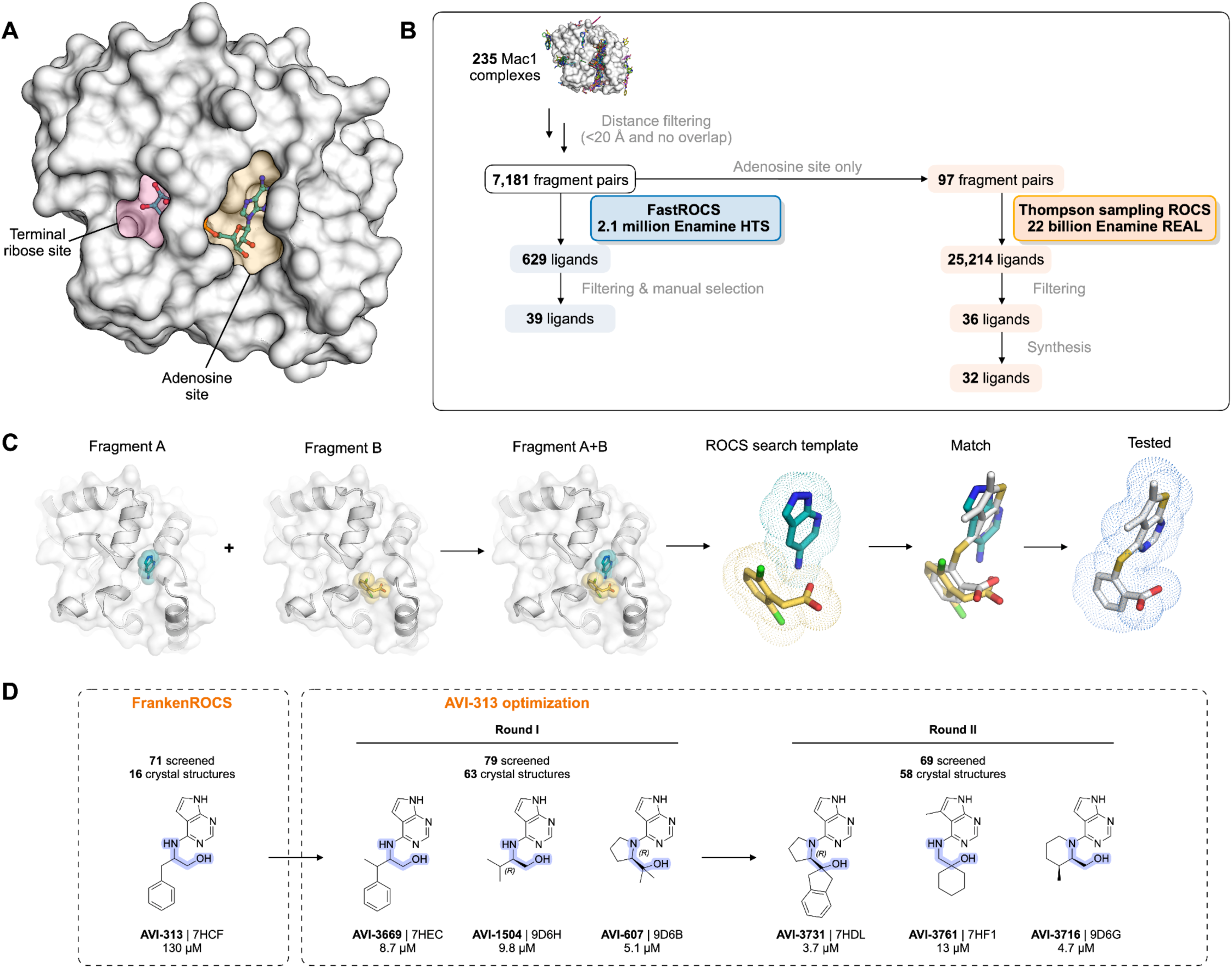
The FrankenROCS workflow. (**A**) The ADP-ribose binding site of the SARS-CoV-2 macrodomain (Mac1) with the terminal ribose site (pink shaded) and the fragment-rich adenosine site (yellow shaded) highlighted. (**B**) An overview of the FrankenROCS workflow. Beginning with 235 fragments, 7181 pairs of fragments were constructed and used to exhaustively search the Enamine HTS collection. A limited set of 97 fragment pairs were used to search the much larger Enamine REAL database using Thompson sampling. (**C**) The binding poses of two fragments are shown in the context of their individually determined structures and merged to make the ROCS search template. The match from the ROCS search is shown in gray superposed with the parent fragments, which are shown in color. After filtering, the most promising matches are experimentally tested. (**D**) Optimization of a non-carboxylic acid hit (AVI-313) identified from the exhaustive search of the Enamine HTS collection identified several compounds with low-micromolar potency. The 2- aminoethanol core of the compounds is highlighted blue. PDB codes are indicated and the IC_50_ values from a homogeneous time resolved fluorescence (HTRF) assay are indicated (best fit value of three technical replicates).

Importantly, by searching the Enamine REAL database using the 2D molecular similarity search engine SmallWorld (http://sw.docking.org) and the substructure browser Arthor (http://arthor.docking.org) (*4*), Fragmenstein is able to efficiently search a large portion of readily available chemical space. These search methods ensure a high similarity of the merged compounds with the parent fragments, but are correspondingly limited in their exploration of different scaffolds. For Mac1, the best analogs emerging from this approach had sub-micromolar affinity, but used a carboxylic acid to engage the Mac1 active site, as this was a common feature of the fragment hits (*1*, *2*). Thus, while Fragmenstein yielded linked molecules with improved potencies, these compounds retained many of the liabilities (e.g. poor cellular permeability, sites for secondary metabolism) inherited from their progenitor fragments.

While fragment screening, as done for Mac1, is a powerful tool in drug discovery, it requires iteratively increasing potency from initial weak (∼mM) hits (*5*). This process normally involves significant medicinal chemistry effort. There are typically two paths for increasing affinity after a fragment screen. The fragment growing strategy uses visual inspection and computational evaluation to design larger analogs, generally from a single fragment hit (*6*). Such analogs can make additional interactions with the receptor, thus increasing the binding affinity. The computational tools for fragment growing resemble those used for analog design from other structurally-enabled screening approaches. The fragment linking strategy involves connecting two experimentally validated fragments to generate a new molecule (*7*). In some cases, the binding affinity of the linked fragment can be orders of magnitude more potent than either of the starting fragments (*8*). Several computational fragment-linking approaches have been published with different emphases on linker design, molecular strain, fidelity to the original binding poses, and synthetic tractability (*7*).

With the explosion of synthesis-on-demand chemistry and the massive expansion of accessible chemical space, several new approaches are emerging to efficiently identify and source new analogs (*9*). V-SYNTHES is a hierarchical synthon-based computational screening method that mirrors the fragment-growing process by elaborating fragment-sized initial seed hits to complete molecules based on docking scores (*10*). This approach identified potent hits from a large chemical space while docking only a small fraction of total library compounds. Another recent computational approach that more closely resembles fragment merging leverages unsupervised machine learning to identify significant pharmacophore distributions across hits from X-ray fragment screens (*11*). By evaluating potential molecules based on their recapitulation of fragment-derived pharmacophore distributions, the approach can identify alternative scaffolds that are likely to bind the receptor. An example of the advantages of the pharmacophore approach was the development of a ∼100 μM Mac1 inhibitor in which the carboxylic acid found in most input fragments was replaced with an amide (*11*). More classic medicinal-chemistry using carboxylate bioisosteres can also lead to relatively potent inhibitors. In the case of Mac1 a pyrrolidinone-containing compound (LRH-0003, denoted AVI-219 herein) was identified that bound at 1.7 μM (*2*). Collectively, these, and other related strategies (*9*), suggest that experimentally-derived fragment binding poses can be used to accelerate computational searches of massive chemical space for potent and diverse compounds.

Here, we develop a new approach for fragment linking - FrankenROCS - that uses an active learning method, Thompson sampling (TS), to efficiently search over 22 billion synthesis-on-demand molecules in the Enamine REAL database (**Fig. 1B**). FrankenROCS takes pairs of fragments as input to query a database using the <u>R</u>apid <u>O</u>verlay of <u>C</u>hemical <u>S</u>tructures (ROCS) method of comparing 3D shape and pharmacophore distribution (*12*) (**Fig. 1C**). By using the ROCS shape-based approach, we avoid the problem of explicitly defining a chemical linker or functional group to anchor the new designs in the original fragments’ crystallographic orientations and interactions. We first exhaustively searched the Enamine HTS collection of 2.1 million compounds to find initial hits using FrankenROCS. While an equivalent ROCS search of the >22 billion virtual molecules in Enamine REAL would have been prohibitive, Thompson sampling (TS) in reagent space allowed us to search the REAL database at a reasonable speed and cost. Notably, this approach delivered multiple neutral Mac1 inhibitors with diverse binding poses determined by X-ray crystallography. After optimization of the most potent hit, several low-micromolar compounds were identified that were ligand efficient, had favorable drug-like properties and represent improved starting points for lead optimization. Moreover, the 137 high resolution X-ray crystal structures, and corresponding affinity measurements described herein, may provide a valuable benchmark for further fragment methods development.

## Results

### FrankenROCS identifies alternative scaffolds in the adenosine site of Mac1 from the Enamine HTS collection

We constructed each FrankenROCS query from pairs of aligned fragments and exhaustively searched the 2.1 million Enamine HTS collection (Methods, **Fig. 1B,C** and **Table S1**). We ranked matched compounds by the Tanimoto Combo (TC) score that evaluates the steric overlap and the color score relative to the paired fragments. After manually inspecting the top 1000 ligands, we selected 39 to purchase (**Table S2**A,B). The final selection weighed several factors including overlap of key pharmacophoric elements between the query and database molecules, complementarity with the protein binding site, avoidance of steric clashes, and a manual assessment of ligand strain.

To evaluate the binding of the FrankenROCS generated compounds to Mac1, we soaked the 39 compounds into crystals of the enzyme and collected X-ray diffraction data using the previously-validated P4_3_ crystal form and soaking procedure (*1*, *2*) (**Table S2**C-E). Binding events in electron density maps were detected and modeled using the PanDDA algorithm (*13*). Structures were determined for 10 of the compounds, with seven bound in the adenosine site and three in distal sites (**Fig. 2A**, **Fig. S1**, **Fig. S2**). The overlap between the parent fragments and the crystallographic poses was high (**Fig. 2B**). A notable exception was AVI-313, where the phenyl ring was shifted by 6 Å relative to the parent fragment (**Fig. 2B**). AVI-313 formed two hydrogen bonds to the adenine subsite, closely matching the parent fragment, however the hydroxyl formed a hydrogen bond with the backbone oxygen of Leu126 instead of the oxyanion subsite (**Fig. 2B**). AVI-313 was purchased as a racemic mixture, and while both isomers were observed in the crystal structure, the *S*-isomer was the major species (∼70%) (**Fig. S2**). Although the position of the *para*-carbon is almost identical, the *R*-isomer phenyl is ∼1 Å closer to the oxyanion subsite. The position of the AVI-313 *S*-isomer phenyl ring matched the ring of AVI-321 (**Fig. 2B**), where the parent fragments feature a furan ring and a trifluoromethyl group. This substitution illustrates the power of shape matching for scaffold hopping (*14*).

**Figure 2.**
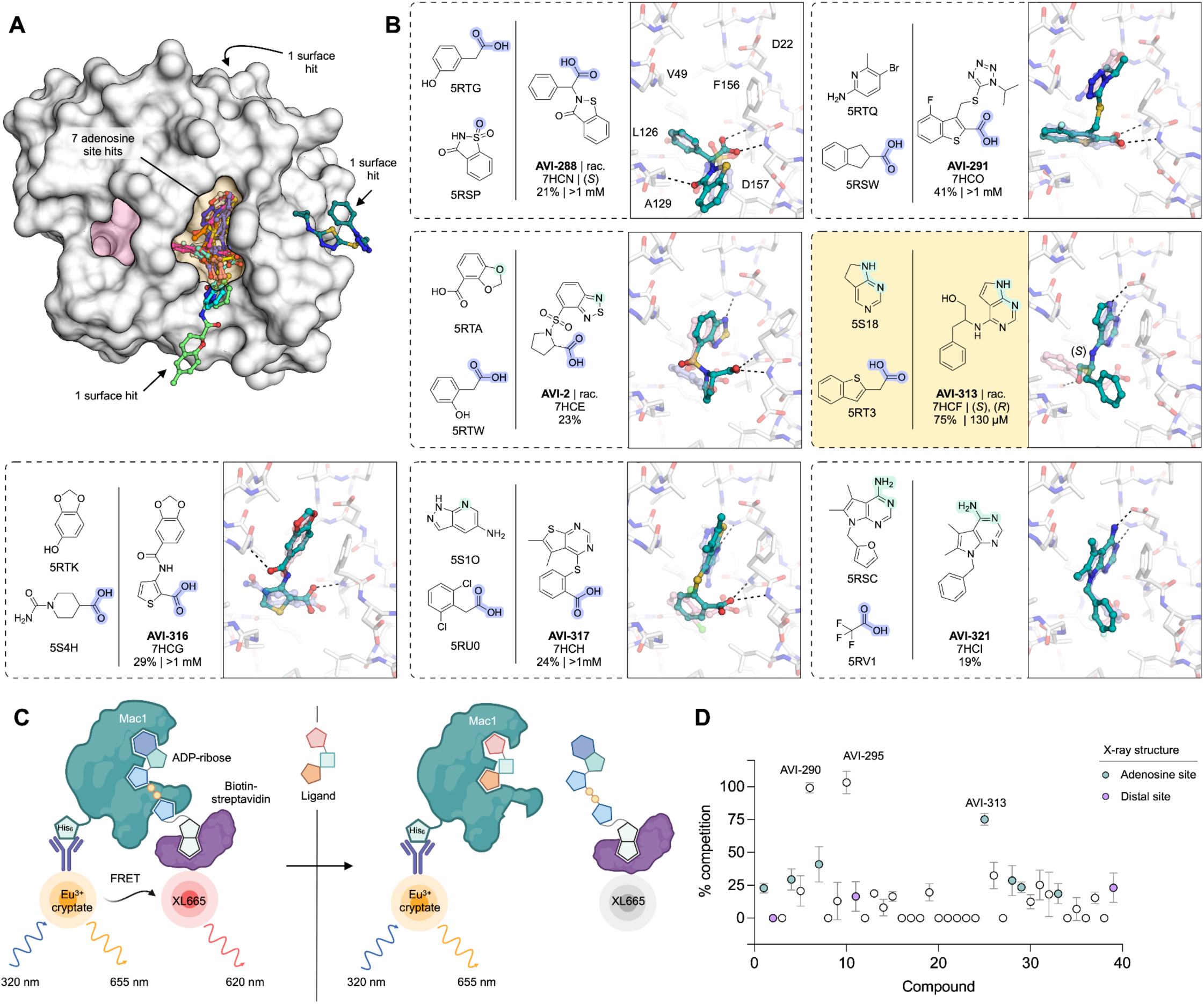
Structural and biophysical characterization of compounds from the FrankenROCS pipeline. (**A**) Surface view of Mac1 showing the binding site of the 10 FrankenROCS hits. (**B**) Chemical structures of parent fragments and ligands identified by FrankenROCS. Compounds were purchased as pure isomers or as racamates (rac.). The stereochemistry of the modeled compound is indicated. The X-ray crystal structure of the FrankenROCS ligand bound to Mac1 (teal sticks) is overlaid with the X-ray crystal structures of the parent fragments (transparent blue/pink sticks). Hydrogen bonds between the FrankenROCS ligand and Mac1 are shown as dashed black lines. Results for the preliminary HTRF screen (% competition at 200 μM, mean of 3-4 technical replicates) and the confirmatory dose-response screen (IC_50_, best fit value of three technical replicates) are indicated. The AVI-313 chemical structure and parent fragments are highlighted (yellow box). (**C**) Schematic of the HTRF peptide displacement assay used to measure ligand binding to Mac1. (**D**) Plot showing % competition in the HTRF assay for the 39 compounds from FrankenROCS. Compounds were assayed at 200 μM and data are presented ± standard deviation (SD) of 3-4 technical replicates.

We observed a higher hit rate in the adenosine site for carboxylic acid containing compounds (5/11 compounds, 45% hit rate) versus non-carboxylic acid containing compounds (2/28 compounds, 7% hit rate). Overall, the X-ray crystal structures show that the compounds identified by FrankenROCS generally capture the key protein interactions of the parent fragments and confirm the propensity of carboxylic-acid-containing compounds to bind to the oxyanion subsite (*1*, *2*). Although all of the parent fragments formed two hydrogen bonds with the oxyanion subsite, only two of the linked fragments formed two hydrogen bonds (AVI-288 and AVI-291, **Fig. 2B**). The two hits that did not contain carboxylic acids, AVI-313 and AVI-321, are of great interest for subsequent medicinal chemistry efforts as they could overcome the permeability liabilities observed in previously identified carboxylic acid containing compounds (*2*).

Next, we measured solution binding of the compounds using a homogeneous time resolved fluorescence (HTRF) competition assay with an ADP-ribose conjugated peptide (*1*, *15*) (**Fig. 2C**, **Fig. S3**A). Compounds were tested in triplicate at 200 μM and IC_50_ values were determined for compounds with >40% competition (**Fig. 2D**). The most potent were a pair of tri-substituted pyridines (AVI-290 and AVI-295) (**Fig. S4**A), which IC_50_ values equal to 56 and 52 μM respectively. We failed to observe binding of these compounds by crystal soaking, despite repeated attempts (**Table S2**C). The failure might be due to compounds being unable to bind in the P4_3_ crystal form, or compound-induced assay interference. The next most potent of the compounds was AVI-313 (IC_50_ = 130 μM). In summary, the FrankenROCS approach resulted in seven new compounds binding in the adenosine site with affinities approaching 100 μM. Two of these compounds lacked a carboxylic acid and are therefore potential starting points for the design of neutral inhibitors of Mac1.

### Using Thompson sampling (TS) to search Billions of molecules with FrankenROCS

While the search of the 2.1 million molecule Enamine HTS collection resulted in promising starting points, we were interested in whether distinct chemical series could be uncovered by searching a larger database of molecules. The Enamine REAL database, which at the time this work was performed consisted of more than 22 billion molecules, is a synthesis-on-demand collection consisting of compounds that can be synthesized and made available at a reasonable cost in 6-8 weeks. While databases consisting of millions of molecules, like the Enamine HTS collection, can be exhaustively evaluated by FrankenROCS, the computational and disk space requirements for searching 22 billion molecules necessitated an alternative approach.

To overcome these computational limitations, we used an active learning method known as Thompson sampling (TS) (*16*). TS makes use of the combinatorics encoded within the REAL database, which leads to synthesizable molecules that are constructed from two or three chemical building blocks using compatible reactions. When this work was performed, the molecules in the REAL database were constructed from 143 chemical reactions and 960,398 unique building synthons. In the case of a two-component reaction, the first building block has many types (R1_1… R1_N) that can be combined with any of the second building block types (R2_1…R2_M), producing N*M reaction products. For example, consider the case of a two-component amide bond forming reaction (**Fig. 3A**). Combining 1,158 primary or secondary amine reagents (R1_1…R1_1158) and 15,439 carboxylic acid building blocks (R2_1…R2_15439) generates almost 18 million amide products. By extension, a three-component reaction that combines 1,000 of each of the three building blocks will yield 1 billion reaction products. With these large numbers of potential products, searching exhaustively is impractical.

**Figure 3.**
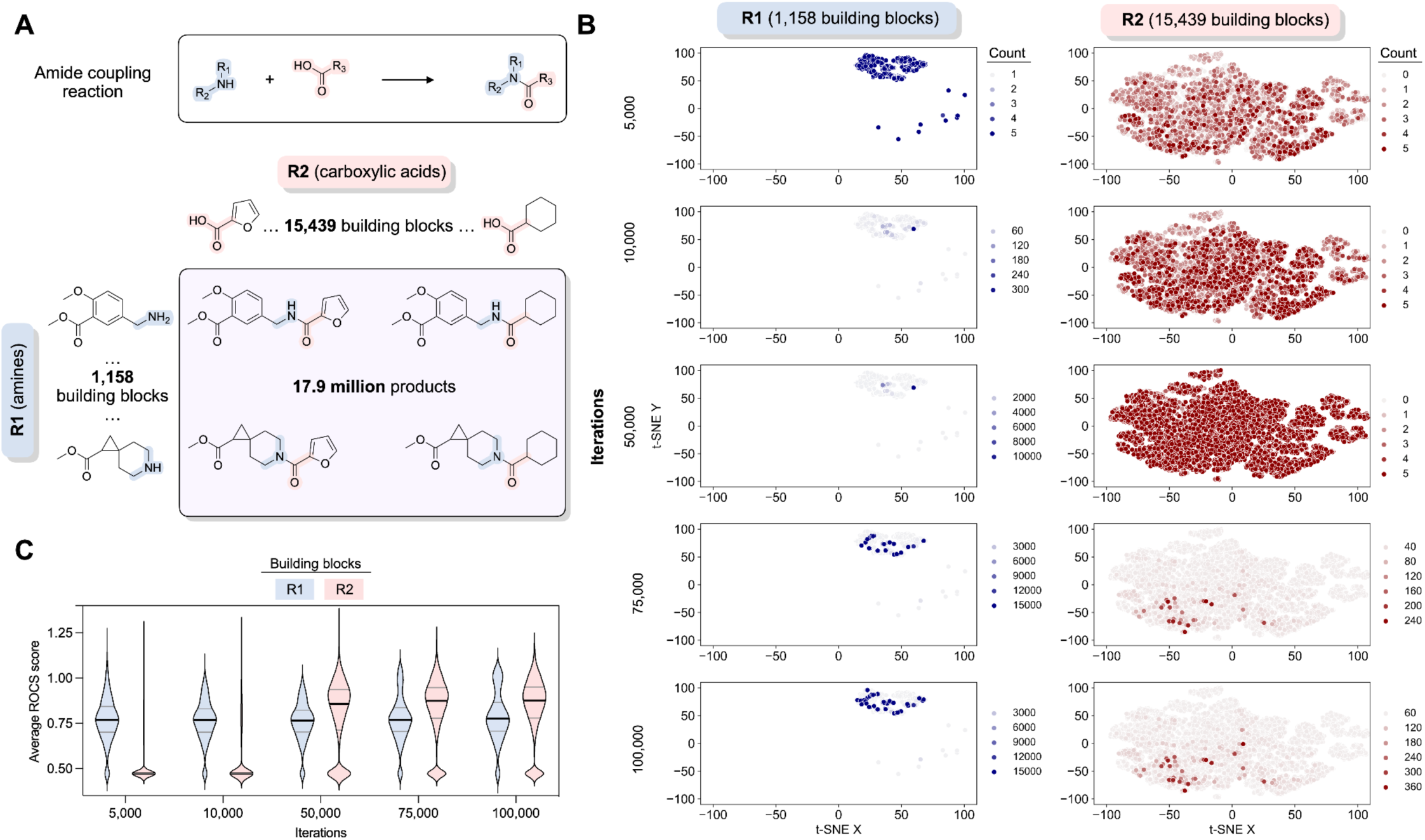
Thompson sampling in reaction space identified potential new Mac1 ligands. (**A**) The relationship between the reaction, reagents, and products used in the Thompson sampling illustrated for an amide coupling reaction that can generate 17.9 million potential products. (**B**) The chemical space covered by two sets of reagents using t-Distributed Stochastic Neighbor Embedding (t-SNE) after a given number of iterations during a Thompson sampling run. Within each chemical space plot, each point represents a molecule, with similar molecules closer together and less similar molecules farther apart. The color of the points represents the number of times a particular reagent has been sampled up to the indicated iteration. The plots show how the algorithm begins with an exploration phase and then moves to an exploitation strategy that focuses on specific regions of chemical space. (**C**) The evolution of average score distributions associated with reagent building blocks (R1 and R2) from the Thompson Sampling run shown in (**B**). As there were fewer R1 reagents, these building blocks were more thoroughly sampled at earlier iterations. After approximately 50,000 iterations, R2 has been fully sampled, and the score distribution for R2 shifts to higher values. Between 75,000 and 100,000 iterations, there are minimal changes in the score distributions for R1 and R2, ending the run.

Thompson sampling achieves efficiency gains by searching in reagent space rather than in product space. As an example, TS for the amide coupling reaction with 1,158 R1 reagents and 15,439 R2 reagents (**Fig. 3A**) begins with a warm-up phase where each reagent is sampled at least 3 times (**Fig. 3B,C**). Each R1 is combined with 3 randomly selected R2 reagents over a total of 3,474 iterations. Following this step, each R2 reagent is sampled with 3 randomly selected R1 reagents, consisting of an additional 46,317 iterations. During this warm-up period of R2, clear frontrunners of R1 begin to emerge. After this first 49,791 (1,158*3+15,439*3) iterations of warm-up, the TS algorithm can fully move from exploration to exploitation for both reagents and intensify sampling to yield products that are more likely to be highly scored. Over the next 25,000 iterations, clear frontrunners emerge for R2 while R1 sampling diversifies slightly. There is little change between 75,000 and 100,000 iterations (**Fig. 3B,C**), meeting the convergence criteria of no improvement over 0.01% of the library. Where the convergence criteria is not met after 0.1% of the total library has been scored, we also stop sampling for those reactions. Based on prior comparisons between TS and exhaustive sampling (*16*), this procedure has likely identified most of the top scoring compounds, by ROCS, from R1 and R2 while only calculating scores for a small fraction of potential products.

In our TS-FrankenROCS search of the REAL database, we limited this search to 97 queries of fragment pairs bound in pockets adjacent to the adenine binding pocket (**Fig. 1B**). To identify the most promising compounds, we augmented the filters used above with additional automated filtering steps (Methods, **Table S1**). After clustering similar molecules and choosing the ones with the best scores, we identified 36 promising match molecules, of which 32 were synthesized for subsequent crystallographic and binding studies (**Fig. 1B**). We note that these 36 compounds were not selected through manual inspection of 100s of possible, high-scoring compounds, but rather from the improved filtering with no human/expert intervention required to filter out “obvious” artifacts.

### TS-FrankenROCS identifies sulfonamides binding to the oxyanion subsite

We soaked 32 compounds identified by TS-FrankenROCS into Mac1 crystals. Structures were determined for six of the compounds, all binding in the adenosine site (**Fig. 4A**, **Fig. S2**). The overlap between the parent fragments and the poses determined by X-ray crystallography was generally high (**Fig. 4B**). The most dissimilar were AVI-338 and AVI-345, where the 3,4-dihydro-quinoline and 2-amino-pyridine groups were bound outside of the adenine subsite (**Fig. 4B**). Despite these structural differences, all six compounds formed hydrogen bonds with the oxyanion site that matched the parent fragments, four via carboxylic acids and two via sulfonamides (**Fig. 4B**). One of the sulfonamide compounds, AVI-345, resulted from a scaffold hop from a carboxylic acid parent. There was a preference for carboxylic acids (4/9 structures, 44% hit rate) over non- carboxylic acids (2/23 structures, 9% hit rate). The most potent compound in the HTRF assay was AVI-328, with an IC_50_ value of 220 μM (**Fig. 4B,C**, **Fig. S3**A). The carboxylic acid of AVI-328 formed two hydrogen bonds with the oxyanion subsite, closely matching the parent fragment. However, the hydrogen bond between the parent fragment and the Asp22 side-chain is not present in AVI-328, suggesting that affinity could be improved by modifications to the 3,4-dihydroquinolinone portion. AVI-328 was synthesized as a racemate, and both the *R* and *S* isomers bound in the X-ray crystal structure (**Fig. S2**). The *R* isomer more closely matched the parent fragments, while the *S* isomer was shifted closer to the adenine subsite (**Fig. S2**).

**Figure 4.**
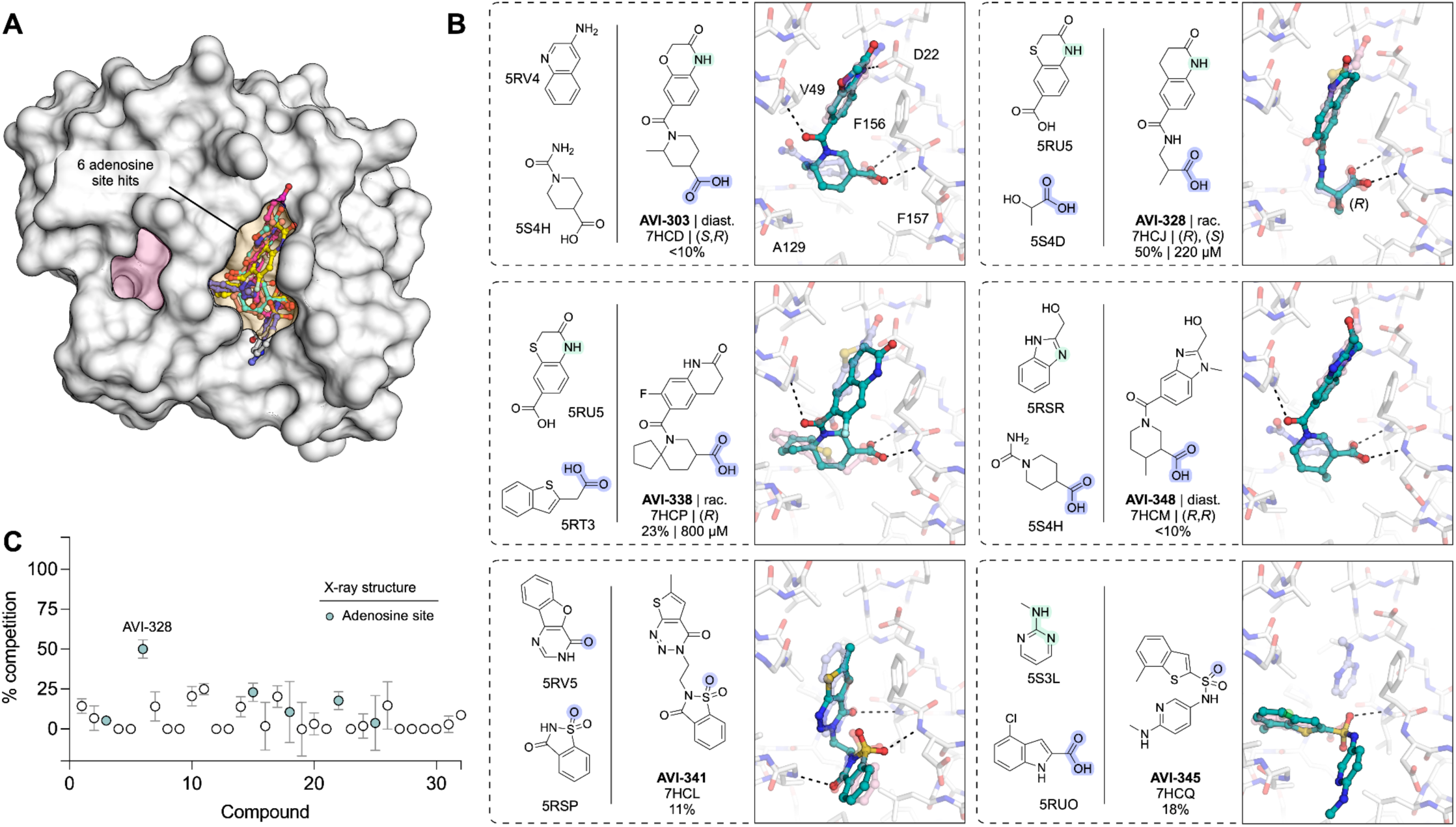
Structural and biophysical characterization of compounds identified by TS-FrankenROCS. (**A**) Surface view of Mac1 showing the binding site of the six TS-FrankenROCS hits. (**B**) Chemical structures of parent fragments and ligands identified by TS-FrankenROCS. The X-ray crystal structure of the ligand bound to Mac1 (teal sticks) is overlaid with the structures of the parent fragments (transparent blue/pink sticks).

Hydrogen bonds between the TS-FrankenROCS ligand and Mac1 are shown as dashed-black lines. Results for the preliminary HTRF screen (% competition at 200 μM, mean of three technical replicates) and the confirmatory dose-response screen (IC_50_, best fit value of three technical replicates) are indicated. (**C**) Plot showing HTRF assay results for the 32 compounds synthesized from the FrankenROCS pipeline.

Although the TS-FrankenROCS approach did not identify ligands with improved binding affinity compared to the original FrankenROCS approach, the Tanimoto Combo scores were significantly higher when searching the REAL database compared to the HTS library (**Fig. S5**). Ultimately, we discovered six new adenosine site-binding ligands, including two sulfonamides that bound to the oxyanion subsite (**Fig. 4B**).The Mac1-AVI-345 X-ray crystal structure shows the potential for a sulfonamide to both contact the oxyanion site and serve as a linker for targeting the phosphate tunnel, for which the original screen identified very few fragments (*1*).

### Extensive exploration of the chemical space around AVI-313

While our previous fragment merging approach yielded sub micromolar compounds carboxylates (*2*), the poor permeability and modest ligand efficiency of these compounds motivated a search for improved leads. From the FrankenROCS searches of the Enamine HTS collection and the REAL database, we observed much lower hit rates for non-carboxylic acid containing matches. Despite this low overall hit rate, the most promising neutral FrankenROCS hit was AVI-313, which contains deazapurine head group bound in the adenine subsite and a 2-aminoethanol core that formed a hydrogen bond to the carbonyl of Leu126 (**Fig. 5A,B**). Other features of AVI-313 that make it an appealing scaffold include its ligand efficiency, accessible vectors for growth, and ease of analog synthesis from amine building blocks.

**Figure 5.**
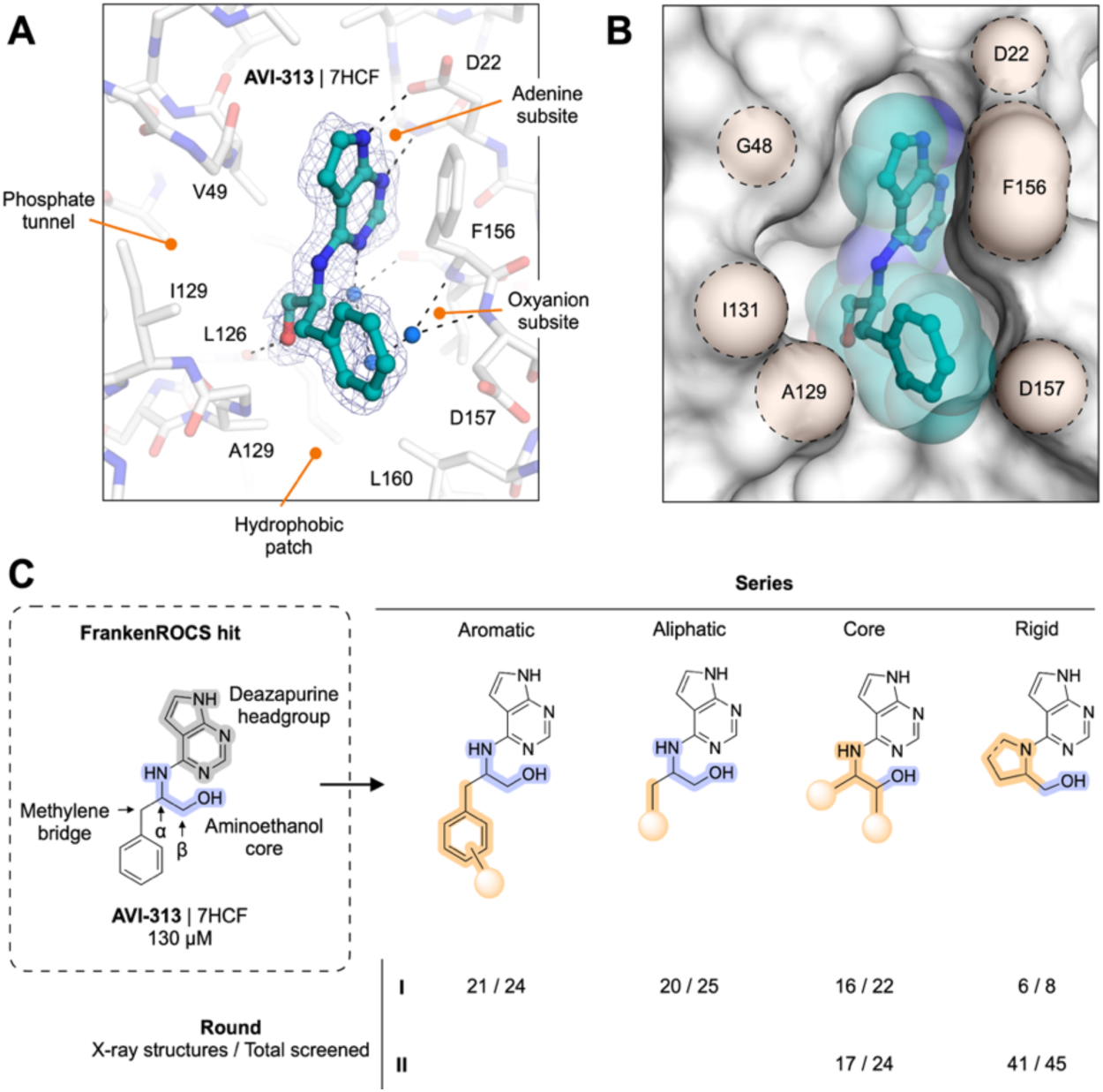
Structure-based optimization of AVI-313. (**A**) Structure of AVI-313 (teal sticks) bound in Mac1 active site (white sticks). Selected ligand-protein hydrogen bonds are shown with black-dashed lines. Water molecules that mediate bridging protein-ligand hydrogen bonds are shown with blue spheres. The PanDDA event map is contoured around the ligand (blue mesh, 2 σ). (**B**) Alternative view of AVI-313 (teal sticks/spheres) bound in Mac1 active site (white surface). Only the major *S* isomer is shown. Important active site residues are shaded yellow. (**C**) Summary of AVI-313 optimization into distinct sub-series with 121 X-ray structures.

To create a more potent inhibitor of Mac1, we aimed to introduce new polar interactions with the oxyanion subsite and phosphate tunnel, while optimizing contacts with the hydrophobic patch that sits below the phenyl ring in the AVI-313 structure (**Fig. 5A,B**). Analogs were synthesized in four series, either by substituting the benzyl group of AVI-313 for other aromatic groups (**aromatic series**), replacing the benzyl group with aliphatic groups (**aliphatic series**), removing the benzyl group and decorating the 2-aminoethanol portion of AVI-313 (**core series**) or cyclizing the 2-aminoethanol nitrogen (**rigid series**) (**Fig. 5C**, **Table S2**B). Compounds were selected from the Enamine REAL database using the 2D molecular similarity search engine SmallWorld (http://sw.docking.org) and the substructure browser Arthor (http://arthor.docking.org) (*4*) or were synthesized in house. To iterate on results from the first round of analogs, we synthesized a second round that focused on the core and rigid series (**Fig. 5C**). In total, we synthesized and screened 148 compounds and determined 121 crystal structures. Compounds were screened for Mac1 binding at 200 μM using the HTRF assay and by soaking into P4_3_ crystals. Because of the large number of compounds with >40% binding at 200 μM, we typically determined IC_50_ values for compounds with >80% binding (**Fig. S3**).

The resulting dataset is notable for the number of high-resolution X-ray crystal structures determined (121 of the 148 compounds screened). We soaked all compounds at 10 or 20 mM into Mac1 crystals and detected binding events using the PanDDA algorithm, which enhances ligand-bound densities via a real space subtraction of the unbound state density (*13*). While the standard PanDDA procedure of using DMSO-only datasets to model the unbound state was effective for the majority of our datasets, this procedure yielded density that was not immediately interpretable for others. We recognized that these datasets were obtained from compounds that were purified as trifluoroacetic acid (TFA) salts. Modeling these compounds, even with the knowledge of the TFA binding pose based on a previously determined structure (*1*), was often difficult, because simultaneous binding of TFA to the oxyanion subsite meant that electron density maps featured both the ligand and TFA. To resolve the ligands, we created a background electron density map for PanDDA using 44 datasets collected from crystals soaked with 10 mM TFA and 10% DMSO rather than the conventional DMSO control datasets (**Fig. S7**). Removing the TFA signal using this background enhanced the interpretability of the PanDDA event maps, and allowed high confidence modeling of 53 ligands where binding events were detected in the presence of TFA (**Table S3**D). Collectively, the affinity and structural data allowed us to parse several SAR trends and improve the potency of this series to single digit micromolar affinities.

#### Aromatic series explore hydrophobic packing in the southern pocket

We tested 24 analogs of AVI-313 where substituents were added to the phenyl ring or where the phenyl ring was replaced with other aromatic groups (**Fig. 6**). Analogs either retained (**Fig. 6A**) or modified (**Fig. 6B**) the methylene bridge adjacent to the phenyl ring of AVI-313. For analogs that retained the methylene bridge, we observed substituent-dependent occupancy of either the oxyanion subsite or the phosphate tunnel by the phenyl substituent (**Fig. 6C-D**). Because the 2-aminoethanol core was present in all molecules tested, this subsite switching is caused by changes to the aromatic group. Analogs with *para*-substituents, and all five-membered rings, matched the AVI-313 pose, with the 2-aminoethanol hydroxyl forming a hydrogen bond to the backbone carbonyl of Leu126 (**Fig. 6C**). The hydroxyl of compounds with an *ortho-* or *meta*-substituted phenyl group were oriented towards the oxyanion subsite, matching the parent fragment of AVI-313 (**Fig. 2B**, **Fig. 6D**). Although several compounds contained aromatic heteroatoms with the potential to act as hydrogen bond acceptors (e.g. AVI-3705, AVI-3703, AVI-3673, AVI-3657), none formed hydrogen bonds with the backbone nitrogen of Ala129 (**Fig. 6C**). This suggests that the methylene bridge needs to be extended to engage Ala129. On the other side of the adenosine site entrance, the *para*-hydroxyl of AVI-4066 formed a hydrogen bond to the sidechain of Asp157 and led to a 60° rotation around the Phe156 χ1 dihedral angle (**Fig. 6F**). Despite structural and binding diversity, and the formation of a new polar contact, none of the analogs that retained the methylene bridge improved binding affinity measured by the HTRF assay compared to the AVI-313 parent (**Fig. 6A**).

**Figure 6.**
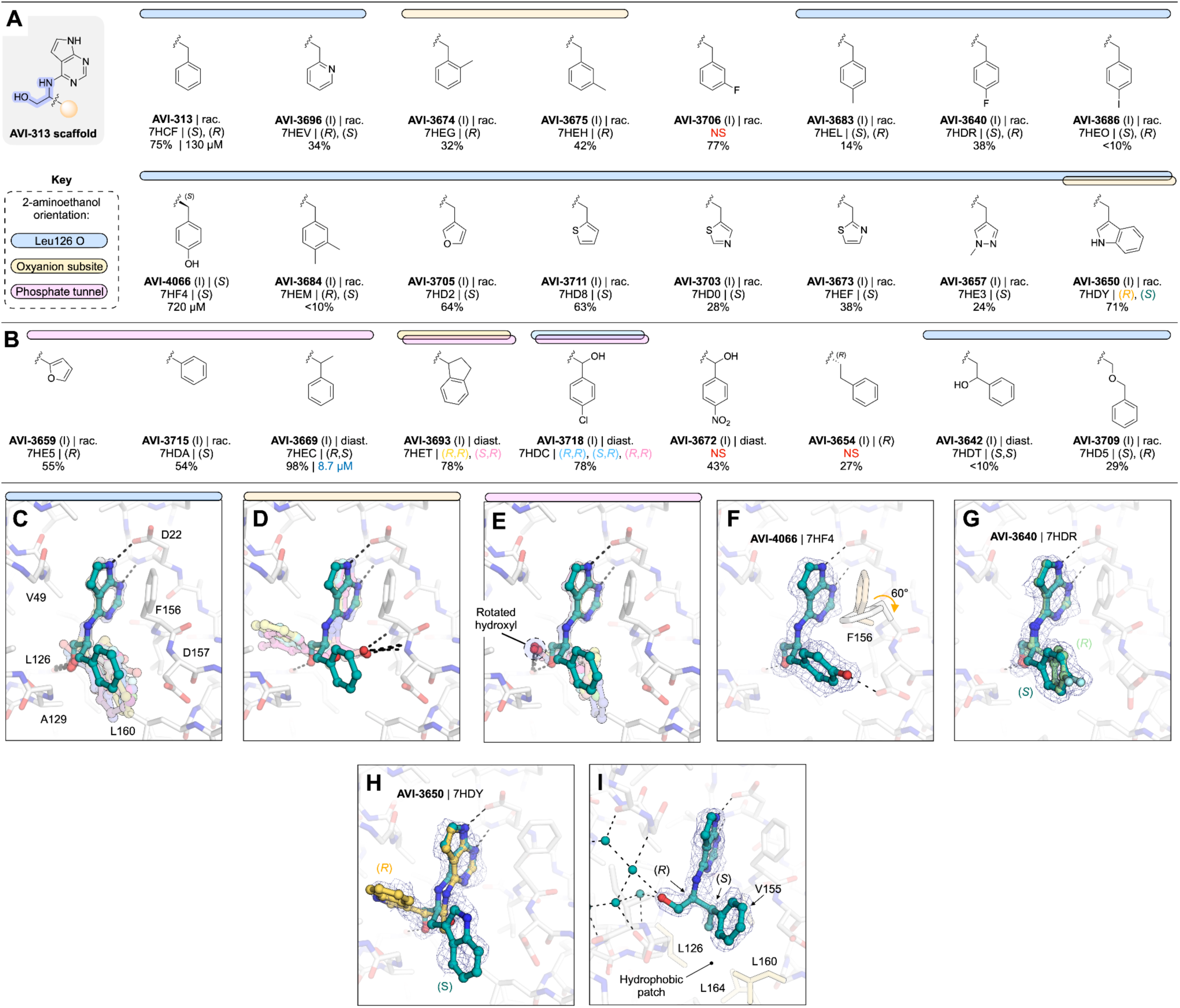
Aromatic series analogs of AVI-313. (**A**) Chemical structures of AVI-313 analogs with an intact methylene bridge. Results for the preliminary HTRF screen (% competition at 200 μM, mean of three technical replicates) and the confirmatory dose-response screen (IC_50_, best fit value of three technical replicates) are indicated. Compounds were synthesized as pure isomers, as racamates (rac.) or as mixtures of diastereomers (diast). (**B**) Chemical structures AVI-313 analogs where the methylene bridge was modified. (**C**,**D**,**E**) Alignment of the X-ray crystal structures of compounds from the aliphatic series where the 2-aminoethanol hydroxyl formed a hydrogen bond with the oxyanion subsite (**C**), the backbone carbonyl of Leu126 (**D**) or where the hydroxyl is rotated toward the phosphate tunnel (**E**). (**F**-**I**) X-ray crystal structures of AVI-4066 (**F**), AVI-3640 (**G**), AVI-3650 (**H**) and AVI-3669 (**I**) bound in the Mac1 active site. PanDDA event maps are shown contoured around the ligands (blue mesh, 2 σ).

For compounds synthesized as racemates that contained a *para* phenyl substituent, or 2-pyridine, we observed electron density consistent with both *R* and *S* isomers, similar to the AVI-313 parent (**Fig. 6G**, **Fig. S2**). Both isomers of the 3-indole containing analog were bound (AVI-3650), however, the aromatic group of the *R* isomer bound in the phosphate tunnel while the *S* isomer matched the AVI-313 parent (**Fig. 6H**, **Fig. S2**). Likewise, analogs with *ortho-* and *meta*-substituted phenyl groups bound in the phosphate tunnel were also the *R* isomers (AVI-3674, AVI-3675). Therefore, the preference of AVI-313 analogs for either subsite is linked to both functional group identity and stereochemistry.

Next, we tested nine AVI-313 analogs where the methylene bridge was modified, either by adding functional groups or changing the length (**Fig. 6B**). In the X-ray crystal structures, the aromatic groups were generally oriented parallel to the oxyanion subsite, similar to the AVI-313 parent. The exception was the indane of AVI- 3693, where two isomers were bound, one matching the AVI-313 parent and the other with the indane portion oriented toward the phosphate tunnel (**Fig. S2**). For the analogs that matched the position of the AVI-313 parent, the hydroxyl typically formed a hydrogen bond to the backbone carbonyl of Leu126. However, the hydroxyl was rotated to an alternative position for several analogs where the methyl bridge was modified (**Fig. 6E**). Despite the rotated hydroxyl, AVI-3669 was >10-fold more potent (IC_50_ = 8.7 μM) than the AVI-313 parent. The electron density in the Mac1-AVI-3669 structure showed that only the (*R,S*)-diastereomer was bound (**Fig. 6I**), with the phenyl ring matching the *R* isomer of AVI-313. Because the compound was tested as a mixture of four diastereomers, the true IC_50_ could be up to four-fold more potent (∼2 μM).

Collectively, the aromatic series revealed binding pose diversity in AVI-313 analogs. Preference for binding of the aromatic group to either the phosphate tunnel or oxyanion subsite was linked to functional group identity and stereochemistry. The diversity of binding suggests that the hydrogen bond of the 2-aminoethanol core to the backbone carbonyl of Leu126 is relatively weak. This is supported by the absence of this hydrogen bond in the most potent compound from the series (AVI-3669) (**Fig. 6I**). Instead of hydrogen bonding to Leu126 or the oxyanion subsite, the hydroxyl formed water-mediated contacts with the phosphate tunnel (**Fig. 6I**). The structural basis for the improved binding of this analog appears to be the hydrophobic contacts made between the methyl group and the hydrophobic patch containing residues Leu126, Val155 and Leu160 (**Fig. 6I**). This link between hydrophobic contact and high affinity binding in the absence of polar contacts uncovers a new driver of potency. At the same time, we considered that the relatively large aromatic/heteroaromatic side chains might limit the repertoire of distinct binding orientations and might explain the inability of this series in general to form polar contacts with the oxyanion site.

#### Aliphatic series reveals phosphate tunnel and hydrophobic patch interactions

In parallel, we tested a series of AVI-313 analogs in which aliphatic groups were joined to the 2-aminoethanol core directly, or via a short alkylene spacer (**Fig. 7A,B**). We tested 25 analogs in total and determined 20 crystal structures (**Fig. S2**). As with the aromatic series, we observed functional group-dependent preference for either the oxyanion subsite or phosphate tunnel (**Fig. 7C,D**). For seven of the compounds, the 2- aminoethanol hydroxyl was rotated away from the Leu126 carbonyl (**Fig. 7E**). The hydroxyethyl-substituted diol AVI-3734 was among the most potent of these analogs in the HTRF assay (IC_50_ = 52 μM). The 2-aminoethanol hydroxyl in AVI-3734 formed a hydrogen bond with the oxyanion subsite, while the hydroxyethyl hydroxyl displaced a tightly-bound water molecule that formed a bridging interaction with ADP-ribose and also a large number of Mac1-binding fragments (*1*, *17*) (**Fig. 7F**). We tested two tertiary amines in this series (AVI-3671 and AVI-3677); while both formed hydrogen bonds to the oxyanion subsite via bridging water molecules, neither had detectable binding at 200 μM in the HTRF assay (e.g. AVI-3671 shown in **Fig. 7G**, **Fig. S2**). In contrast, replacing the benzyl group of AVI-313 with isopropyl increased the binding affinity ∼10-fold (AVI- 1504, IC_50_ = 9.8 μM). Notably, with AVI-1504, the 2-aminoethanol formed two hydrogen bonds with the oxyanion sub-site (2.9 and 3.1 Å), while the isopropyl group faces the phosphate tunnel (**Fig. 7H**). AVI-1504 contained the same 2-amino-3-methylbutan-1-ol core as AVI-3669, the most potent compound from the aromatic series (**Fig. 6I**). Although both compounds placed methyl groups in the hydrophobic patch, the stereochemistry at the α-carbon was inverted and there was a 1.3 Å difference in the methyl group that occupied the hydrophobic patch as well as a slight tilt in the deazapurine head group (**Fig. 7I**). Despite this tilt, both compounds formed short hydrogen bonds to the adenine subsite (2.6-2.8 Å) (**Fig. 7I**). We also tested several cyclic alkanes and ethers branching at the methylene carbon (**Fig. 7B**), however none improved binding affinity in the HTRF assay compared to the AVI-313 parent and all were much weaker binders than AVI-1504.

**Figure 7.**
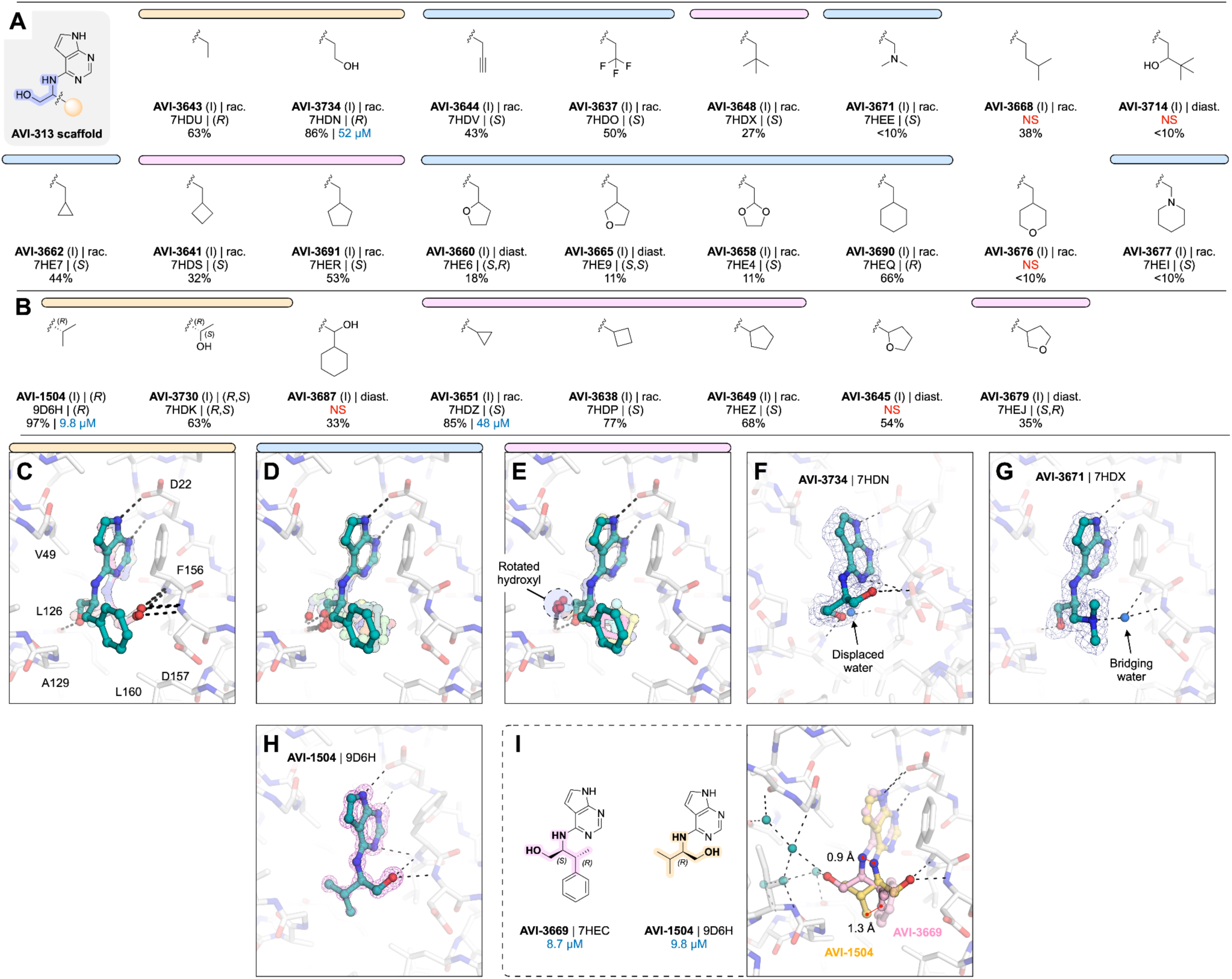
Aliphatic series analogs of AVI-313. (**A**) Chemical structures AVI-313 analogues with an intact methylene bridge. Results for the preliminary HTRF screen (% competition at 200 μM, mean of three technical replicates) and the confirmatory dose-response screen (IC_50_, best fit value of 3-4 technical replicates) are indicated. (**B**) Chemical structures AVI-313 analogs with groups added to the methylene bridge. (**C**,**D**,**E**) Alignment of the X-ray crystal structures of compounds from the aliphatic series where the 2-aminoethanol hydroxyl formed a hydrogen bond with the oxyanion subsite (**C**), the backbone carbonyl of Leu126 (**D**) or where the hydroxyl is rotated toward the phosphate tunnel (**E**). (**F**-**H**) X-ray crystal structures of AVI-3734 (**F**), AVI-3671 (**G**) and AVI-1504 (**H**) bound in the Mac1 active site. PanDDA event maps are shown contoured around the ligands (blue mesh, 2 σ), except for AVI-1504, where the F_O_-F_C_ difference map is shown prior to ligand placement (purple mesh, 5 σ). (**I**) Structural comparison of the most potent AVI-313 analogs emerging from the aromatic and aliphatic series. The distance between the methyl groups that occupy the hydrophobic patch is indicated.

Collectively, analogs tested in the aliphatic series illustrate the flexibility of the 2-aminoethanol core in the Mac1 adenosine site. Modifying the group at the α-carbon switches the preference of hydrogen bonding from the phosphate tunnel (Leu126 backbone oxygen) to the adenine subsite (Leu126 backbone N-H, Ala154 backbone oxygen) or the oxyanion subsite (Phe156 backbone N-H, Asp157 backbone N-H). Although flexibility allows the adenosine site to accommodate larger aliphatic groups, the most potent compound from this series (AVI-1504, IC_50_ = 9.8 μM) was among the smallest tested, and thus the most ligand efficient (**Fig. 7H**). Identification of AVI-1504 as a potent Mac1 binder illustrates the importance of both polar and non-polar contacts for optimal binding to the adenosine site. Whereas the most potent compound from the aromatic series also placed a methyl group in the hydrophobic patch (AVI-3669, IC_50_ = 8.7 μM), the phenyl ring contributes little to the binding affinity and ligand efficiency is reduced. Alignment of the AVI-1504- and AVI-3669-Mac1 X-ray crystal structures shows that despite chemical similarity, the compounds bind with the hydroxyl vector pointing toward the phosphate tunnel (AVI-3669) or oxyanion subsite (AVI-1504) (**Fig. 7I**).

#### Core series highlights the contributions of protein flexibility and water displacement

Inspired by the potent and ligand efficient binding of AVI-1504, we searched for alternative scaffolds by modifying the 2-aminoethanol core (**Fig. 8**). We initially tested 21 analogs with groups at either (or both) of the α- and β-carbons (**Fig. 8A**). Similar to analogs in the aromatic and aliphatic series, we observed mobility of the 2-aminoethanol core between the oxyanion subsite and the phosphate tunnel (**Fig. 8** B-D). Although none of the compounds improved binding affinity relative to AVI-1504, eight of the analogs had IC_50_ values below 50 μM (**Fig. 8A**). The most potent was AVI-3761 (IC_50_ = 13 μM), where a cyclohexyl ring containing the β-carbon occupied the hydrophobic patch (**Fig. 8E**). This contact was similar to the low micromolar compounds identified from the aromatic and aliphatic series (AVI-3669 and AVI-1504), although the larger size of the cyclohexyl group led to rotation of the Leu160 side-chain away from the ligand (**Fig. 8E**). Unlike AVI-1504, which formed two hydrogen bonds to the oxyanion subsite, the hydroxyl of AVI-3761 formed a single hydrogen bond to the backbone nitrogen of Phe156 (3.0 Å), along with an intramolecular hydrogen bond to the pyrimidine nitrogen (2.6 Å) (**Fig. 8E**). The importance of hydrophobic contacts at this subsite is supported by the complete loss of binding when cyclobutane at the α-carbon was replaced by 3-oxetane (**Fig. 8A**). In addition, the similar analog bearing a gem-dimethyl at the β-carbon was relatively potent (AVI-3758, IC_50_ = 28 μM), despite the 2- aminoethanol hydroxyl not forming a hydrogen bond with the oxyanion subsite (**Fig. 8F**). In the Mac1-AVI-3758 X-ray crystal structure, two alternative conformations of the ligand were modeled (**Fig. 8F**). In both conformations, the methyl groups occupied the hydrophobic patch, while the Asp157 backbone carbonyl is rotated towards the adenosine site. The same conformational change was observed for cyclobutyl and cyclopentyl analogs, as well as several other analogs in the core series (**Fig. S2**). We also tested 10 analogs where groups were added to both α and β carbons of the 2-aminoethanol core, or where the α and β carbon were fused into a ring (**Fig. 8A**). None of these analogs had IC_50_ values below 50 μM.

**Figure 8.**
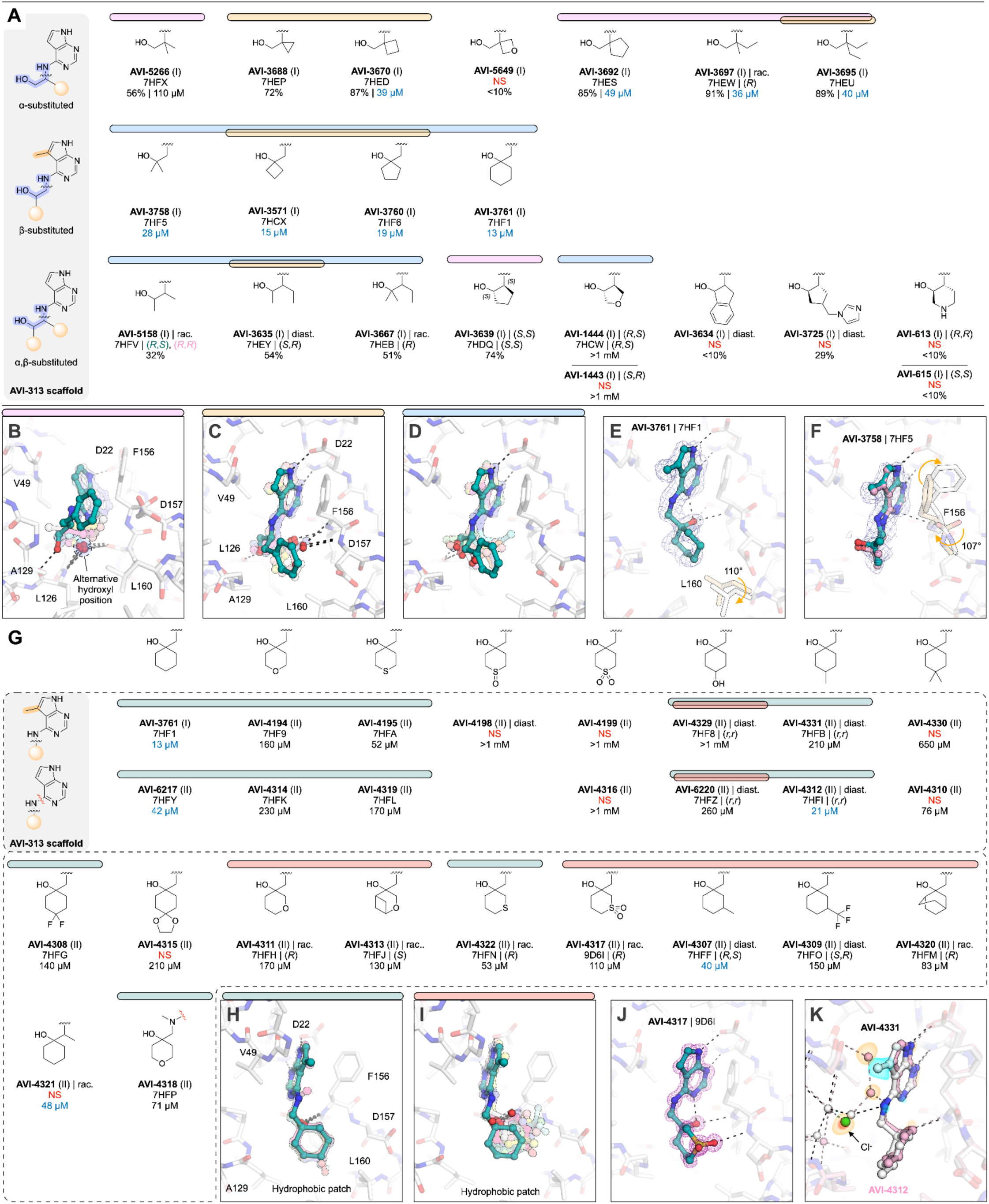
Core series analogs of AVI-313. (**A**) Chemical structures of AVI-313 analogs where the ethylene bridge was substituted at the α, β or both α and β positions. Results for the preliminary HTRF screen (% competition at 200 μM, mean of three technical replicates) and the confirmatory dose-response screen (IC_50_, best fit value of 3-4 technical replicates) are indicated. (**B**,**C**,**D**) Alignment of the X-ray crystal structures of the AVI-313 analogs where the 2-aminoethanol hydroxyl formed a hydrogen bond with either the backbone nitrogen of Leu126 (**B**), the oxyanion subsite (**C**) or where the hydroxyl was rotated towards the phosphate tunnel (**D**). (**E**,**F**) X-ray crystal structures of AVI-3761 (**E**) and AVI-3758 (**F**) bound in the Mac1 active site. PanDDA event maps are shown contoured around the ligands (blue mesh, 2 σ). (**G**) Chemical structures of AVI-3761 subseries. Several analogs (top row) were synthesized with a methyl group on the 5 position of the 7H-pyrrolo[2,3-d]pyrimidine. (**H**, **I**) Alignment of the X-ray crystal structures of AVI-3761 analogs where the six- membered ring occupied a conformation similar to the AVI-3761 parent (**H**) or where the six-membered ring was rotated away from the hydrophobic patch (**I**). (**J**) X-ray crystal of AVI-4317 bound in the Mac1 active site. The F_O_-F_C_ difference electron density map prior to ligand placement is contoured around the ligand (purple mesh, 5 σ). (**K**) Alignment of the X-ray crystal structures of AVI-4331 (white sticks/spheres) and AVI-4312 (pink sticks/spheres) bound to Mac1. F_O_AVI-4331_-F_O_AVI-4312_ isomorphous difference map contoured within 5 Å of the ligand at +5 σ (blue surface) and -5 σ (orange surface). The chloride ion and nearby water network are perturbed in the AVI-4331 crystal structure relative to the AVI-4312 crystal structure.

To optimize the contact between the AVI-3761 and the hydrophobic patch, we designed and synthesized a set of 25 analogs with additional modifications to the cyclohexyl ring (**Fig. 8G-I**). A subset was synthesized with a methyl at the 5-position of the deazapurine headgroup, matching the AVI-3761 parent. Analogs that contained a 3- or 4-methyl group on the cyclohexyl ring were of similar potency to the AVI-3761 parent (AVI-4312, AVI- 4307, **Fig. 8G**). Two conformations of the 4-hydroxyl-containing analogs were observed in the X-ray crystal structures (**Fig. S2**). Although the 2-aminoethanol hydroxyl formed a hydrogen bond with the oxyanion subsite in both conformations, the cyclohexyl ring was rotated out of the hydrophobic patch in one of the conformations (**Fig. 8I**). Similar rotation of the cyclohexyl ring was observed for six other analogs (**Fig. 8G, I**). Addition of a sulfonyl to the 3-position in AVI-4317 created a new hydrogen bond to the backbone nitrogen of Asp157 (**Fig. 8J**), however, this analog was ∼4-fold less potent compared to the AVI-3761 parent (IC_50_ = 110 μM). Despite six analogs retaining the AVI-3761-like pose, none improved binding compared to the AVI-3761 parent (**Fig. 8G**). Differences between binding of the methyl- and des-methyl analogs could be linked to ligand- induced changes in the phosphate tunnel water network (**Fig. 8K**). Taken together, these results suggest that the cyclohexyl group of AVI-3761 is well suited for contact with the hydrophobic patch.

#### Rigidification identifies ligand-induced conformational changes in the adenosine site

Next, we constrained the 2-aminoethanol core into a series of more rigid, cyclic pyrrolidine- and piperidine-containing analogs (**Fig. 9A-D**). Initial screening of eight compounds identified the pyrrolidine AVI-607 as a low micromolar binder (IC_50_ = 5.1 μM) (**Fig. 9A**). The Mac1-AVI-607 X-ray crystal structure showed that the hydroxyl formed a single hydrogen bond with the backbone nitrogen of Phe156 (3 Å), along with an intramolecular hydrogen bond to the pyrimidine nitrogen (2.6 Å) and a hydrogen bond to the buried water in the adenine subsite (2.8 Å) (**Fig. 9E**). These features make the binding of AVI-607 similar to AVI-3761 (**Fig. 8E**). In addition, AVI-607 positioned a methyl group in the hydrophobic patch, matching the low micromolar affinities of compounds from the aromatic and aliphatic series (AVI-3669, AVI-1504). The Asp157 backbone carbonyl was also rotated in a similar manner to AVI-3758 (**Fig. 8F**). To explore the AVI-607 scaffold, we tested seven analogs that modified the gem-dimethyl motif (**Fig. 9A**). Removing one methyl group led to a ∼3-fold decrease in affinity (AVI-4062, IC_50_ = 55 μM). Although the ligand pose was identical to AVI-607, the Asp157 carbonyl retained the apo-like conformation. This indicates that the carbonyl flip is triggered by non-polar contact with the ligand. Removing both methyl groups from AVI-607 led to a >20-fold decrease in affinity (AVI-3685, 48% inhibition at 200 μM) and the hydroxyl was now oriented towards the phosphate tunnel (**Fig. S2**). This pose was similar to the *S*-isomer of AVI-607, which was ∼5-fold less potent compared to AVI-607 (AVI-1445, IC_50_ = 110 μM). Furthermore, replacing the single methyl in AVI-4062 with ethyl led to a ∼4-fold decrease in binding affinity (AVI-3666, IC_50_ = 46 μM). This demonstrates that for this scaffold, methyl best occupies the hydrophobic patch. Next, we tested AVI-607 analogs where the gem dimethyl was replaced by hydroxyethyl, phenyl or indane groups (**Fig. 9A**). While none of these modifications substantially improved binding, the indane analog was equipotent to AVI-607 (AVI-3731, IC_50_ = 3.7 μM). The X-ray crystal structure showed that AVI-3731 caused a large-scale conformational change in the Phe132 loop, similar to a conformation observed previously and serendipitously in the presence of compounds obtained from docking (*2*) (**Fig. S2**, **Fig. S7**).

**Figure 9.**
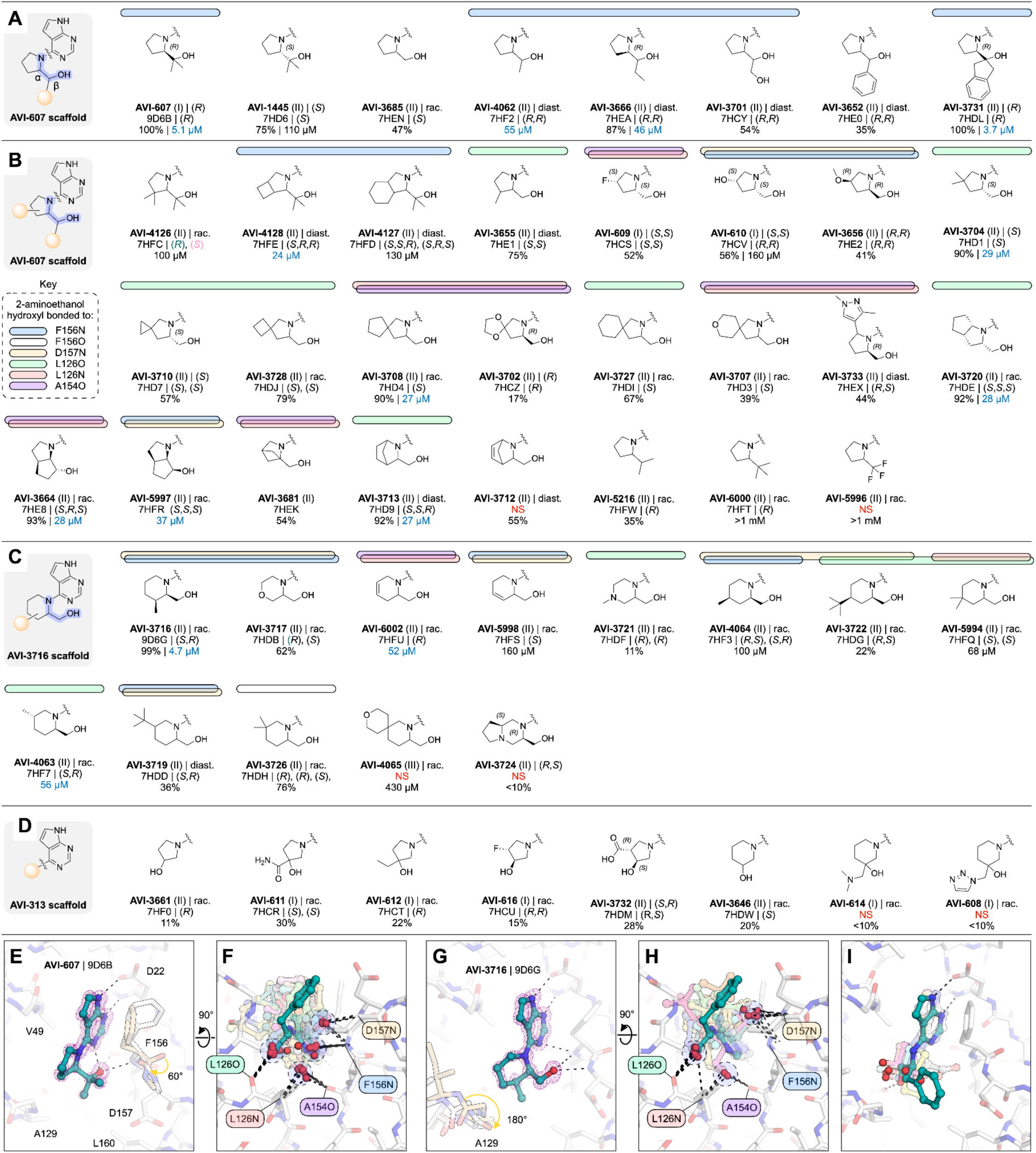
Rigid series analogous of AVI-313. (**A**,**B**,**C**,**D**) Chemical structures of pyrrolidine analogs with groups at the β-position of the 2-aminothanol core (**A**), analogs with groups on the pyrrolidine ring (**B**), piperidine analogs (**C**) or pyrrolidine/piperidine cyclised at the β carbon. (**D**). Results for the preliminary HTRF screen (% competition at 200 μM, mean of three technical replicates) and the confirmatory dose-response screen (IC_50_, best fit value of 3-4 technical replicates) are indicated. (**E**) X-ray crystal structure of AVI-607 bound in the Mac1 active site. The F_O_-F_C_ difference electron density map prior to ligand placement is contoured around the ligand (purple mesh, 5 σ). (**F**) Alignment of the 31 X-ray crystal structures of pyrrolidines from the rigid series. Hydrogen bonds are indicated with dashed black lines. (**G**) X-ray crystal structure of AVI-3716 bound in the Mac1 active site. The F_O_-F_C_ difference electron density map prior to ligand placement is contoured around the ligand (purple mesh, 5 σ). (**H**) Alignment of the 13 X-ray crystal structures of piperidines from the core series. (**I**) Alignment of the six X-ray crystal structures of analogs containing a pyrrolidine or piperidine cyclized at the β-carbon.

We next explored AVI-607 analogs bearing pyrrolidine ring substitutions, including additional fused, bridged, or spirocyclic ring systems (**Fig. 9B**), as well as ring-expansion of pyrrolidine to six-membered heterocycles (**Fig. 9C**) and alternative cyclic amines (**Fig. 9D**). Because few of the relevant building blocks were available with gem-dimethyl substitution, the majority of pyrrolidine analogs were tested without this motif (**Fig. 9B**).

Alignment of all the pyrrolidine analogs shows the diversity of hydrogen bonding within the adenosine site, despite the common, cyclized 2-aminoethanol core (**Fig. 9F**). Three of the AVI-607 analogs with groups at the 3- or 4-positions of the pyrrolidine ring stabilized the open conformation of the Phe132 loop (**Fig. S2**, **Fig. S7**). The most potent analog contained a cyclobutane ring fused to the pyrrolidine (AVI-4128, IC_50_ = 24 μM) and had a similar binding pose to AVI-607 (**Fig. S2**). Although none of the other pyrrolidine analogs improved binding affinity relative to AVI-607, seven had IC_50_ values below 50 μM, five of which had inverted stereochemistry at the α-carbon relative to the AVI-607 parent (**Fig. 9B**).

In parallel, we screened a series of analogs that contained the 2-aminoethanol core in cyclic piperidines (**Fig. 9C**). The most potent was AVI-3716, with an IC_50_ equal to 4.5 μM. The X-ray crystal structure showed that the AVI-3716 hydroxyl formed a short hydrogen bond with the backbone nitrogen Asp157 (2.9 Å) and a longer (3.3 Å) hydrogen bond with the Phe156 backbone nitrogen, while the piperidine methyl group occupied the hydrophobic patch (**Fig. 9G**). The binding pose of AVI-3716 closely matches the binding of AVI-1504 (RMSD = 0.21 Å for matching atoms). There was a flip in the Ala129-Gly130 peptide bond in the AVI-3716 structure, similar to the conformational change seen in X-ray crystal structure of Mac1 in complex with ADP-ribose (*1*, *17*). Alignment of the X-ray crystal structures of the 12 additional piperidine analogs showed that the 2-aminoethanol core switched between the phosphate tunnel and oxyanion subsite (**Fig. 9H**). None of the piperidine analogs, or eight cyclic analogs containing a pyrrolidine or piperidine cyclized at the β-carbon, improved binding relative to AVI-3716 (**Fig. 9C,D**,I).

Collectively, analogs with pyrrolidine and other rigidification reinforce the importance of hydrogen bonding to the oxyanion subsite and non-polar contacts in the hydrophobic patch for potent binding. Despite chemical similarity, the diverse crystallographic binding poses of the most potent ligands (e.g. AVI-607, AVI-3731 and AVI-3716) suggests that these compounds offer distinct starting points for lead optimization. In addition, the 49 X-ray crystal structures, including structures for many weak binders, identified ligand-induced conformational changes in the adenosine site.

### Extensive analoging reveals affinity determinants in the oxyanion subsite and hydrophobic patch

The extensive testing of AVI-313 analogs allowed us to uncover two features of 2-aminoethanol containing ligands that were strongly associated with potent Mac1 binding. Firstly, four out of five most potent compounds formed hydrogen bonds to the oxyanion subsite with the 2-aminoethanol hydroxyl. For AVI-607 and AVI-3761, the hydroxyl formed a hydrogen bond with the Phe156 backbone N–H, as well as an intramolecular hydrogen bond with the N3 pyrimidine nitrogen of the deazapurine head group and a hydrogen bond to the tightly bound water molecule in the adenine subsite (**Fig. 9E**, **Fig. 8E**). In contrast, the 2-aminoethanol hydroxyl of AVI-1504 and AVI-3716 formed a hydrogen bond to the backbone N–H of Asp157 (**Fig. 7H**, **Fig. 9G**). Surprisingly, no direct hydrogen bond was observed to the AVI-3669 hydroxyl. Instead, the hydroxyl formed water-mediated hydrogen bonds with the phosphate tunnel (**Fig. 6I**).

The second feature of the low micromolar scaffolds reported here is the placement of methyl/alkyl groups in the hydrophobic patch of the adenosine site, exemplified by AVI-3669 (**Fig. 10A,B**). Removing groups that interact with the hydrophobic patch (e.g., Me in AVI-3669) led to a 5-20-fold decrease in binding affinity (c.f. des-Me AVI-313 versus AVI-3669). Contact with the hydrophobic patch was also a feature of the previously reported pyrrolidinone inhibitor of Mac1 (*2*) (**Fig. 10A**). Furthermore, the X-ray crystal structure of the most potent compound from TS-FrankenROCS (AVI-328) showed that a methyl group makes contact with the hydrophobic patch (**Fig. 10A**). Although this molecule binds weakly (IC_50_ = 220 μM), substitution of the 3,4- dihydroquinolinone with an alternative headgroup that forms hydrogen bonds to the adenine subsite might improve affinity. Going forward, the affinity of compounds that interact with the oxyanion subsite could be rapidly improved by the addition of nearby methyl groups. It’s possible that alternative hydrophobic groups (e.g. chloro) might provide a better fit to the pocket. In summary, exhaustive testing of AVI-313 analogs reveal the structural determinants of high affinity binding for deazapurine derived ligands containing a 2-aminoethanol core.

**Figure 10.**
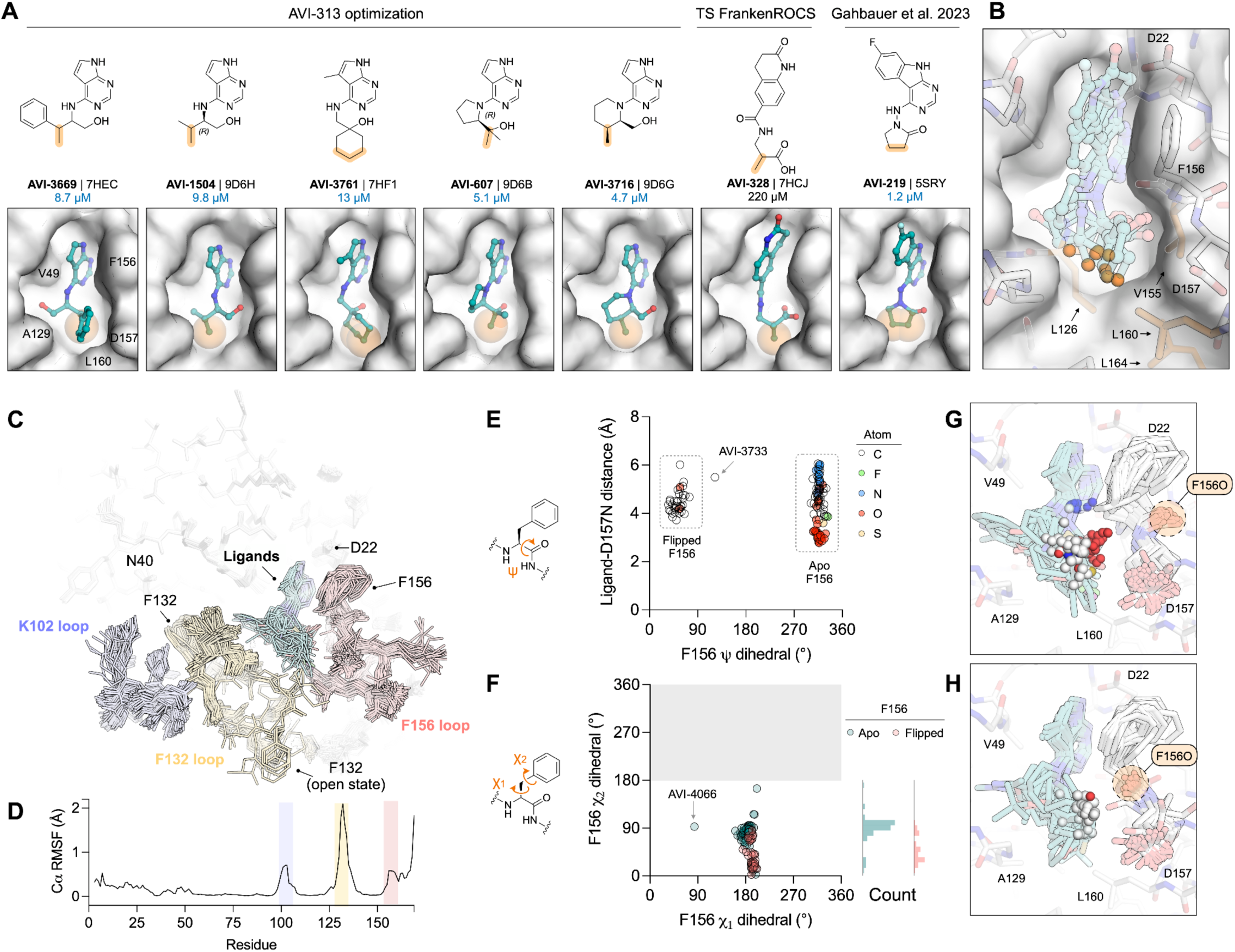
Hydrophobic contacts and protein flexibility underlie potent binding of AVI-313 analogs.(**A**) Chemical structures (top) and surface view (bottom) of selected ligands that occupy the hydrophobic pocket. Carbon atoms that make contact with the hydrophobic patch are shown with transparent orange spheres. (**B**) Alignment of the seven X-ray crystal structures for compounds shown in (**A**). Carbon atoms that contact the hydrophobic patch are shown with transparent orange spheres. The apo Mac1 X-ray crystal structure is shown with a white surface (PDB 7KQO). (**C**) Alignment of chain A of the 121 X-ray crystal structures of AVI-313 analogs. Protein residues are shown with sticks colored by loop and the ligand is shown with teal sticks. The position of the Phe132 side-chain in the open-state structures is indicated (see **Fig. S7**). (**D**) Cα root-mean-square fluctuation (RMSF) calculated for chain A across the 121 AVI-313 X-ray crystal structures. (**E**) Plot of backbone Phe156 ψ dihedral angle for all AVI-313 analog structures plotted as a function of the distance from the Asp157 backbone nitrogen to the closest ligand atom. AVI-3733 stabilized the Phe156-Asp157 peptide bond in the cis-conformation (**Fig. S2**). (**F**) Plot of side-chain Phe156 χ_1_ side chain dihedral angle for AVI-313 analog structures plotted as a function of side-chain χ_2_ dihedral angle. Points are colored by Phe156 backbone ψ dihedral angle. The symmetry of the phenylalanine side-chain means that only χ_2_ angles 0-180° are sampled. (**G**,**H**) Alignment of X-ray crystal structures of all ligands with the Phe156 backbone ψ dihedral angle in the apo state (**G**) or the flipped state (**H**). The ligand atoms closest to the Asp157 backbone nitrogen are shown as solid spheres.

### Multiple scales of protein flexibility revealed across crystallography

Alignment of the X-ray crystal structures of the 121 AVI-313 analog reveals flexibility in the active site loops upon ligand binding (**Fig. 10C,D**). Although flexibility in the Phe156 backbone and side chain have been reported previously (*1*, *2*), here, several structures contained a flip in the Phe156 backbone carbonyl (e.g. AVI- 3758 (**Fig. 8F**), AVI-607 (**Fig. 9E**). Alignment of all the X-ray crystal structures shows that this conformation change is induced by ligands that place non-polar groups near the oxyanion subsite (**Fig. 10E**). Rotation of the Phe156 backbone occurs with a correlated rotation of the side-chain away from the adenosine site (**Fig. 10F**- H). Although rotation of the Phe156 backbone breaks the hydrogen bond to nearby Asp22, this appears to be favorable based on the comparison of AVI-607 (IC_50_ = 5.1 μM) with AVI-4062 (IC_50_ = 55 μM). AVI-4062 binds with a nearly identical pose to AVI-607, but lacks the second methyl group and does not induce the peptide flip. Other examples of small scale flexibility in the adenosine site include a flip of the Ala129 carbonyl in the AVI- 3716-Mac1 structure that mimics the ADP-ribose-Mac1 structure (**Fig. 9G**) and rotation of the Leu160 side-chain to accommodate alkyl groups in the hydrophobic patch (**Fig. 8E**).

In addition to the small-scale flexibility, large-scale changes in the Phe132 loop were observed in the X-ray crystal structures for several ligands, leading to opening of the terminal ribose site (**Fig. S7**), similar to those previously observed, serendipitously, from a compound docked to the canonical conformation (*2*). The most potent open state binder reported here is 10-50-fold more potent compared to those reported previously (AVI- 3731, IC_50_ = 3.7 μM). The higher affinity binding of AVI-3731 is likely due to improved hydrogen bonding to the oxyanion subsite and contact with the hydrophobic patch. Rearrangement of the Phe132 loop in the AVI-3731- Mac1 X-ray crystal structure leads to Phe132 stacking against the ligand, whereas Phe132 is exposed to solvent in other open structures (**Fig. S7**). Collectively, the X-ray crystal structure of 121 AVI-313 analogs capture protein flexibility that underpins the observed diversity in ligand binding pose (e.g. **Fig. 9F,H**). Protein flexibility helps to explain how ligands containing conserved motifs (2-aminoethanol core and deazapurine headgroup) can achieve similar high affinity binding despite adopting diverse binding poses.

### Lead-like AVI-313 analogs have promising solubility, permeability, metabolic stability and selectivity

A major goal of this work was to identify additional chemical matter beyond AVI-92 and AVI-219 (*2*) for our lead optimization efforts against the SARS-CoV-2 macrodomain. Although potent, AVI-92 contains a carboxylic acid and urea, two stereocenters, and five H-bond donors, features that limit cellular permeability and challenge analog synthesis (**Table S3**). In this regard, AVI-219 represents a notable improvement, with increased ligand efficiency realized in a neutral, achiral scaffold. The efforts described herein focused on the AVI-313 core and provide additional lead-like starting points, with good solubility, permeability, and excellent mouse microsomal stability (for example AVI-1504) (**Table S3**). Moreover, their improved ligand efficiency suggests that these AVI-313 analogs comprise a minimal pharmacophore for the adenosine site in Mac1. Promisingly, none of the five most potent scaffolds (AVI-3669, AVI-1504, AVI-3761, AVI-607 or AVI-3716) showed binding to the human macrodomain MacroD2 or Targ1 (**Fig. S3**B,C). Combining features of these various leads in more fully elaborated analogs is ongoing in our groups and will be described in due course. Overall this work demonstrates the value of using scaffold hopping techniques, such as FrankenROCS, to identify alternative starting points for Mac1 medicinal chemistry from fragment hits that contained carboxylic acids as a key binding determinant.

## Discussion

Here, we developed a fragment linking method - FrankenROCS - and used it to discover new inhibitors of the SARS-CoV-2 NSP3 macrodomain using fragments identified from a large-scale X-ray crystallography screen (*1*). Since our fragment screen was performed, several groups have published macrodomain inhibitors (*15*, *18*–*23*), but the major drivers of binding affinity remain relatively unexplored and these compounds have not yet been optimized for *in vivo* tests. Our previous efforts at fragment linking led to AVI-92, a sub-micromolar inhibitor that bound to the oxyanion subsite with a carboxylic acid and the adenine subsite with a urea (*2*, *3*). These features likely contributed to poor membrane permeability and this, combined with the potential for metabolic liabilities, prompted us to search for alternative scaffolds. Inspired by compounds identified through virtual screening, we previously discovered the pyrrolidinone-containing AVI-219 as a single digit micromolar macrodomain inhibitor with favorable properties (*2*). Although optimization of this scaffold is ongoing, we were motivated to revisit fragment linking approaches because of the large number of fragment structures available, as well as ever expanding synthesis-on-demand libraries and improved computational methods to search the ultra-large libraries.

One of the promises of X-ray fragment screening is the ability to search vast regions of chemical space with a small number of compounds. However, building on fragment hits often requires synthesizing hundreds or even thousands of compounds. Using 3D similarity approaches like FrankenROCS; we can streamline the fragment expansion process by purchasing commercially available compounds. The advent of ultra-large synthesis-on-demand libraries has provided inexpensive access to tens of billions of molecules, however, exhaustive searches can be computationally prohibitive, even when accelerated with machine learning (ML) algorithms (*16*, *24*). Rather than exhaustively searching ultra-large libraries, alternative methods rely on the fact that libraries are synthesized from a relatively small number of chemical building blocks. This paper shows how Thompson sampling (TS) can dramatically reduce the computational requirements for searching ultra-large libraries and how improved filtering of those compounds can be achieved without human intervention. Similar approaches have searched ultra-large libraries by computationally docking individual building blocks and only expanding those with the highest scores (*10*, *25*). While these methods significantly reduce the runtime for virtual screens, they are limited by the accuracy of the method used to dock the building blocks.

While TS enables processing billions of molecules on commodity computing hardware, it still has some limitations. To use TS, a library cannot contain an arbitrary set of molecules: it must be available as reagents and reactions. Like other methods that search in reagent space, TS can sometimes focus too heavily on single building blocks and therefore select a set of similar molecules (*16*). Another confounding factor for any virtual screening method, including TS, is the identification of false positive molecules. We often have molecules whose high score is due to computational artifacts, such as incorrect tautomer assignments and torsional strain. One means of avoiding these artifacts is to employ more rigorous computational methods to filter the top 10,000 to 100,000 molecules. Newer methods that integrate ML and quantum chemical calculations are fast enough to be applied to thousands or even tens of thousands of molecules. Improvements to the TS process are an active area of research, and we hope to report additional advances in the future. The limited affinity we observe here is likely due to the limitations of the ROCS-based scoring, as we expect TS to have identified the best scoring molecules from the >22 Billion molecule REAL collection. While the strength of our approach is identifying alternative chemical scaffolds that effectively merge structurally-resolved fragments, other methods that leverage TS with score functions that better correlate with affinity may be more fruitful in identifying even more potent small molecules.

The most potent compound identified from the FrankenROCS approach was AVI-313, where the carboxylic acid of the parent fragment was replaced by a hydroxyl (**Fig. 2B**). Although the hydroxyl unexpectedly did not interact with the oxyanion subsite, subsequent optimization of AVI-313 led to several analogs with low μM IC_50_ values and where the hydroxyl hydrogen-bond with the oxyanion subsite, mimicking the parent fragment. This illustrates the power of shaped-based search methods and establishes the combination of the deazapurine head group with the 2-aminoethanol core as a privileged scaffold for targeting a macrodomain adenosine site. This scaffold has favorable membrane permeability compared to previously identified carboxylic acid-containing compounds (**Table 1**), and will allow multiple paths to lead optimization based on the distinct binding poses revealed by X-ray crystallography.

The optimization of AVI-313 offers lessons for the use of synthesis-on-demand chemistry and ultra-large libraries. Notably, chiral molecules were generally synthesized as stereoisomeric mixtures. This is a powerful strategy because the binding mode and affinity of stereoisomers can vary dramatically. This is illustrated by AVI-3669, which was synthesized as a mixture of four diastereomers, with electron density indicating that a single isomer was bound (**Fig. 6I**). One of the disadvantages of screening mixtures is that the effective concentration is inversely proportional to the number of isomers. However, because compounds were typically soaked at 10 mM into crystals, even a mixture of eight stereoisomers (e.g. a compound with three stereocenters) would reach 1.25 mM in crystallization drops and be detectable with the P4_3_ crystal system, where the protein concentration is 1.1 mM. Another related disadvantage to screening mixtures is that electron density maps may be ambiguous with regard to the isomer(s) bound, especially if multiple isomers of the same compound can bind simultaneously. This was seen for several compounds in the aromatic series (e.g. AVI- 3683, AVI-3640, AVI-3686 and AVI-3684). Tools such as PanDDA greatly assist the identification and modeling of these states, where the lower occupancy and overlapping binding sites makes traditional electron density maps difficult to interpret (*13*). The 137 Mac1 crystal structures reported in this work, including many weak binders, will aid the development of methods to automatically identify and model ligands where both conformational and compositional heterogeneity exists (*26*, *27*).

In summary, ultra-large synthesis-on-demand libraries like the Enamine REAL database allow for the broad exploration of chemical space during hit identification campaigns. However, the size of these databases makes an exhaustive evaluation difficult or sometimes impossible. Over the last few years, several active learning approaches have been developed to enable efficient searches through billions of molecules. In this work, we combined Thompson Sampling, an active learning technique capitalizing on the combinatorial nature of these large libraries, with FrankenROCS, a technique that uses pairs of fragments as 3D search queries.

Subsequent optimization achieved a fifty-fold potency gain without resorting to an expensive, time-consuming, bespoke chemistry effort. Future work will focus on combining features of the most potent ligands here, as well as further exploration of the northern region and phosphate tunnel. The approaches described in this work led to the discovery of several additional neutral scaffolds with low micromolar potency. Structurally validated, cell permeable and metabolically stable, these ligand efficient hits are poised for lead optimization and the *in vivo* testing of the effect of Mac1 inhibition on viral pathogenesis. Finally, the large number of structures, with accompanying assay data, will comprise a valuable dataset for driving innovations in ML-based methods for drug discovery.

## Methods

### FrankenROCS

#### Clustering of previous identified fragments

Our development of *FrankenROCS* relies on leveraging the information from the large fragment screen of Mac1, which identified 214 unique binders by X-ray crystallography (*1*). After separating the PDBs into separate chains, we used the PDB 5RS7, which contains only a single chain, as a reference structure for alignment. Next, we assigned bond orders to all ligands. Because five structures contained ligands bound in two different pockets, and one structure had three ligands bound in different pockets, we created unique copies of the receptor:ligand complex for each ligand location. We identified the center of mass of each ligand and used self-organizing maps to cluster the ligands. The workflow is available as a CoLab (https://colab.research.google.com/github/PatWalters/blog_posts/blob/main/align_proteins_extract_and_cluster

<u>_ligands.ipynb</u>). We inspected the resulting clusters to ensure that ligands could be used for the shape-based merging procedure outlined below. We identified all ligand pairs where the centers of mass were within 20 Å. We also excluded any pairs with van der Waals overlap between the fragments. This matching procedure resulted in 7181 pairs of fragments.

#### ROCS search from paired fragments

Each pair of ligands was concatenated into a single molecule representation using the function CombineMols from the RDKit. These merged RDKit molecules were used to create an input file for the OpenEye FastROCS program. FastROCS uses a graphics processing unit (GPU) to rapidly search a database containing conformers for millions of molecules. Since searches can be performed at a rate of more than 1 million conformers per second, the 7181 query molecules could be searched in a couple of hours. FastROCS operates by computationally superimposing a query molecule, Q, with conformers for each molecule, M, in a database, D. In this case, the query molecules, Q, are constructed from pairs of fragments, and the database, D, is composed of conformers from the 2.1 million molecules in the Enamine HTS collection. For each superposition, FastROCS calculates a measure of the overlap known as a Tanimoto Combo (TC) score. This score is the sum of two components, each between 0 and 1. The shape score evaluates the steric overlaps of the molecules, and the color score quantifies the overlap of similar pharmacophoric groups. We used this FrankenROCS workflow to search over the 2.1 million molecules in the Enamine HTS collection. The code required to run FrankenROCS is available on GitHub (https://github.com/PatWalters/frankenrocs).

#### Thompson sampling FrankenROCS

TS achieves efficiency gains by searching in reagent space rather than in product space. A pair of building blocks R1_n and R2_m are combined using a virtual reaction transform to generate a reaction product (P_nm). At each iteration, a set of three-dimensional conformers is generated for P_nm, and the maximum TC score from these conformers is associated with each building block (R1_n and R2_m). In an initial “warm-up” phase, each R1_n is randomly combined with three distinct R2_ building blocks, and each R2_m is randomly combined with three R1_ building blocks to generate the corresponding reaction products. The maximum TC scores are associated with the individual reagents, such that R1_n has at least 3 scores: the initial “warm-up” scores resulting from R1_n combined with 3 distinct R2_ building blocks and any scores from a product involving R1_n when it is selected as one of the 3 R1 building blocks during a R2_m “warm-up” phase. After the warm-up phase, each reagent will have an associated distribution of at least 3 scores, which is then transformed into a probability distribution used to guide sampling. The probability distributions continue to update in reagent space after each iteration as the algorithm exploits reagents that have a higher probability of generating a high scoring product. The TS terminates when there is no improvement in average reagent scores over 0.01% of the library, which indicates that the exploitation phase is no longer leading to changes in the probability distributions. Where the convergence criteria is not met after 0.1% of the total library has been scored, we also stop sampling for those reactions.

#### Using TS to search REAL

We used the REAL database from Enamine as a set of 242 reactions and 1,568,425 reagents. A fully enumerated version of this database would comprise more than 22 billion molecules. For each of the 97 dual fragment queries, we evaluated ∼22 million molecules with ROCS during TS (∼0.1% of the library). Of these, we selected 1,671,892 molecules with a TanimotoCombo score ≥ 1.2. Our initial inspection of a sample of the selected molecules revealed several molecules with torsion angles that appeared to be in high-energy conformations. To remove potentially strained molecules, we checked each ligand for torsional strain using the script RDKitFilterTorsionStrainEnergyAlerts.py from the MayaChemTools software suite. This script builds on previously published work highlighting the importance of ligand strain (*28*) by using distributions of dihedral angles from the Cambridge Structure Database to filter torsionally strained molecules. After filtering, 1,160,933 molecules remained. To reduce the computational overhead, we raised the TanimotoCombo threshold to 1.35, yielding 25,232 molecules.

These ligands were energy minimized into the binding site of PDB 5RSG using the superimposed pose. We could not correctly obtain molecular mechanics force field parameters for 18 molecules, which were removed from further consideration. Before minimization, the protein and ligand structures were prepared using the OEChemToolkit from OpenEye Scientific software. All water molecules were removed from the protein structures. Energy minimization was performed using the OpenForceField ff14sb_parsley as implemented in the OEForceField Toolkit from OpenEye Scientific Software. For each energy-minimized molecule, we recorded the RMSD from the initial pose and the energy of the complex. We performed additional torsional strain filtering to remove any strain induced during the minimization. We retained molecules with a total torsional energy score ≤ 6 and a maximum single torsion energy score ≤ 1.8. This additional post-minimization torsion filtering left 6,719 molecules.

Next, we removed molecules with steric clashes and those that moved from the original pose during minimization. We filtered cases where the total energy of the complex was >0 or the RMSD was >1.5. After this step, 1,791 molecules remained. When selecting molecules for synthesis, we wanted to confirm the interactions from the query fragments were preserved. To evaluate whether interactions were preserved, we calculated the similarity of interaction fingerprints for each query molecule and the corresponding posed and minimized ROCS hits. Interaction fingerprints were calculated using the ProLif protein-ligand interaction fingerprints. Rather than calculating a fingerprint similarity using a Tanimoto coefficient, which weights both molecules equally, we used an asymmetric Tversky index. The Tversky index is defined as

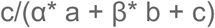

Where a is the number of “on” bits in fingerprint A, b is the number of “on” bits in fingerprint B, and c is the number of bits common to both fingerprints A and B. The parameters α and β adjust the weights of a and b. If we consider our query molecule as A and the molecule found through the ROCS search as B, we want to put more weight on a than b. We can do this by setting α to 1 and β to 0. After calculating and comparing interaction fingerprints, we retained molecules with a Tversky index ≥ 0.9. After filtering, 447 molecules remained.

To increase the diversity of molecules, we implemented three final filters. First, we wanted to avoid molecules that completely overlapped with the adenine moiety of the product ADP-ribose. We calculated a simple overlap volume between each posed, minimized ligand, and adenine (taken from X-ray structure 7TWX) to accomplish this. Molecules with a normalized overlap volume ≥ 0.85 were removed. After filtering, 223 molecules remained. Second, visual inspection of the remaining molecules highlighted known problematic functional groups, such as azides, furans, and nitro groups. Standard medicinal chemistry rules were designed to highlight reactive groups and assay interference to filter the results to 181 molecules (*29*). Upon inspection of the results from the previous step, we noticed several similar molecules. As a third and final step, we clustered the remaining molecules using the Taylor-Butina method implemented in the RDKit. Morgan fingerprints were used as molecular descriptors, and the clustering cutoff was set to 0.65. After clustering, we selected the molecule from each cluster with the highest Tversky index. The Thompson Sampling code is available on GitHub (https://github.com/PatWalters/TS).

#### Analog searching

To identify analogs of AVI-313, we searched the Enamine REAL database using the 2D molecular similarity search engine SmallWorld (http://sw.docking.org) and the substructure browser Arthor (http://arthor.docking.org) (*4*).

#### Protein expression and purification

The P4_3_ construct (residues 3-169, **Table S2**F) of the SARS-CoV-2 NSP3 macrodomain was expressed and purified as described previously (*1*, *2*, *17*). Briefly, plasmid DNA encoding the His_6_-tagged protein with a Tobacco Etch Virus (TEV) protease site for tag cleavage was transformed into BL21(DE3) *E. coli* and cells were grown on LB-agar supplemented with 100 μg/ml carbenicillin overnight at 37°C. Single colonies were used to inoculate small scale cultures (50 ml, lysogeny broth supplemented with 100 μg/ml carbenicillin, 150 ml baffled flasks). Cultures were grown at 37°C for eight hours with shaking at 180 rpm, then 10 ml of the small scale culture was used to inoculate a large scale culture (1 l terrific broth supplemented with 100 μg/ml carbenicillin, 2.8 l baffled flasks). Cultures were grown at 37°C until an optical density 0.8 at 600 nm, followed by 15 minutes at 4°C and induction of protein expression with 1 mM IPTG. The cultures were grown at 20°C for 14-16 hours and the cell pellets were harvested by centrifugation (4,000 g, 4°C, 20 minutes). Pellets were frozen at -80°C prior to purification.

All purification steps were performed on ice or at 4°C. Cells were resuspended in HisTrap loading buffer (500 mM NaCl, 50 mM TRIS pH 8.0, 10 mM imidazole, 2 mM DTT, 5% glycerol) supplemented with 2 U/ml Turbonuclease (Sigma, T4330-50KU) and lysed by sonication (Branson Sonifier 250, 5 minutes, 50% power, 0.5 second pulses). Cell debris was collected by centrifugation (30,000 g, 4°C, 30 minutes) and the lysate was filtered (0.22 μm, cellulose acetate) before loading onto a 5 ml HisTrap HP column (Cytiva, 17524801) pre-equilibrated with HisTrap loading buffer. The column was washed with 25 ml 5% HisTrap elution buffer (loading buffer supplemented with 300 mM imidazole) and then the protein was eluted with 25 ml 100% elution buffer.

Eluted fractions containing Mac1 were desalted into TEV reaction buffer (100 mM NaCl, 50 mM TRIS pH 8.0, 1 mM DTT, 1% glycerol) using a HiPrep desalt column (Cytiva, 17508701) and TEV protease was added to a mass:mass ratio of 1:20 TEV:Mac1. After incubation at 4°C overnight, the reaction mixture was loaded onto a 5 ml HisTrap HP column pre-equilibrated with TEV reaction buffer and the flow through containing cleaved protein was collected. After concentration with a 10 kDa molecular weight cutoff spin concentrator (Amicon, UFC901008) to 5 ml, the sample was loaded onto a HiLoad 16/600 Superdex 75 pg column (Cytiva, 28989333) pre-equilibrated with size exclusion buffer (150 mM NaCl, 20 mM TRIS pH 8, 5% glycerol, 2 mM DTT). The protein was eluted and concentrated to 40 mg/ml before being frozen in liquid nitrogen and stored at -80°C.

### Protein crystallization and ligand soaking

Crystals were grown using microseeding with sitting drop vapor diffusion as described previously (*1*, *2*, *17*). Briefly, protein (200 nl), reservoir solution (100 nl of 30 μl in each well, 28% PEG 3000 and 100 mM CHES pH 9.5) and seed stock (100 nl) were dispensed into wells of 3-well SwissCI crystallization plates (Hampton, HR3- 125) using a Mosquito crystallization robot (SPT Labtech). Plates were sealed (Hampton, HR4-521) and incubated at 19°C overnight. Compounds were prepared in DMSO to 100 mM (or the highest soluble concentration) and added to crystallization drops using an acoustic liquid handler (Echo 650, Beckman Coulter) to a final concentration of DMSO to 10-20% in drops (ligand concentrations varied from 3-20 mM depending on stock concentration) (**Table S2**B). Plates were incubated at room temperature for between 2-5 hours. Crystals were looped with the assistance of a Crystal Shifter (Oxford Lab Technologies) and vitrified in liquid nitrogen. To generate a background electron density map of TFA bound in the active site of Mac1, we soaked Mac1 crystals with 10 mM TFA (from a 100 mM stock prepared in DMSO). Crystals were vitrified as described previously.

### X-ray data collection, ligand identification, model building and refinement

X-ray diffraction data were collected at beamline 8.3.1 of the Advanced Light Source (ALS) and beamlines 9-2 and 12-2 of the Stanford Synchrotron Radiation Lightsource (SSRL). Data collection strategies are listed in **Table S2**C. Data were integrated and scaled with XDS (*30*) and merged with Aimless (*31*). Initial refinement for all datasets was performed using a refinement pipeline based on Dimple (*32*). The pipeline performed initial rigid body refinement with phenix.refine followed by two cycles of restrained refinement in Refmac, the first with four cycles of refinement with harmonic distance restraints (jelly-body restraints) and the second with eight cycles of restrained refinement. We used a 1.0 Å Mac1 structure refined against data collected from a DMSO soaked crystal as the starting model (dataset UCSF-P0075 in **Table S2**C) (*2*). Ligand binding was detected using the PanDDA algorithm (*13*) packaged in CCP4 version 7.0 (*33*) using a background electron density map calculated from 34 DMSO-soaked crystals as described previously (*1*, *2*). Where multiple datasets were recorded for a given ligand, we modeled the ligand with the highest occupancy based on inspection of F_O_-F_C_ difference maps and PanDDA event maps. Ligands with occupancy greater than 80% (based on the PanDDA background density correction value (1-BDC) and visual inspection), were modeled in COOT (version 0.9.6) (*34*) using F_O_-F_C_ difference electron density maps (shown in **Fig. S2**). Ligand restraint files for refinement were generated with phenix.elbow (*35*) from 3D coordinates generated by LigPrep (Schrödinger, version 2022-1). Structure refinement was performed with phenix.refine (*36*) (version 1.21.1-5286) starting from the Mac1 DMSO reference structure (UCSF-P0075). We initially performed rigid body refinement, followed by iterative cycles of refinement (coordinate, ADP and occupancy) and model building in COOT. Water placement was performed automatically in phenix.refine for the first four interactions, then the ligand was modeled and water picking was performed manually in subsequent rounds of refinement. For later cycles of refinement, hydrogens were added to the protein and ligand using phenix.readyset and were refined using a riding model. After one round of refinement with hydrogens, ADPs were refined anisotropically for non-hydrogen atoms. Four ligands were modeled and refined using this protocol (AVI-1504, AVI-607, AVI-3716 and AVI-4317). Refinement statistics are summarized in **Table S2**E.

The remaining ligands at <80% occupancy were modeled using PanDDA event maps and refined as multi-state models as described previously (*2*). Ligands were placed using COOT (version 0.8.9.2) and changes in protein conformation and solvent were modeled. Alternative conformations were modeled for residues when the RMSD exceeded 0.15 Å from the apo structure. The apo conformation was assigned alternative location (altloc) A and the ligand bound conformation was assigned altloc B. A chloride ion was modeled in the phosphate tunnel in 71/133 of the structures, typically forming a polar interaction with the ligand and water in the phosphate tunnel. The multi-state models were initially refined with five phenix.refine macrocycles without hydrogens, followed by five macrocycles with riding hydrogens followed by five macrocycles with anisotropic ADP refinement. Occupancy of the ligand- and apo-states was refined at either 2*(1-BDC) (*2*, *13*, *37*) or at ligand occupancy values from 10-90% at 10% increments. Ligand occupancy was determined by inspection of F_O_-F_C_ difference map peaks after refinement. The occupancy at 2*(1-BDC) was appropriate for 97/134 ligands, while the occupancy was decreased for three ligands (by 18% on average) and increased for 33 ligands (by 16% on average). For ligands where electron density for TFA binding to the oxyanion subsite overlapped with the ligand, we re-ran PanDDA with a background electron density map calculated from 44 datasets obtained from crystals soaked in 10 mM TFA (**Table S2**C). Using a 1.0 Å resolution dataset (UCSF-P3816 in **Table S2**C), we refined a model of the TFA-bound state as described above. Because the ligand-soaked crystals contained a mixture of the apo, ligand-bound and TFA-bound states, we refined models containing all three states, with the TFA-bound state assigned altloc T. Based on the pairwise Cα-Cα distances between the refined model from the reference DMSO-soaked structure and the reference TFA-soaked structure (**Fig. S6**A), we included alternative protein conformations for residues 155-160 in chain A, as well as TFA and two water molecules that were relatively well ordered in the TFA-bound structure (**Fig. S6**B,C). We initially refined the ligand occupancies at the 2*(1-BDC) value from the PanDDA run with the DMSO-background event or at values from 10-90% at 10% increments. For each of the ligand occupancies, we also refined models with varying TFA occupancy from 10% up to 80% at 10% increments, with the combined ligand+TFA occupancy capped at 90% (**Fig. S6**D). The TFA occupancy was determined by inspection of F_O_-F_C_ difference maps after refinement: TFA occupancies ranged from 10-40% (**Table S2**D). Examples of the PanDDA event maps with the DMSO and TFA map subtraction procedures are shown in **Fig. S6**E, along with the 2F_O_-F_C_ map after refinement of the multi-state model.

### Homogeneous time-resolved fluorescence (HTRF) assay

The HTRF assay was performed as described previously (*1*, *2*). Briefly, a Mac1 construct (residues 2-175, **Table S2**F) containing a His_6_-tag and a 3C-protease site was expressed and purified in the same manner as the P4_3_ construct, except the His_6_-tag was retained for labeling with the HTRF donor. After the first HisTrap step, the protein was immediately purified by size-exclusion chromatography using a buffer containing 300 mM NaCl, 25 mM 4-(2-hydroxyethyl)-1-piperazineethanesulfonic acid (HEPES) pH 7.5, 5% glycerol and 0.5 mM tris(2-carboxyethyl)phosphine (TCEP). The eluted fractions were pooled and concentrated to 5 mg/ml before freezing in liquid nitrogen and storage at -80°C. Human MacroD2 and Targ1 were purified in a similar manner, using constructs containing N-terminal His-tags (**Table S2**F) (*2*).

Compounds dissolved in DMSO were dispensed into white ProxiPlate-384 Plus 384-well assay plates (PerkinElmer,6008280) using an acoustic liquid handler (Echo 650, Beckman Coulter). Assays were conducted in a final volume of 16 μl with 12.5 nM Mac1, 200 nM peptide (ARTK(Bio)QTARK(Aoa- RADP)S, Cambridge Peptides), 1:20000 HTRF donor (Anti-His6-Eu^3+^ cryptate, PerkinElmer, AD0402) and 1:500 HTRF acceptor (Streptavidin-XL665, PerkinElmer 610SAXLB) prepared in assay buffer (25 mM HEPES) pH 7.0, 20 mM NaCl, 0.05% bovine serum albumin and 0.05% Tween-20). Mac1 protein and peptide (8 μl, prepared in the assay buffer) were dispensed using an electronic multichannel pipette and preincubated for 30 minutes at room temperature. Next, HTRF detection reagents were added (8 μl prepared in assay buffer) and the plates were incubated for 1 hour at room temperature. Fluorescence was measured using a Perkin Elmer EnVision 2105- 0010 Dual Detector Multimode microplate reader with dual emission protocol (A = excitation of 320 nm, emission of 665 nm, and B = excitation of 320 nm, emission of 620 nm). Compounds were tested in triplicate in a 14-point dose response. Raw data were processed to give an HTRF ratio (channel A/B × 10,000) and percentage competition was calculated relative to control reactions containing DMSO only (0% competition) and DMSO only and no enzyme (100% competition). IC_50_ values were calculated by fitting a sigmoidal dose-response equation using non-linear regression in GraphPad Prism (v.10.0.2, GraphPad Software, CA, USA) with the top constrained to 100% competition and the bottom constrained to 0% competition. *K*_i_ values were calculated as described previously (*2*), using the Cheng-Prusoff equation with the ADP-ribose conjugated peptide *K*_D_ equal to 2.68 μM. Binding of selected compounds to the human macrodomains Targ1 and MacroD2 was determined using the same method, except the concentration of Targ1 was 25 nM and the concentration of MacroD2 was 12.5 nM.

### Chemical synthesis

#### **AVI-1504** (Z3080312524)

(2R)-2-amino-3-methylbutan-1-ol (47 mg, 0.46 mmol), 4-chloro-7H-pyrrolo[2,3-d]pyrimidine (73 mg, 0.48 mmol) and triethylamine (Et_3_N) (60 mg, 0.59 mmol) were mixed in dry DMSO (2 ml). The reaction was sealed and stirred for 4 hours at 100°C. The mixture was cooled to ambient temperature and the solvent was evaporated under reduced pressure. The residue was dissolved in the DMSO (2 ml) and the solution was filtered, analyzed by LCMS, and purified by HPLC (0.1% NH_4_OH in water, 0%-20% MeCN with 0.1% NH_4_OH) to give (2R)-3- Methyl-2-({7H-pyrrolo[2,3-d]pyrimidin-4-yl}amino)butan-1-ol (AVI-1504) as a colorless sticky oil (17 mg, 17%). ^1^H NMR (500 MHz, DMSO-d6) δ 11.39 (br s, 1H), 8.02 (s, 1H), 7.01 (m, 1H), 6.89 (d, J = 8.88 Hz, 1H), 6.64 (m, 1H), 4.61 (m, 1H), 4.13 (m, 1H), 3.53 (m, 2H), 1.99 (m, 1H), 0.90 (m, 6H). LCMS (EI): m/z = 221.2 (MH+).

#### **AVI-607** (ZINC000082200734)

2-[(2R)-pyrrolidin-2-yl]propan-2-ol hydrochloride (70 mg, 0.42 mmol), 4-chloro-7H-pyrrolo[2,3-d]pyrimidine (70 mg, 0.46 mmol) and Et_3_N (96 mg, 0.95 mmol) were mixed in dry DMSO (2 ml). The reaction was sealed and stirred for 4 hours at 100°C. The mixture was cooled to ambient temperature and the solvent was evaporated under reduced pressure. The residue was dissolved in the DMSO (2 ml) and the solution was filtered, analyzed by LCMS, and purified by HPLC (0.1% NH_4_OH in water, 30%-80% MeOH with NH_4_OH) to give 2-[(2R)-1-{7H- Pyrrolo[2,3-d]pyrimidin-4-yl}pyrrolidin-2-yl]propan-2-ol (AVI-607) as a beige powder (36 mg, 34%). ^1^H NMR (500 MHz, DMSO-d6) δ 11.69 (s, 1H), 8.03 (s, 1H), 7.14 (m, 1H), 6.61 (m, 1H), 6.35 (br s, 1H), 4.51 (m, 1H), 3.94 (m, 1H), 3.88 (m, 1H), 2.06 (m, 1H), 1.88 (m, 3H), 1.11 (s, 3H), 1.02 (s, 3H). LCMS (EI) m/z = 247.2 (MH+).

#### **AVI-1445** (RLA-5681)

To a solution of 4-chloro-7H-pyrrolo[2,3-d]pyrimidine (120 mg, 0.78 mmol) in dry DMSO (5 ml) was added (S)- 2-(pyrrolidin-2-yl)propan-2-ol (110 mg, 0.86 mmol) and Et_3_N (400 mg, 3.9 mmol), the mixture was stirred at 110°C for 16 hours. The mixture was diluted with ethyl acetate (50 ml) and washed with water (10 ml), brine (10 ml). The organic layer was dried over Na_2_SO_4_ and concentrated under reduced pressure. The residue was purified by HPLC (0.1% NH_4_HCO_3_ in water, 5%-45% ACN) to give (S)-2-(1-(7H-pyrrolo[2,3-d]pyrimidin-4- yl)pyrrolidin-2-yl)propan-2-ol (AVI-1445) as a white solid (53 mg, yield: 27%). ^1^H NMR (500 MHz, MeOD) δ 8.06 (s, 1H), 7.12 (d, J = 3.6 Hz, 1H), 6.71 (d, J = 3.6 Hz, 1H), 4.56 – 4.38 (m, 1H), 4.16 (dt, J = 9.6, 6.9 Hz, 1H), 3.96 (dt, J = 10.3, 6.5 Hz, 1H), 2.19 – 2.03 (m, 2H), 2.01 – 1.82 (m, 2H), 1.28 (d, J = 17.1 Hz, 3H), 1.12 (s, 3H). LCMS (ESI): m/z= 247.3 (M+H)+.

#### **AVI-3758** (RLA-5748)

A mixture of 4-chloro-5-methyl-7H-pyrrolo[2,3-d]pyrimidine (15 mg, 90 μmol), 1-amino-2-methylpropan-2-ol (16 mg, 180 μmol) in 0.22 ml of mixed solvent IPA/H_2_O (10:1) was stirred at 100°C for 16 hours. The residue was purified by HPLC (water, 0%-20% ACN with 0.1% formic acid) to give 2-methyl-1-((5-methyl-7H-pyrrolo[2,3- d]pyrimidin-4-yl)amino)propan-2-ol, formic acid salt (AVI-3758) as a white solid (9.6 mg, yield: 40%). ^1^H NMR (400 MHz, MeOD) (mixture of rotamers was observed) δ 8.07 (s, 1H), 6.86 (br s, 1H), 3.58 (s, 2H), 2.47 (br s, 3H), 1.29 (br s, 6H). LCMS (ESI): m/z=221 (M+H)+.

#### **AVI-3571** (RLA-5703)

A mixture of 4-chloro-5-methyl-7H-pyrrolo[2,3-d]pyrimidine (15 mg, 90 μmol), 1-(aminomethyl)cyclobutan-1-ol (18 mg, 180 μmol) in 0.22 lL of mixed solvent IPA/H2O (10:1) was stirred at 100°C for 4 days. The residue was purified by HPLC (water, 0%-5% ACN with 0.1% formic acid) to give 1-(((5-methyl-7H-pyrrolo[2,3-d]pyrimidin- 4-yl)amino)methyl)cyclobutan-1-ol (AVI-3571), formic acid salt as a white solid (7.7 mg, yield: 31%). ^1^H NMR (400 MHz, MeOD) δ 8.08 (s, 1H), 6.87 (s, 1H), 3.74 (s, 2H), 2.46 (s, 3H), 2.19-2.07 (m, 4H), 1.83-1.75 (m, 1H), 1.70-1.63 (m, 1H). LCMS (ESI): m/z=233 (M+H)+.

#### **AVI-3760** (RLA-5750)

A mixture of 4-chloro-5-methyl-7H-pyrrolo[2,3-d]pyrimidine (10 mg, 60 μmol), 1-(aminomethyl)cyclopentan-1-ol, HCl (18 mg, 120 μmol) and Et_3_N (30.2 mg, 41.6 μmol) in 0.22 ml of DMSO was stirred at 100°C for 3 days.

The residue was purified by HPLC (water, 0%-40% ACN with 0.1% formic acid) to give 1-(((5-methyl-7H- pyrrolo[2,3-d]pyrimidin-4-yl)amino)methyl)cyclopentan-1-ol, formic acid salt (AVI-3760) as a white solid (9.2 mg, yield: 53%). ^1^H NMR (400 MHz, MeOD) (mixture of rotamers was observed) δ 8.07 (br s, 1H), 6.86 (br s, 1H), 3.67 (s, 2H), 2.47 (br s, 3H), 1.89-1.67 (m, 8H). LCMS (ESI): m/z=247 (M+H)+.

#### **AVI-3761** (RLA-5751)

A mixture of 4-chloro-5-methyl-7H-pyrrolo[2,3-d]pyrimidine (20.0 mg, 119 μmol), 1-(aminomethyl)cyclohexan-1- ol, HCl (40 mg, 240 μmol) and Et_3_N (60 mg, 600 μmol) in 0.2 ml DMSO was stirred at 100°C for 18 hours. The residue was purified by HPLC (water, 0%-40% ACN with 0.1% formic acid) to give 1-(((5-methyl-7H- pyrrolo[2,3-d]pyrimidin-4-yl)amino)methyl)cyclohexan-1-ol (AVI-3761), formic acid salt as a white solid (21 mg, yield: 56%). ^1^H NMR (MeOH-d4, 400 MHz) (mixture of rotamers was observed) δ 8.08 (br s, 1H), 6.87 (s, 1H), 3.60 (s, 2H), 2.47 (br s, 3H), 1.55-1.65 (m, 10H). LCMS (ESI): m/z=261 (M+H)+.

#### **AVI-1444** (RLA-5679)

To a solution of 4-chloro-7H-pyrrolo[2,3-d]pyrimidine (150 mg, 1.0 mmol) in dry DMSO (5 ml) was added (3R,4S)-4-aminotetrahydrofuran-3-ol (110 mg, 1.1 mmol) and t-BuOK (560 mg, 5 mmol), the mixture was stirred at 110°C for 3 hours. The mixture was diluted with ethyl acetate (50 ml) and washed with water (10 ml), brine (10 ml). The organic layer was dried over Na_2_SO_4_ and concentrated under reduced pressure. The residue was purified by HPLC (0.1% NH_4_HCO_3_ in water, 5%-35% ACN) to give (3R,4S)-4-((7H-pyrrolo[2,3- d]pyrimidin-4-yl)amino)tetrahydrofuran-3-ol (AVI-1444) as a white solid (103 mg, yield: 47%). ^1^H NMR (500 MHz, MeOD) δ 8.35 (s, 1H), 7.25 (dd, J = 7.8, 3.6 Hz, 1H), 6.53 (d, J = 3.4 Hz, 1H), 5.44 – 5.35 (m, 1H), 4.37 – 4.25 (m, 1H), 4.14 (dd, J = 8.9, 5.4 Hz, 1H), 3.98 (d, J = 10.7 Hz, 1H), 3.69 – 3.55 (m, 2H). LCMS (ESI): m/z= 221.3 (M+H)+.

#### **AVI-4194** (RLA-5827)

To a solution of 4-chloro-5-methyl-7H-pyrrolo[2,3-d]pyrimidine (150 mg, 0.89 mmol) in dry DMSO (5 ml) was added 4-(aminomethyl)tetrahydro-2H-pyran-4-ol (120 mg, 0.89 mmol) and Et_3_N (180 mg, 1.8 mmol), the mixture was stirred at 110°C for 16 hours. The mixture was diluted with ethyl acetate (50 ml) and washed with water (10 ml), brine (10 ml). The organic layer was dried over Na_2_SO_4_ and concentrated under reduced pressure. The residue was purified by HPLC (0.1% NH_4_HCO_3_ in water, 5%-40% ACN) to give 4-(((5-methyl- 7H-pyrrolo[2,3-d]pyrimidin-4-yl)amino)methyl)tetrahydro-2H-pyran-4-ol (AVI-4194) as a white solid (33 mg, yield: 14%). ^1^H NMR (500 MHz, MeOD-d4) δ 8.05 (s, 1H), 6.83 (d, J = 0.9 Hz, 1H), 3.86 – 3.68 (m, 4H), 3.63 (s, 2H), 2.45 (d, J = 0.8 Hz, 3H), 1.80 – 1.66 (m, 2H), 1.60 (d, J = 12.9 Hz, 2H). LCMS (ESI): m/z= 261.1 (M- H)-.

#### **AVI-4195** (RLA-5828)

To a solution of 4-chloro-5-methyl-7H-pyrrolo[2,3-d]pyrimidine (400 mg, 2.4 mmol) in dry DMSO (10 ml) was added 4-(aminomethyl)tetrahydro-2H-thiopyran-4-ol (350 mg, 2.4 mmol) and Et_3_N (480 mg, 4.8 mmol), the mixture was stirred at 110°C for 16 hours. The mixture was diluted with ethyl acetate (100 ml) and washed with water (30 ml), brine (30 ml). The organic layer was dried over Na_2_SO_4_ and concentrated under reduced pressure. The residue was purified by HPLC (0.1% NH_4_HCO_3_ in water, 5%-45% ACN) to give 4-(((5-methyl- 7H-pyrrolo[2,3-d]pyrimidin-4-yl)amino)methyl)tetrahydro-2H-thiopyran-4-ol (AVI-4195) as a white solid (300 mg, yield: 45%). ^1^H NMR (500 MHz, MeOD-d4) δ 8.05 (s, 1H), 6.82 (s, 1H), 3.57 (s, 2H), 3.01 – 2.83 (m, 2H), 2.46 (d, J = 18.7 Hz, 5H), 1.93 (d, J = 15.8 Hz, 2H), 1.85 – 1.70 (m, 2H). LCMS (ESI): m/z= 279.2 (M+H)+.

### Solubility, permeability and metabolic stability

#### Kinetic solubility

Compounds dissolved in DMSO (10 mM) were diluted with 190 μl of phosphate buffered saline (pH 7.4) in 96- well solubility filter plates (Millipore, MSSLBPC50). Solutions were mixed by shaking for 1.5 hours at room temperature and samples were filtered by vacuum into fresh 96-well plates. Compound concentration was determined by absorbance (220 nm, 254 nm, and 280 nm) relative to a three-point standard curve prepared in DMSO (500, 50 and 5 μM). Mean solubility was determined from triplicate measurements.

#### Bi-directional Transport in Caco-2 Cells

Caco-2 cell plates were obtained commercially and were maintained for 21 days at 37°C with 5% CO_2_. Cells were washed with Hank’s Balanced Salt Solution (HBSS) with 5 mM HEPES for 30 minutes before starting the experiment. Test compound solutions are prepared by diluting DMSO stock into HBSS buffer, resulting in a final DMSO concentration of 0.1%. Prior to the assay, cell monolayer integrity was verified by transendothelial electrical resistance (TEER) to ensure that wells have resistance above the acceptance cut-off (1 kΩ).

Transport experiments were initiated by adding test compounds to the apical (75 μl) or basal (250 μl) side. Transport plates were incubated at 37°C in a humidified incubator with 5% CO_2_. Samples were taken from the donor and acceptor compartments after one hour and analyzed by liquid chromatography with tandem mass spectrometry (LC/MS/MS) using an AB Sciex API 4000 instrument, coupled to a Shimadzu LC-20AD LC Pump system. Samples were separated using a Waters Atlantis T3 dC18 reverse phase HPLC column (20 mm x 2.1 mm) at a flow rate of 0.5 ml/min. The mobile phase consisted of 0.1% formic acid in acetonitrile (solvent A) and 0.1% formic acid in water (solvent B).

Apparent permeability (P_app_) values were calculated using the following equation:

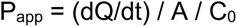

where dQ/dt is the initial rate of compound transport across the cell monolayer, A is the surface area of the filter membrane, and C_0_ is the initial concentration of the test compound, calculated for each direction using a four-point calibration curve by LC/MS/MS.

Net flux ratio between the two directional transports was calculated by the following equation:

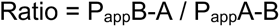

where P_app_B-A and P_app_A-B represent the apparent permeability of compounds from the basal-to-apical and apical-to-basal side of the cellular monolayer, respectively. Recovery was calculated based on the compound concentration at the end of the experiment, compared to that at the beginning of the experiment, adjusted for volumes.

#### Microsomal stability

The assay was carried out in 96-well plates at 37°C. Reaction mixtures (25 μl) contained a final concentration of 1 μM compound, 0.5 mg/ml liver microsomes protein, and 1 mM NADPH and/or 1 mM UDPGA (with alamethicin) in 100 mM potassium phosphate (pH 7.4) with 3 mM MgCl_2_. After 0, 15, 30, and 60 minutes, 150 μl of quench solution (100% acetonitrile with 0.1% formic acid) with internal standard was transferred to each well. Reactions containing the same components except the NADPH were prepared as the negative control. Plates were sealed, vortexed, and centrifuged at 4°C for 15 minutes at 4000 rpm. The supernatant was transferred to fresh plates for LC/MS/MS analysis using an AB Sciex API 4000 instrument, coupled to a Shimadzu LC-20AD LC Pump system. Analytical samples are separated using a Waters Atlantis T3 dC18 reverse phase HPLC column (20 mm x 2.1 mm) at a flow rate of 0.5 ml/min. The mobile phase consisted of 0.1% formic acid in water (solvent A) and 0.1% formic acid in 100% acetonitrile (solvent B).

The extent of metabolism was calculated as the disappearance of compounds, compared to the 0-min time incubation. Initial rates were calculated for the compound concentration and used to determine t1/2 values and subsequently, the intrinsic clearance, CLint = (0.693)(1/ t1/2 (min))(g of liver / kg of body weight)(ml incubation/mg of microsomal protein)(45 mg of microsomal protein/g of liver weight).

## Acknowledgements

This work was supported by U19AI171110, a Sanghvi-Agarwal Innovation Award (J.S.F.), and a sponsored research agreement with Relay Therapeutics (J.S.F.). The synchrotron X-ray diffraction data used to determine Mac1 structures were collected at beamline 8.3.1 of the Advanced Light Source (ALS) and beamlines 9-2, 12-1 and 12-2 of the Stanford Synchrotron Radiation Lightsource (SSRL). The ALS, a U.S. DOE Office of Science User Facility under contract no. DE-AC02-05CH11231, is supported in part by the ALS-ENABLE program funded by the NIH, National Institute of General Medical Sciences, grant P30 GM124169. Use of the SSRL, SLAC National Accelerator Laboratory, is supported by the U.S. Department of Energy, Office of Science, Office of Basic Energy Sciences under Contract No. DE-AC02-76SF00515. The SSRL Structural Molecular Biology Program is supported by the DOE Office of Biological and Environmental Research, and by the National Institutes of Health, National Institute of General Medical Sciences (P30GM133894).

Coordinates and structure factors have been deposited in the Protein Data Bank with the accession codes: 7HCB, 7HCC, 7HCD, 7HCE, 7HCF, 7HCG, 7HCH, 7HCI, 7HCJ, 7HCK, 7HCL, 7HCM, 7HCN, 7HCO, 7HCP, 7HCQ, 7HCR, 7HCS, 7HCT, 7HCU, 7HCV, 7HCW, 7HCX, 7HCY, 7HCZ, 7HD0, 7HD1, 7HD2, 7HD3, 7HD4, 7HD5, 7HD6, 7HD7, 7HD8, 7HD9, 7HDA, 7HDB, 7HDC, 7HDD, 7HDE, 7HDF, 7HDG, 7HDH, 7HDI, 7HDJ, 7HDK, 7HDL, 7HDM, 7HDN, 7HDO, 7HDP, 7HDQ, 7HDR, 7HDS, 7HDT, 7HDU, 7HDV, 7HDW, 7HDX, 7HDY, 7HDZ, 7HE0, 7HE1, 7HE2, 7HE3, 7HE4, 7HE5, 7HE6, 7HE7, 7HE8, 7HE9, 7HEA, 7HEB, 7HEC, 7HED, 7HEE, 7HEF, 7HEG, 7HEH, 7HEI, 7HEJ, 7HEK, 7HEL, 7HEM, 7HEN, 7HEO, 7HEP, 7HEQ, 7HER, 7HES, 7HET, 7HEU, 7HEV, 7HEW, 7HEX, 7HEY, 7HEZ, 7HF0, 7HF1, 7HF2, 7HF3, 7HF4, 7HF5, 7HF6, 7HF7, 7HF8, 7HF9, 7HFA, 7HFB, 7HFC, 7HFD, 7HFE, 7HFF, 7HFG, 7HFH, 7HFI, 7HFJ, 7HFK, 7HFL, 7HFM, 7HFN, 7HFO, 7HFP, 7HFQ, 7HFR, 7HFS, 7HFT, 7HFU, 7HFV, 7HFW, 7HFX, 7HFY, 7HFZ, 9D6B, 9D6G, 9D6H, 9D6I. Structure factor intensities (unmerged, merged, and merged/scaled), PanDDA input and output files including Z-map and event maps in CCP4 format, and refined models including the ligand-bound state extracted from multi-state models were uploaded to Zenodo (DOI: 10.5281/zenodo.13368547).

