## Supplemental Information for "Extensive exploration of structure activity relationships for the SARS-CoV-2 macrodomain from shape-based fragment merging and active learning"

**Affiliations**
[1]Department of Bioengineering and Therapeutic Sciences, University of California San Francisco, San Francisco, CA 94158.
[2]Department of Pharmaceutical Chemistry, University of California San Francisco, San Francisco, CA 94158.
[3]Helen Diller Family Comprehensive Cancer Center, University of California San Francisco, San Francisco, CA 94158.
[4]Relay Therapeutics, Cambridge, MA 02142.

[†]Contributed equally

**Table S1**. Summary of compound selection and filtering for FrankenROCS pipelines.

| Pipeline | Step | Action | Criteria | Number of Ligands |
|---|---|---|---|---|
| FrankenROCS (Enamine HTS collection) | 1 | Initial x-ray structures | | 235 complexes |
| | 2 | Align to 5RS7 | | 235 complexes |
| | 3 | Assign bond orders and extract ligands | | 242 ligands |
| | 4 | Calculate distances between ligand centers | < 20 Å + no overlap | 7181 pairs |
| | 5 | FastROCS w/Enamine HTS | TC ≥ 1.3 | 692 ligands |
| | 6 | Manual selection | | 39 ligands |
| FrankenROCS with Thompson Sampling (TS-FrankenROCS (Enamine REAL database) | 1 | Enamine REAL | 97 queries with Thomson Sampling | 22,000,000,000 |
| | 2 | TS ROCS | Tanimoto Combo ≥ 1.35 | 25,214 |
| | 3 | Energy minimize complexes | | 25,214 |
| | 4 | Filter torsions | Total ≤ 6 and Single ≤ 1.8 | 6,719 |
| | 5 | Filter RMS and strain | RMS ≤1.5 and Energy < 0 | 1,791 |
| | 6 | Interaction fingerprint | Tversky similarity > 0.9 | 447 |
| | 7 | Substrate overlap | Overlap < 0.85 | 221 |
| | 8 | Functional group filters | Filtering rules | 181 |
| | 9 | Clustering | Taylor Butina cutoff 0.65 | 36 |

**Table S2**. Summary of compounds tested in this work. Excel spreadsheet.

**Table S3.** Solubility, permeability and mouse microsomal metabolic stability for AVI-313 analogs described herein, and for previously described lead compounds AVI-92 and AVI-219.

| Compound | **AVI-92**[a] | **AVI-219**[a] | **AVI-3669** | **AVI-1504** | **AVI-3761** | **AVI-607** | **AVI-3716** |
|---|---|---|---|---|---|---|---|
| Structure | 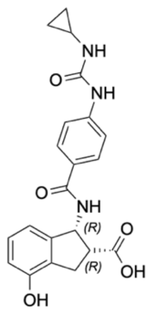 | 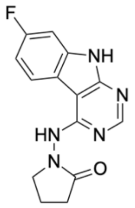 | 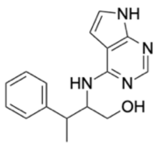 | 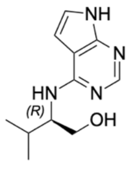 | 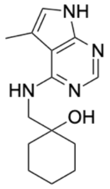 | 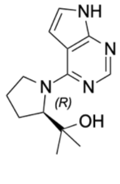 | 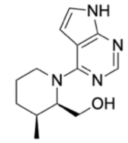 |
| PDB code | 5SQJ | 5SRY | 7HEC | 9D6H | 7HF1 | 9D6B | 9D6G |
| HTRF $IC_{50}$ (µM) | 0.5 | 1.2 | 8.7 | 9.8 | 13 | 5.1 | 4.7 |
| $K_i$ (µM) | 0.5 | 1.1 | 8.1 | 9.1 | 12 | 4.7 | 4.4 |
| M.W. (g/mol) | 395.4 | 285.3 | 282.4 | 220.3 | 260.3 | 246.3 | 246.3 |
| Heavy atoms | 29 | 21 | 21 | 16 | 19 | 18 | 18 |
| Ligand efficiency[c] | 0.30 | 0.39 | 0.33 | 0.44 | 0.36 | 0.41 | 0.41 |
| Caco-2 $P_{app}$ A-B ($10^{-6}$cm/s) | 0.04 | 10.6 | 19.4 | 14.9 | 21.5 | 22.0 | 25.9 |
| Caco-2 $P_{app}$ B-A ($10^{-6}$cm/s) | 1.7 | 16.8 | 14.5 | 11.3 | 11.2 | 15.7 | 18.9 |
| Caco-2 ratio B-A/A-B | 40.7 | 1.6 | 0.7 | 0.8 | 0.52 | 0.7 | 0.7 |
| Mean solubility (µM) | 524 | 127 | 529 | 370 | 555 | 518 | 523 |
| Mouse microsome $T_{1/2}$ (min) | >120 | 26.1 | 8.3 | >120 | 7.49 | 19 | 31.2 |
| CLint (mouse) (µl/min/mg protein) | <11.6 | 53.1 | 166.4 | <11.6 | 185 | 73 | 44.4 |

[a]Reported in (2)

[c]Binding energy per heavy atom.

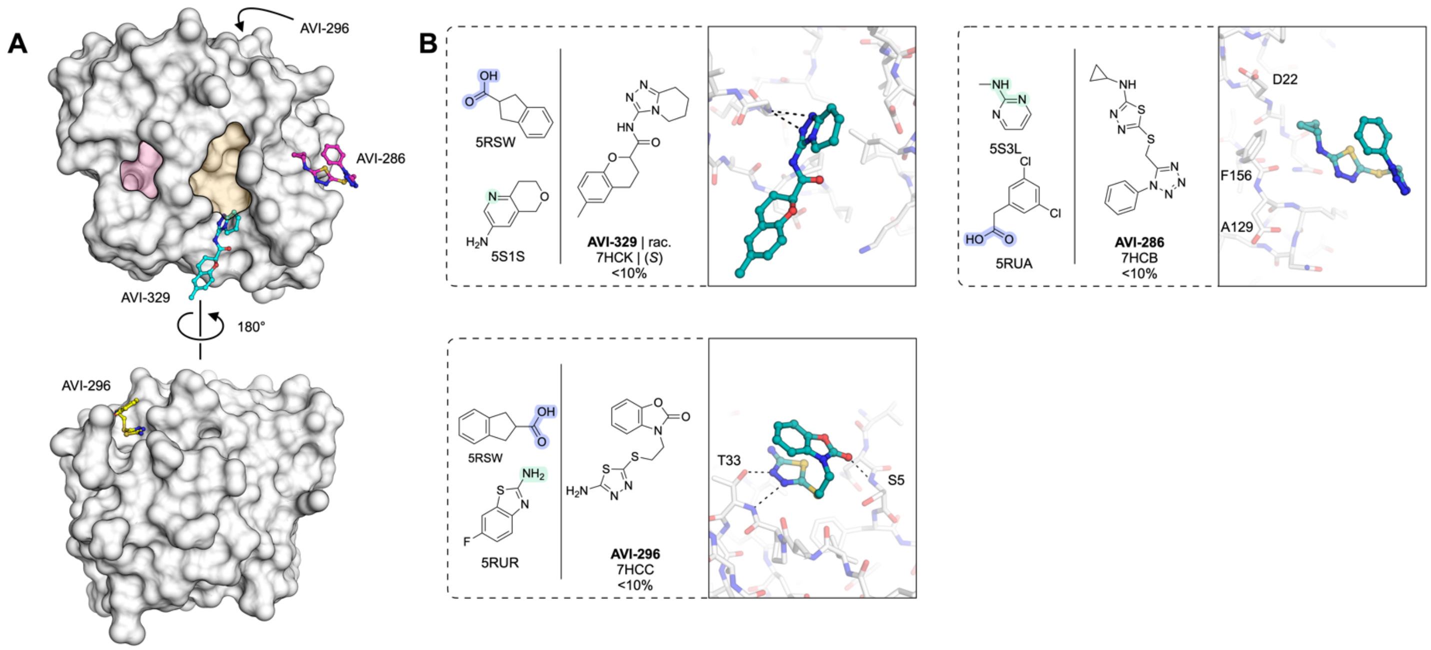


**Figure S1.** Three hits from the FrankenROCS pipeline were identified binding outside of the Mac1 adenosine site. (**A**) Surface view of Mac1 with the adenosine (yellow) and terminal ribose sites (red) shaded. (**B**) Chemical structures of the hits shown in (**A**). X-ray crystal structures are shown with hydrogen bonds shown as dashed black lines. Results for the preliminary HTRF screen (% competition at 200 μM, mean of three technical replicates) are indicated.

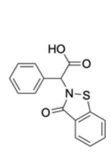

**AVI-288** | rac.
7HCN | (*S*)
21% | >1 mM

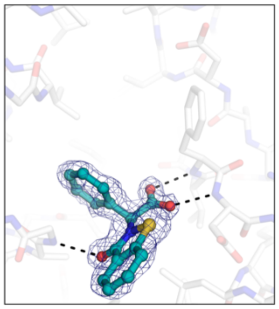

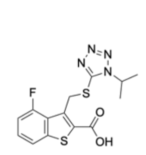

**AVI-291**
7HCO
41% | >1 mM

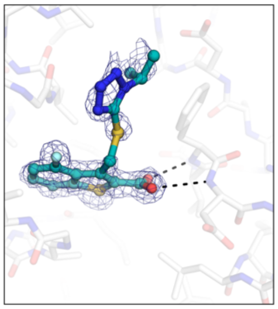

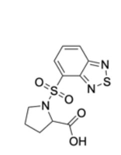

**AVI-2** | rac.
7HCE
23%

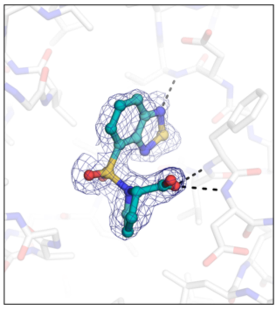

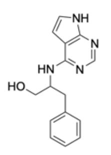

**AVI-313** | rac.
7HCF | (*S*), (*R*)
75% | 130 µM

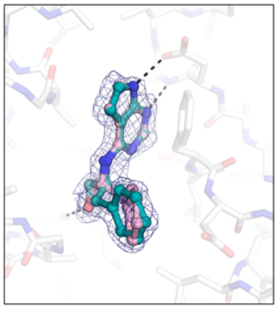

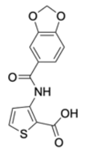

**AVI-316**
7HCG
29% | >1 mM

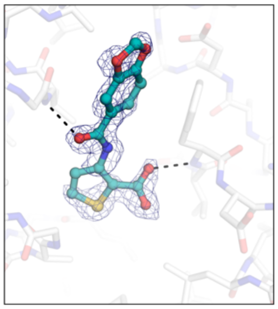

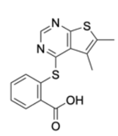

**AVI-317**
7HCH
24% | >1mM

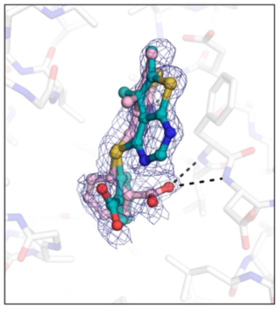

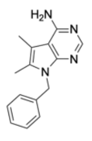

**AVI-321**
7HCI
19%

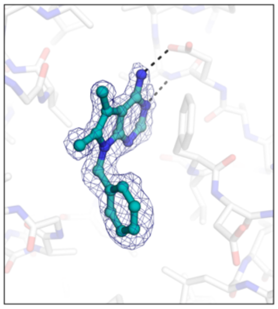

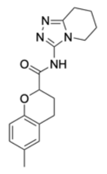

**AVI-329** | rac.
7HCK | (*S*)
<10%

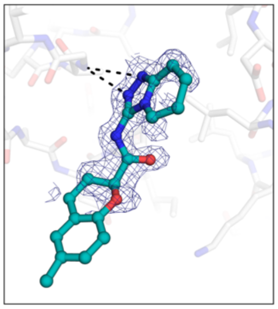

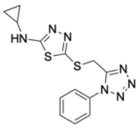

**AVI-286**
7HCB
<10%

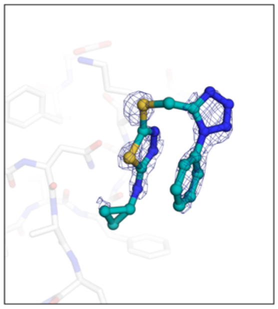

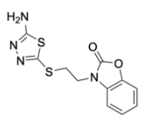

**AVI-296**
7HCC
<10%

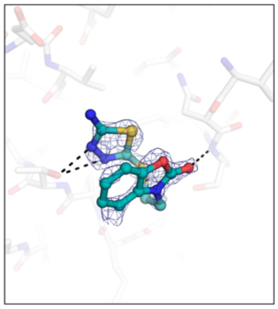

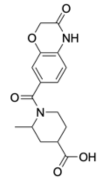

**AVI-303** | diast.
7HCD | (*S*,*R*)
<10%

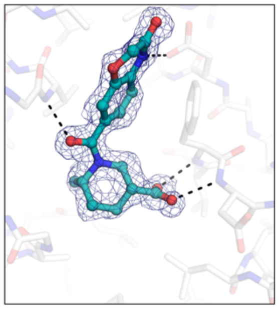

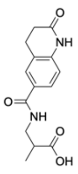

**AVI-328** | rac.
7HCJ | (*R*), (*S*)
50% | 220 µM

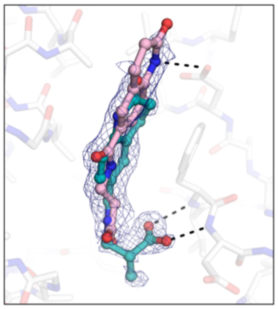

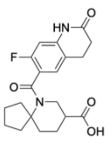

**AVI-338** | rac.
7HCP | (*R*)
23% | 800 µM

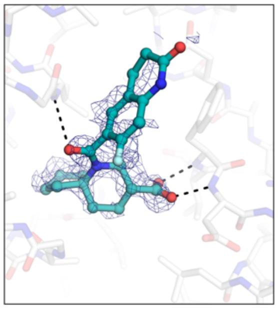

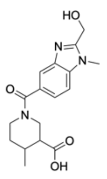

**AVI-348** | diast.
7HCM | (*R*,*R*)
<10%

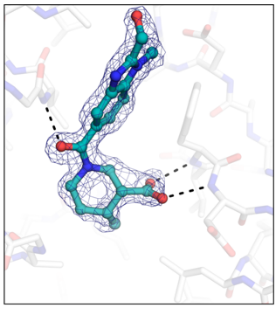

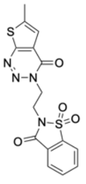

**AVI-341**
7HCL
11%

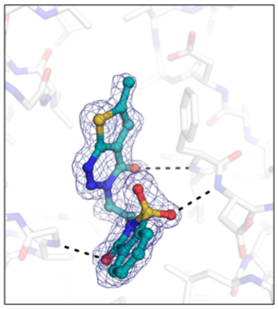

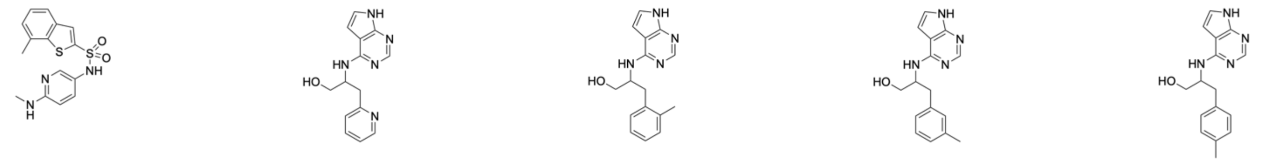

**AVI-345**
7HCQ
18%

**AVI-3696** (I) | rac.
7HEV | (*R*), (*S*)
34%

**AVI-3674** (I) | rac.
7HEG | (*R*)
32%

**AVI-3675** (I) | rac.
7HEH | (*R*)
42%

**AVI-3683** (I) | rac.
7HEL | (*S*), (*R*)
14%

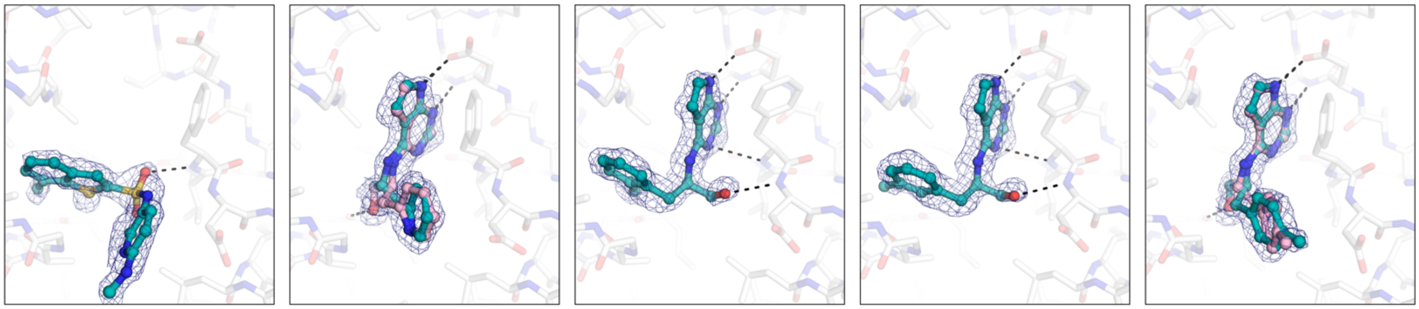

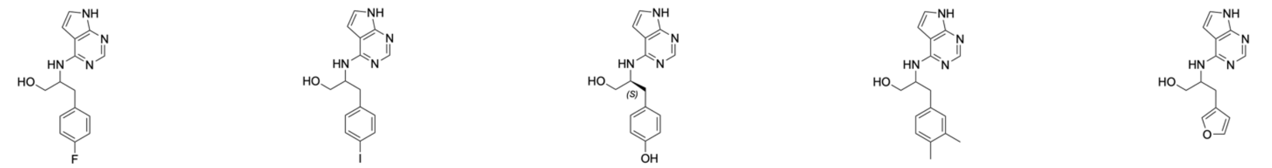

**AVI-3640** (I) | rac.
7HDR | (*S*), (*R*)
38%

**AVI-3686** (I) | rac.
7HEO | (*S*), (*R*)
<10%

**AVI-4066** (I) | (*S*)
7HF4 | (*S*)
720 µM

**AVI-3684** (I) | rac.
7HEM | (*R*), (*S*)
<10%

**AVI-3705** (I) | rac.
7HD2 | (*S*)
64%

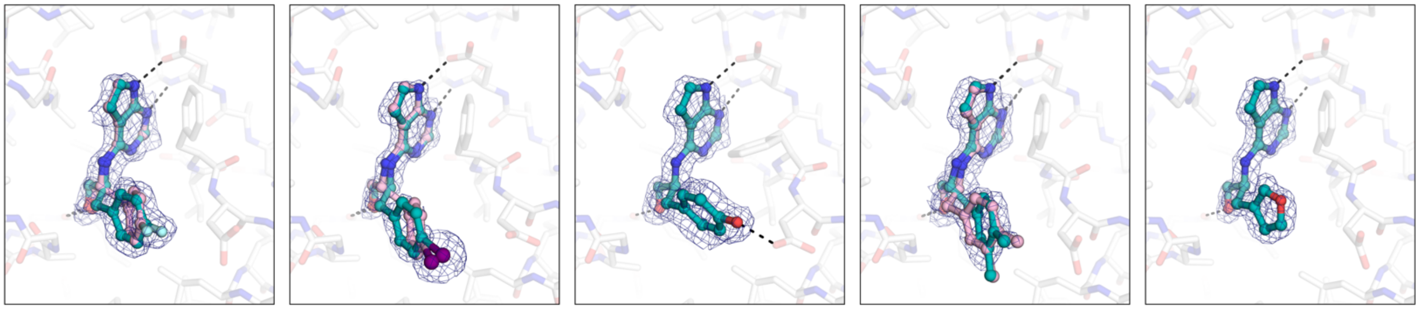

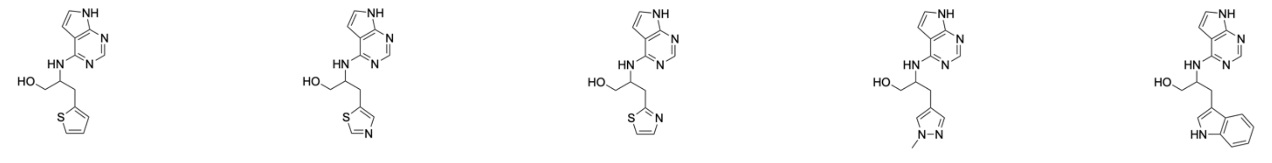

**AVI-3711** (I) | rac.
7HD8 | (*S*)
63%

**AVI-3703** (I) | rac.
7HD0 | (*S*)
28%

**AVI-3673** (I) | rac.
7HEF | (*S*)
38%

**AVI-3657** (I) | rac.
7HE3 | (*S*)
24%

**AVI-3650** (I) | rac.
7HDY | (*R*), (*S*)
71%

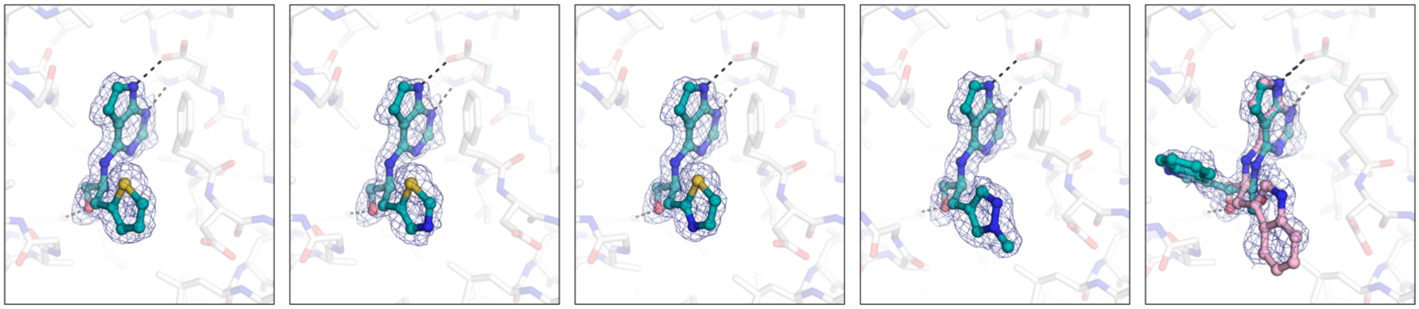

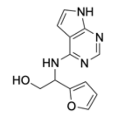


**AVI-3659** (I) | rac.
7HE5 | (*R*)
55%

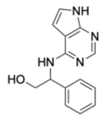


**AVI-3715** (I) | rac.
7HDA | (*S*)
54%

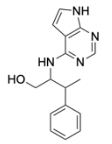


**AVI-3669** (I) | diast.
7HEC | (*R*,*S*)
98% | 8.7 µM

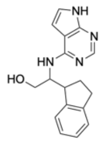


**AVI-3693** (I) | diast.
7HET | (*R*,*R*), (*S*,*R*)
78%

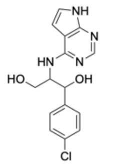


**AVI-3718** (I) | diast.
7HDC | (*R*,*R*), (*S*,*R*), (*R*,*R*)
78%

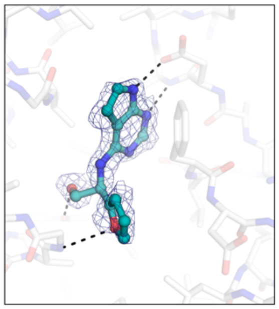

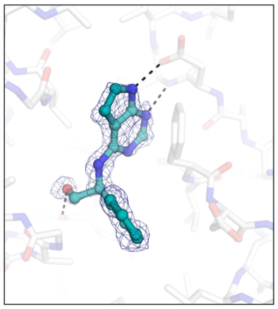

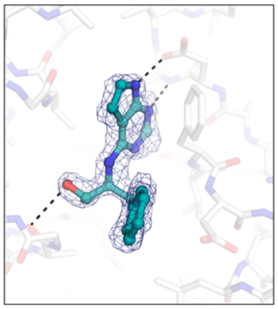

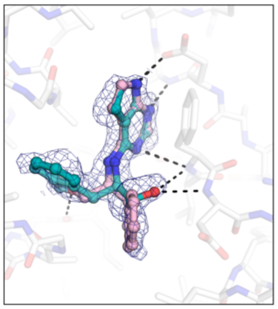

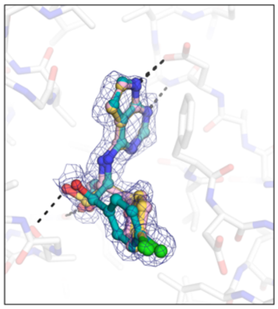

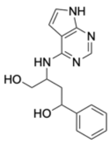


**AVI-3642** (I) | diast.
7HDT | (*S*,*S*)
<10%

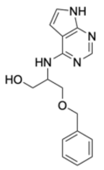


**AVI-3709** (I) | rac.
7HD5 | (*S*), (*R*)
29%

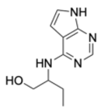


**AVI-3643** (I) | rac.
7HDU | (*R*)
63%

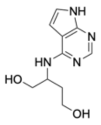


**AVI-3734** (I) | rac.
7HDN | (*R*)
86% | 52 µM

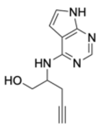


**AVI-3644** (I) | rac.
7HDV | (*S*)
43%

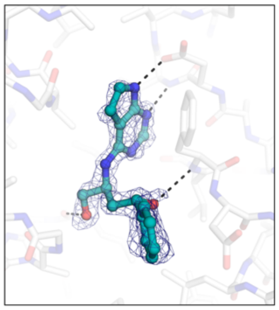

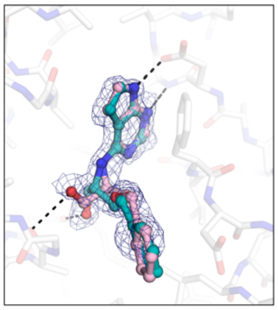

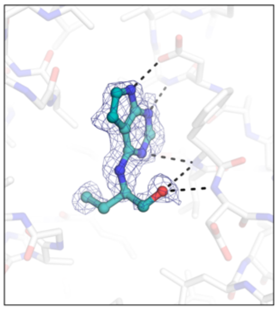

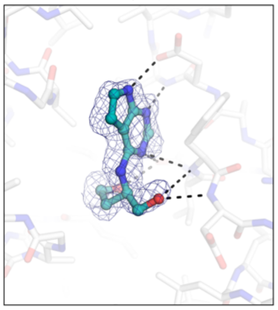

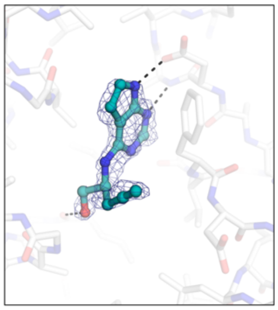

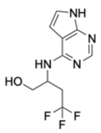


**AVI-3637** (I) | rac.
7HDO | (*S*)
50%

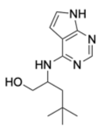


**AVI-3648** (I) | rac.
7HDX | (*S*)
27%

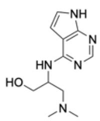


**AVI-3671** (I) | rac.
7HEE | (*S*)
<10%

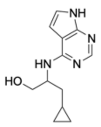


**AVI-3662** (I) | rac.
7HE7 | (*S*)
44%

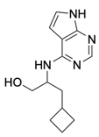


**AVI-3641** (I) | rac.
7HDS | (*S*)
32%

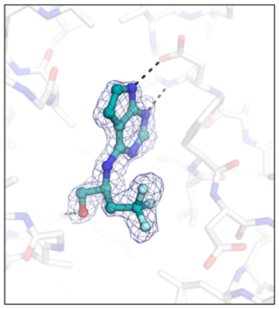

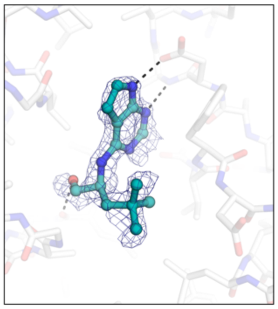

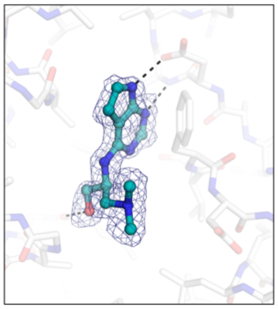

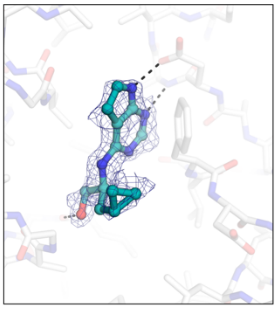

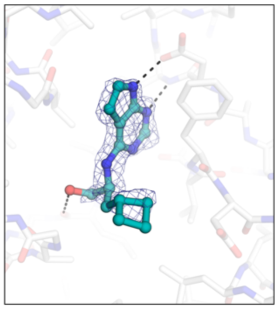

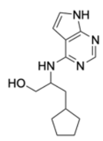

**AVI-3691** (I) | rac.
7HER | (*S*)
53%

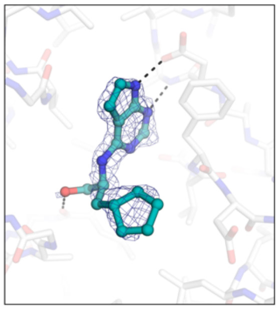

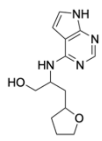

**AVI-3660** (I) | diast.
7HE6 | (*S*,*R*)
18%

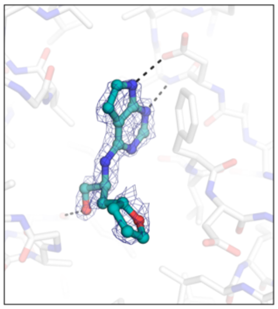

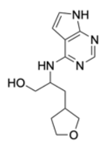

**AVI-3665** (I) | diast.
7HE9 | (*S*,*S*)
11%

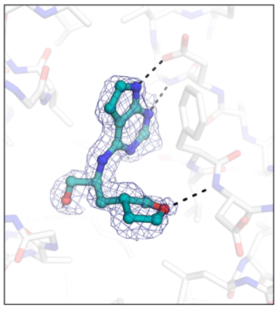

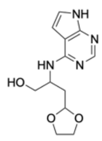

**AVI-3658** (I) | rac.
7HE4 | (*S*)
11%

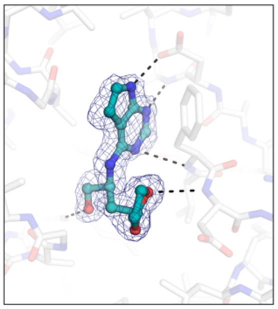

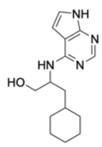

**AVI-3690** (I) | rac.
7HEQ | (*R*)
66%

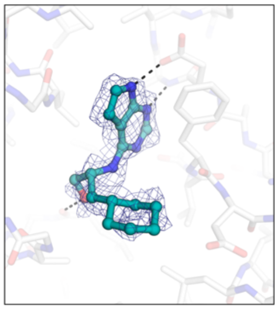

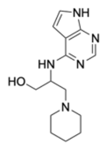

**AVI-3677** (I) | rac.
7HEI | (*S*)
<10%

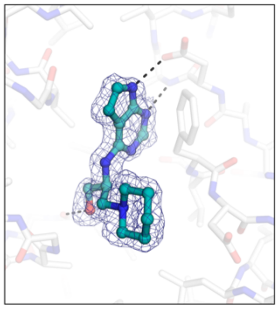

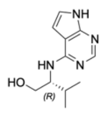

**AVI-1504** (I) | (*R*)
9D6H | (*R*)
97% | 9.8 µM

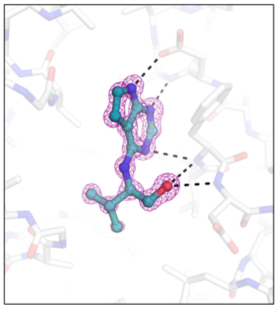

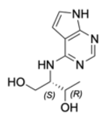

**AVI-3730** (I) | (*R*,*S*)
7HDK | (*R*,*S*)
63%

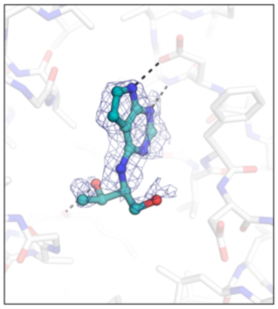

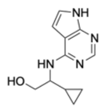

**AVI-3651** (I) | rac.
7HDZ | (*S*)
85% | 48 µM

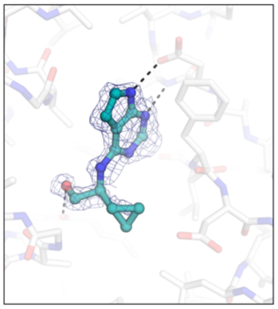

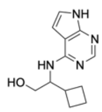

**AVI-3638** (I) | rac.
7HDP | (*S*)
77%

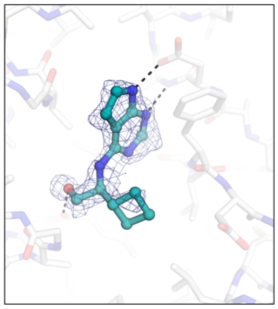

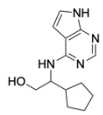

**AVI-3649** (I) | rac.
7HEZ | (*S*)
68%

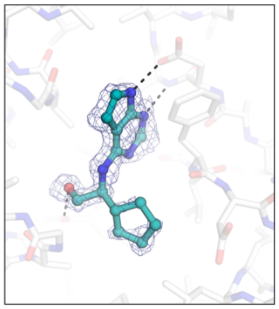

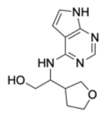

**AVI-3679** (I) | diast.
7HEJ | (*S*,*R*)
35%

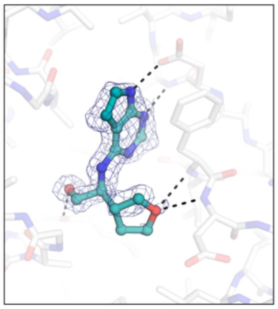

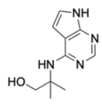

**AVI-5266** (I)
7HFX
56% | 110 µM

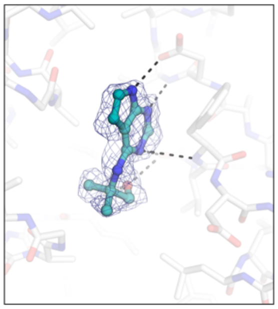

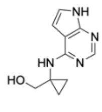

**AVI-3688** (I)
7HEP
72%

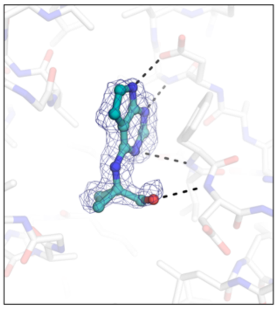

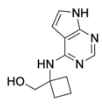

**AVI-3670** (I)
7HED
87% | 39 µM

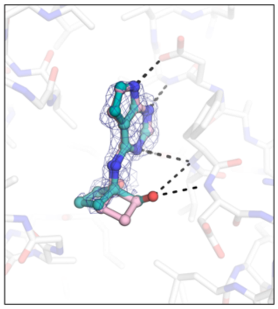

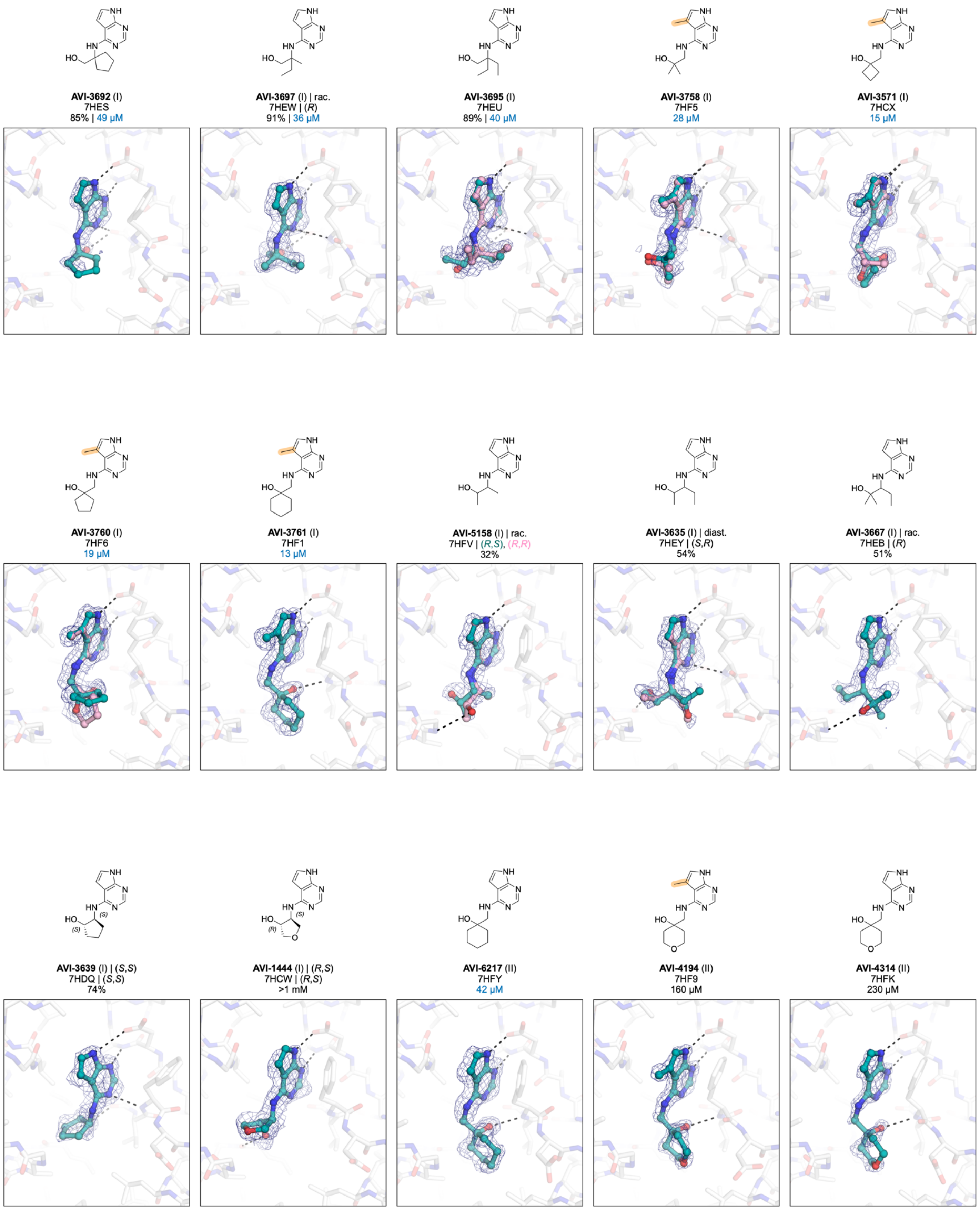
AVI-3692 (I)
7HES
85% | 49 µM
AVI-3697 (I) | rac.
7HEW | (R)
91% | 36 µM
AVI-3695 (I)
7HEU
89% | 40 µM
AVI-3758 (I)
7HF5
28 µM
AVI-3571 (I)
7HCX
15 µM
AVI-3760 (I)
7HF6
19 µM
AVI-3761 (I)
7HF1
13 µM
AVI-5158 (I) | rac.
7HFV | (R,S), (R,R)
32%
AVI-3635 (I) | diast.
7HEY | (S,R)
54%
AVI-3667 (I) | rac.
7HEB | (R)
51%
AVI-3639 (I) | (S,S)
7HDQ | (S,S)
74%
AVI-1444 (I) | (R,S)
7HCW | (R,S)
>1 mM
AVI-6217 (II)
7HFY
42 µM
AVI-4194 (II)
7HF9
160 µM
AVI-4314 (II)
7HFK
230 µM

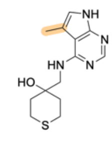

**AVI-4195** (II)
7HFA
52 μM

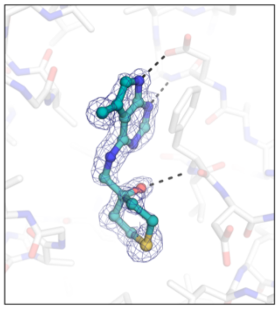

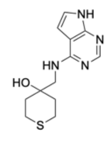

**AVI-4319** (II)
7HFL
170 μM

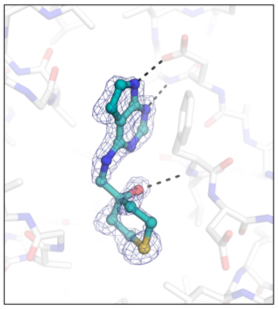

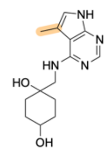

**AVI-4329** (II) | diast.
7HF8 | (*r*,*r*)
>1 mM

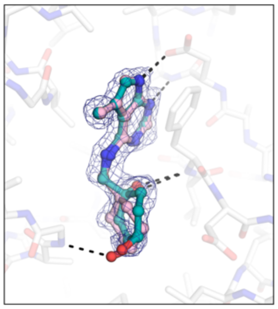

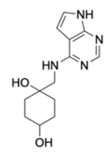

**AVI-6220** (II) | diast.
7HFZ | (*r*,*r*)
260 μM

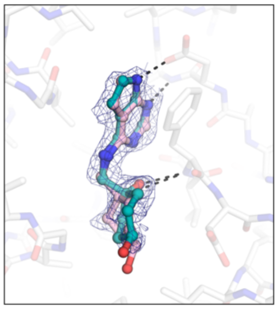

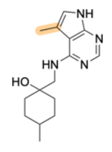

**AVI-4331** (II) | diast.
7HFB | (*r*,*r*)
210 μM

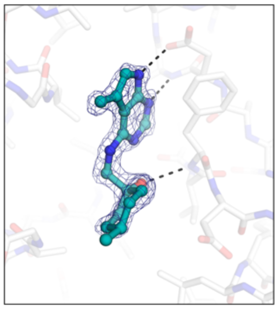

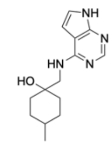

**AVI-4312** (II) | diast.
7HFI | (*r*,*r*)
21 μM

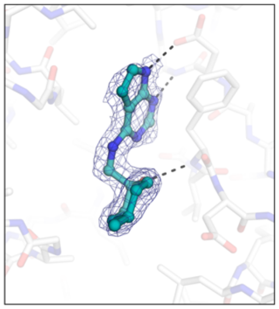

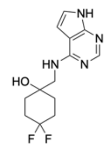

**AVI-4308** (II)
7HFG
140 μM

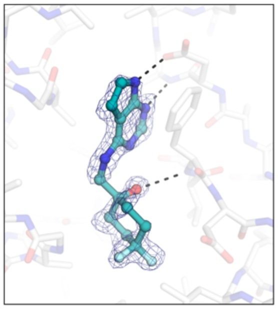

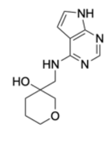

**AVI-4311** (II) | rac.
7HFH | (*R*)
170 μM

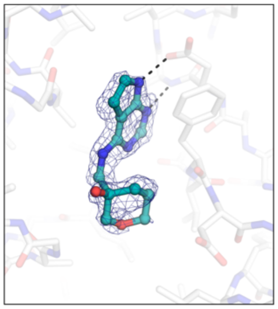

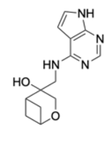

**AVI-4313** (II) | rac..
7HFJ | (*S*)
130 μM

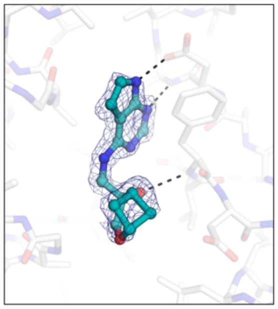

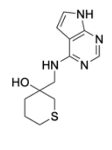

**AVI-4322** (II) | rac.
7HFN | (*R*)
53 μM

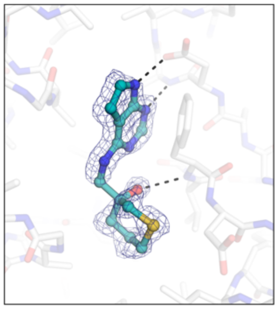

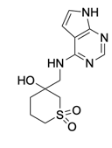

**AVI-4317** (II) | rac.
9D6I | (*R*)
110 μM

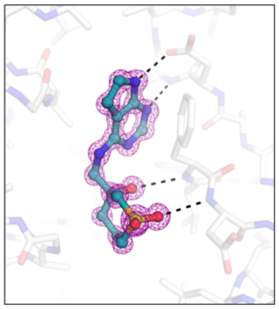

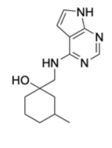

**AVI-4307** (II) | diast.
7HFF | (*R*,*S*)
40 μM

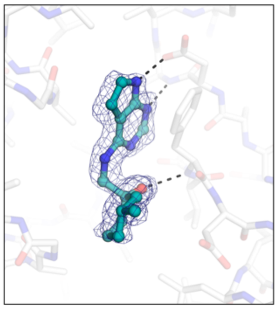

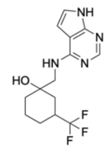

**AVI-4309** (II) | diast.
7HFO | (*S*,*R*)
150 μM

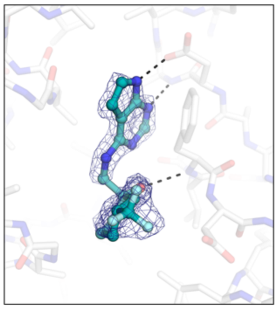

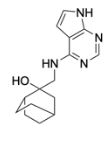

**AVI-4320** (II) | rac.
7HFM | (*R*)
83 μM

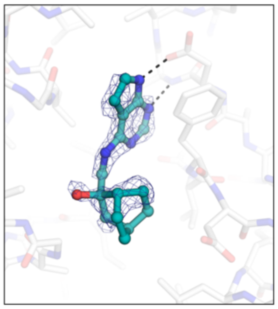

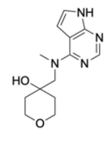

**AVI-4318** (II)
7HFP
71 μM

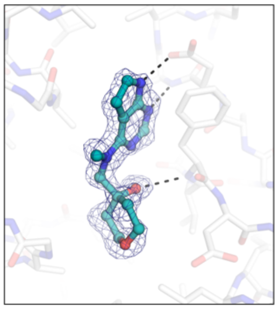

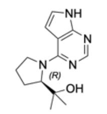

**AVI-607** (I) | (*R*)
9D6B | (*R*)
100% | 5.1 µM

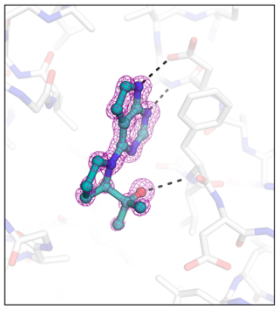

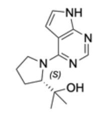

**AVI-1445** (II) | (*S*)
7HD6 | (*S*)
75% | 110 µM

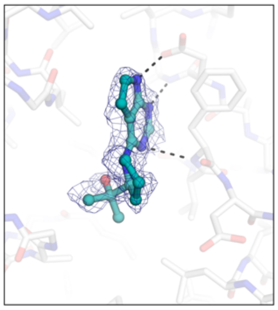

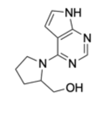

**AVI-3685** (II) | rac.
7HEN | (*S*)
47%

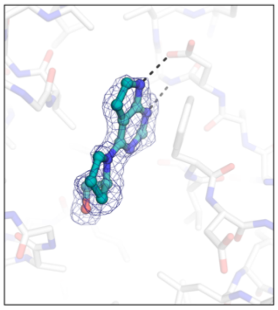

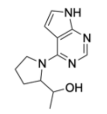

**AVI-4062** (II) | diast.
7HF2 | (*R*,*R*)
55 µM

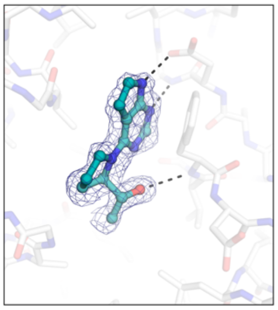

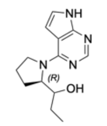

**AVI-3666** (II) | diast.
7HEA | (*R*,*R*)
87% | 46 µM

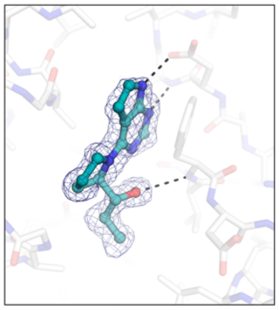

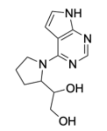

**AVI-3701** (II) | diast.
7HCY | (*R*,*R*)
54%

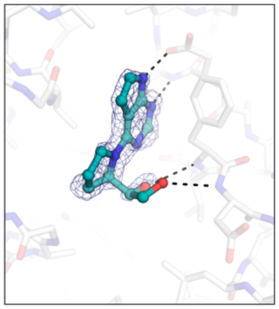

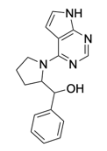

**AVI-3652** (II) | diast.
7HE0 | (*R*,*R*)
35%

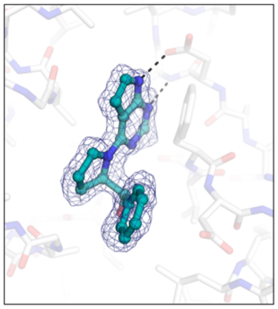

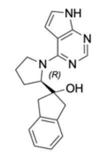

**AVI-3731** (II) | (*R*)
7HDL | (*R*)
100% | 3.7 µM

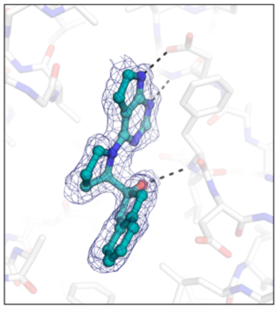

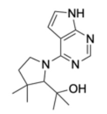

**AVI-4126** (II) | rac.
7HFC | (*R*), (*S*)
100 µM

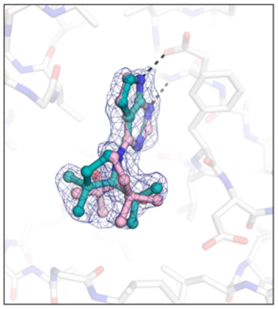

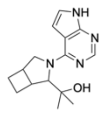

**AVI-4128** (II) | diast.
7HFE | (*S*,*R*,*R*)
24 µM

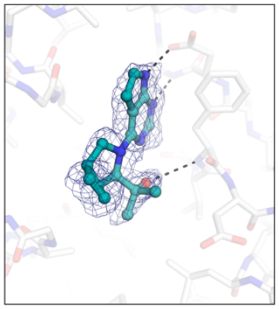

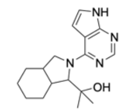

**AVI-4127** (II) | diast.
7HFD | (*S*,*S*,*R*), (*S*,*R*,*S*)
130 µM

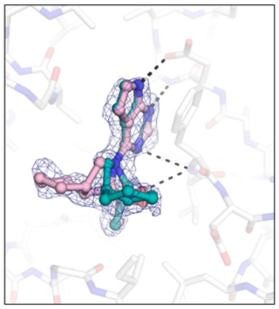

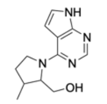

**AVI-3655** (II) | diast.
7HE1 | (*S*,*S*)
75%

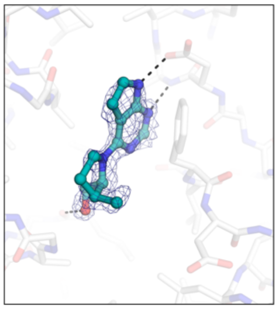

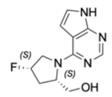

**AVI-609** (I) | (*S*,*S*)
7HCS | (*S*,*S*)
52%

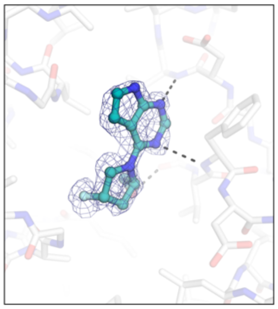

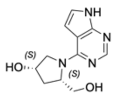

**AVI-610** (I) | (*S*,*S*)
7HCV | (*R*,*R*)
56% | 160 µM

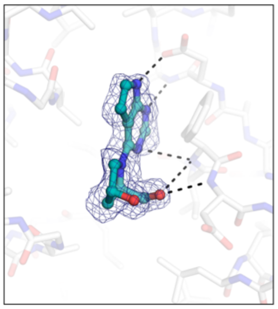

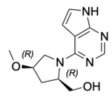

**AVI-3656** (II) | (*R*,*R*)
7HE2 | (*R*,*R*)
41%

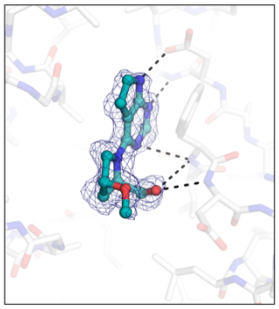

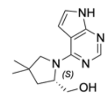

**AVI-3704** (II) | (*S*)
7HD1 | (*S*)
90% | 29 μM

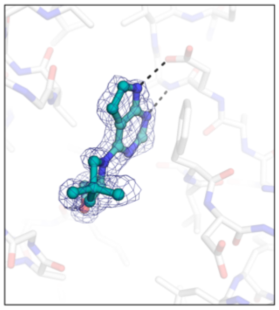

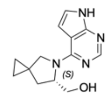

**AVI-3710** (II) | (*S*)
7HD7 | (*S*), (*S*)
57%

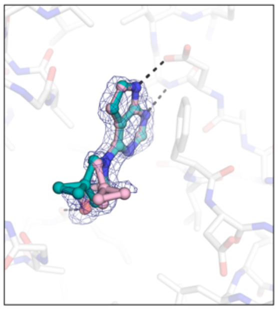

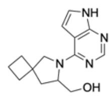

**AVI-3728** (II) | rac.
7HDJ | (*S*), (*S*)
79%

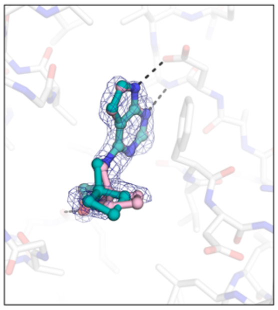

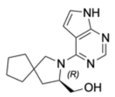

**AVI-3708** (II) | rac.
7HD4 | (*S*)
90% | 27 μM

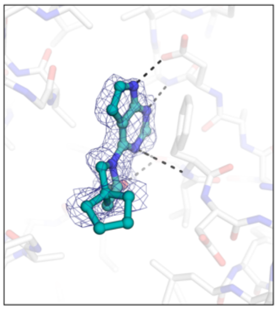

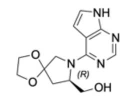

**AVI-3702** (II) | (*R*)
7HCZ | (*R*)
17%

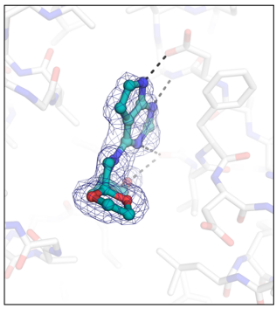

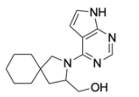

**AVI-3727** (II) | rac.
7HDI | (*S*)
67%

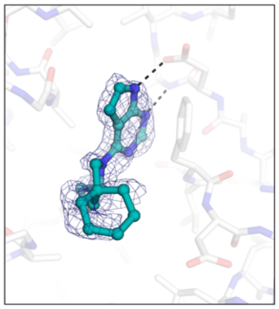

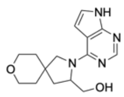

**AVI-3707** (II) | rac.
7HD3 | (*S*)
39%

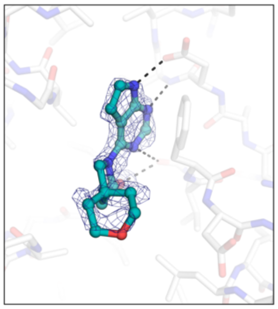

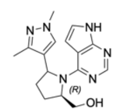

**AVI-3733** (II) | diast.
7HEX | (*R*,*S*)
44%

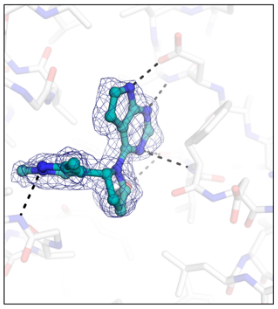

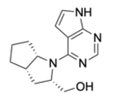

**AVI-3720** (II) | rac.
7HDE | (*S*,*S*,*S*)
92% | 28 μM

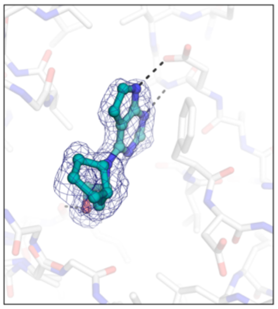

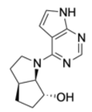

**AVI-3664** (II) | rac.
7HE8 | (*S*,*R*,*S*)
93% | 28 μM

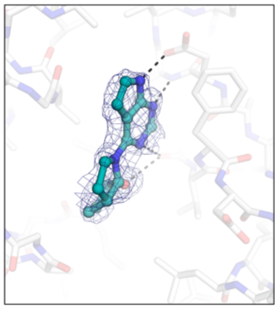

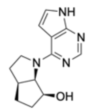

**AVI-5997** (II) | rac.
7HFR (*S*,*S*,*S*)
37 μM

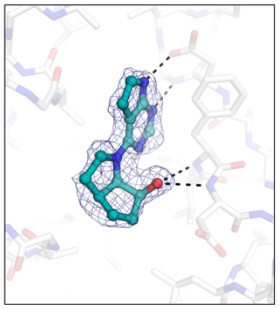

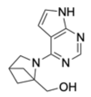

**AVI-3681** (II)
7HEK
54%

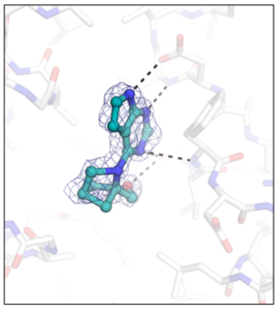

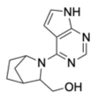

**AVI-3713** (II) | diast.
7HD9 | (*S*,*S*,*R*)
92% | 27 μM

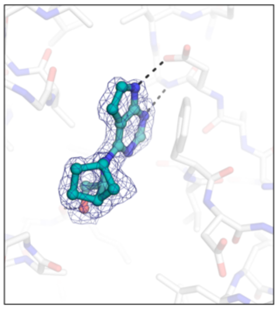

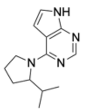

**AVI-5216** (II) | rac.
7HFW | (*R*)
35%

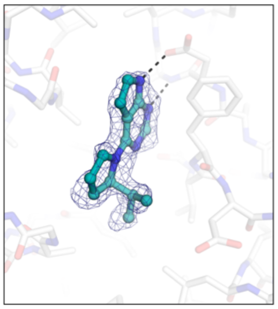

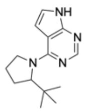

**AVI-6000** (II) | rac.
7HFT | (*R*)
>1 mM

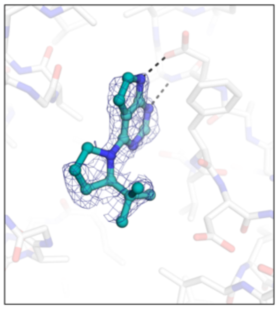

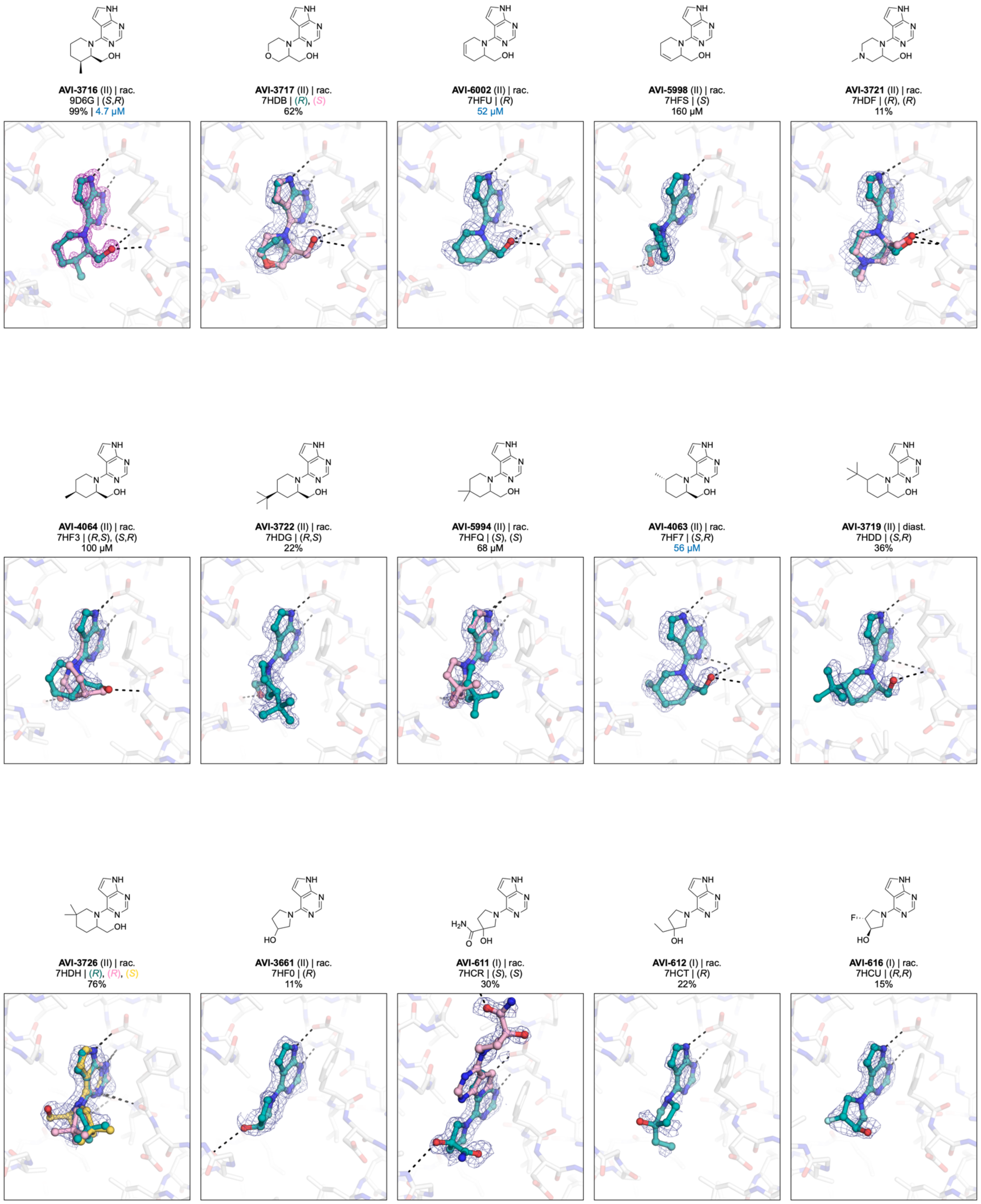
**AVI-3716** (II) | rac.
9D6G | (*S*,*R*)
99% | 4.7 µM
**AVI-3717** (II) | rac.
7HDB | (*R*), (*S*)
62%
**AVI-6002** (II) | rac.
7HFU | (*R*)
52 µM
**AVI-5998** (II) | rac.
7HFS | (*S*)
160 µM
**AVI-3721** (II) | rac.
7HDF | (*R*), (*R*)
11%
**AVI-4064** (II) | rac.
7HF3 | (*R*,*S*), (*S*,*R*)
100 µM
**AVI-3722** (II) | rac.
7HDG | (*R*,*S*)
22%
**AVI-5994** (II) | rac.
7HFQ | (*S*), (*S*)
68 µM
**AVI-4063** (II) | rac.
7HF7 | (*S*,*R*)
56 µM
**AVI-3719** (II) | diast.
7HDD | (*S*,*R*)
36%
**AVI-3726** (II) | rac.
7HDH | (*R*), (*R*), (*S*)
76%
**AVI-3661** (II) | rac.
7HF0 | (*R*)
11%
**AVI-611** (I) | rac.
7HCR | (*S*), (*S*)
30%
**AVI-612** (I) | rac.
7HCT | (*R*)
22%
**AVI-616** (I) | rac.
7HCU | (*R*,*R*)
15%

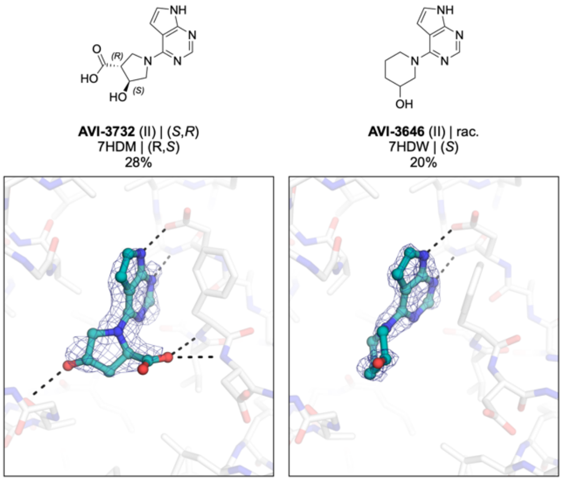


**Figure S2.** Chemical structures and electron density maps for the 137 X-ray crystal structures reported in this work. PanDDA event maps (blue mesh, 2 σ) or $F_O$-$F_C$ difference electron density maps prior to ligand placement (purple mesh, 5 σ) are contoured around ligands. Compounds were synthesized as pure isomers, as racamates (rac.) or as mixtures of diastereomers (diast.) and the stereochemistry of the modeled compound is indicated. Results for the preliminary HTRF screen (% competition at 200 μM, mean of three technical replicates) and the confirmatory dose-response screen ($IC_{50}$, best fit value of 3-4 technical replicates) are indicated.

A

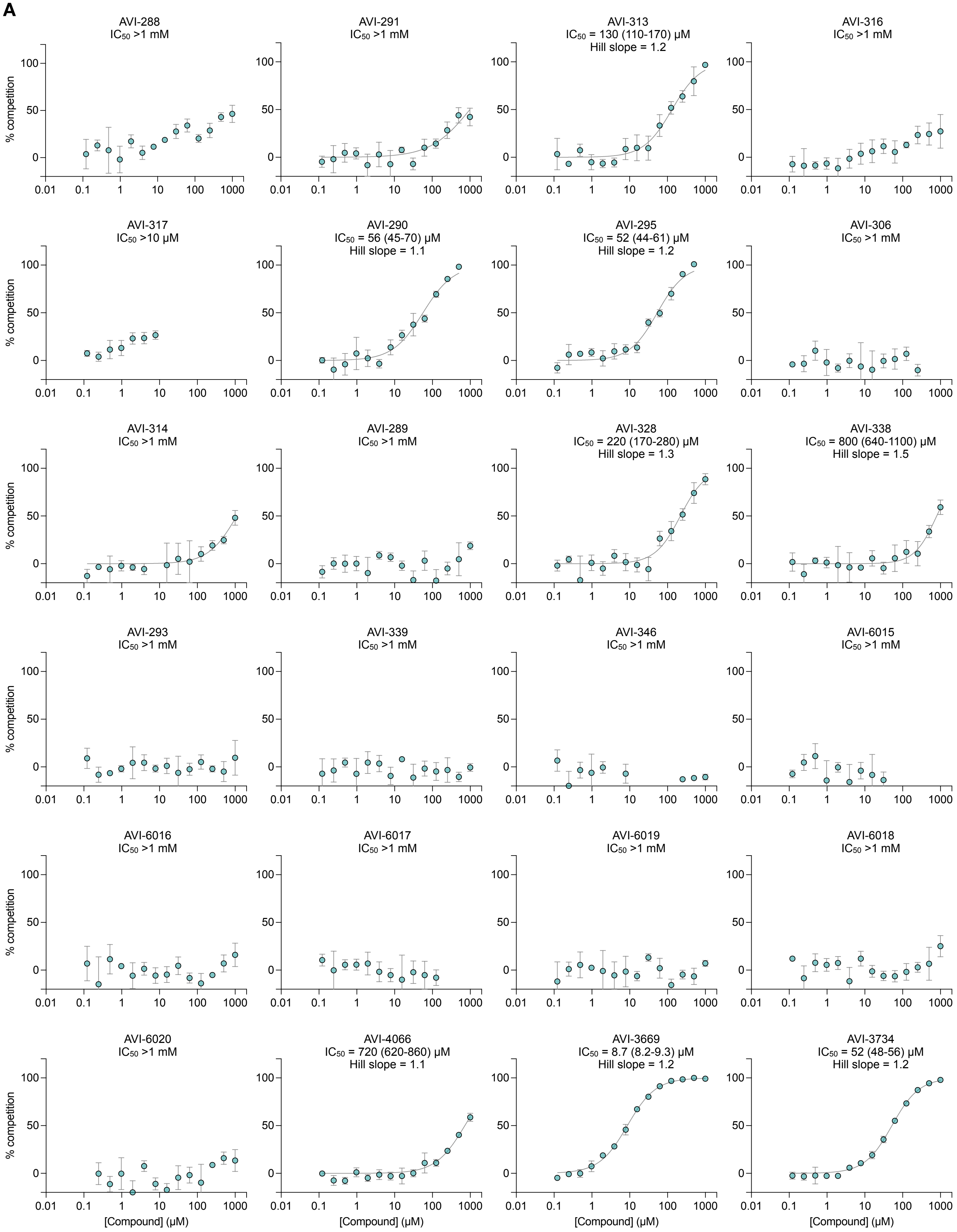
AVI-288
$IC_{50}$ >1 mM
AVI-291
$IC_{50}$ >1 mM
AVI-313
$IC_{50}$ = 130 (110-170) μM
Hill slope = 1.2
AVI-316
$IC_{50}$ >1 mM
AVI-317
$IC_{50}$ >10 μM
AVI-290
$IC_{50}$ = 56 (45-70) μM
Hill slope = 1.1
AVI-295
$IC_{50}$ = 52 (44-61) μM
Hill slope = 1.2
AVI-306
$IC_{50}$ >1 mM
AVI-314
$IC_{50}$ >1 mM
AVI-289
$IC_{50}$ >1 mM
AVI-328
$IC_{50}$ = 220 (170-280) μM
Hill slope = 1.3
AVI-338
$IC_{50}$ = 800 (640-1100) μM
Hill slope = 1.5
AVI-293
$IC_{50}$ >1 mM
AVI-339
$IC_{50}$ >1 mM
AVI-346
$IC_{50}$ >1 mM
AVI-6015
$IC_{50}$ >1 mM
AVI-6016
$IC_{50}$ >1 mM
AVI-6017
$IC_{50}$ >1 mM
AVI-6019
$IC_{50}$ >1 mM
AVI-6018
$IC_{50}$ >1 mM
AVI-6020
$IC_{50}$ >1 mM
AVI-4066
$IC_{50}$ = 720 (620-860) μM
Hill slope = 1.1
AVI-3669
$IC_{50}$ = 8.7 (8.2-9.3) μM
Hill slope = 1.2
AVI-3734
$IC_{50}$ = 52 (48-56) μM
Hill slope = 1.2
% competition
[Compound] (μM)

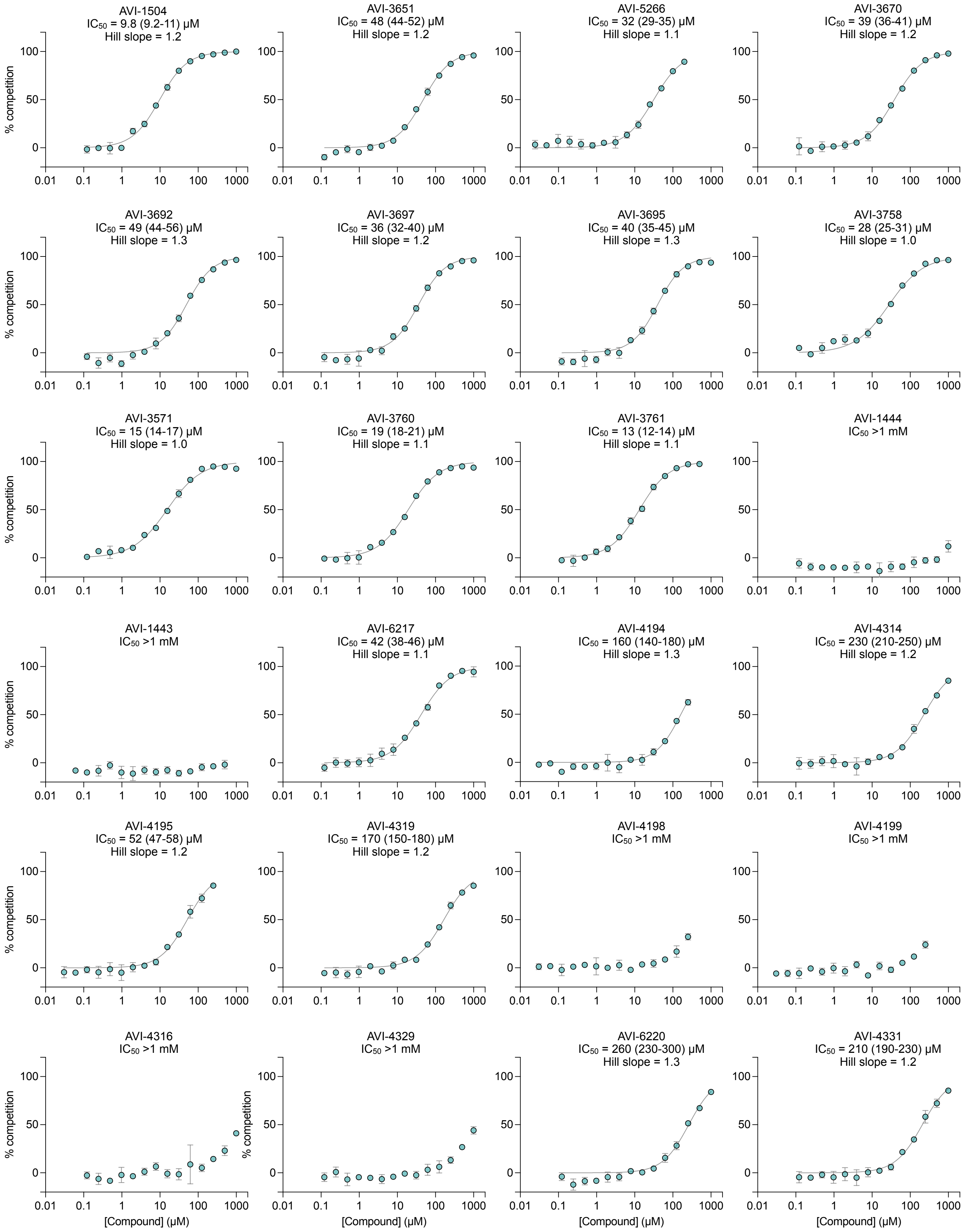
AVI-1504
$IC_{50}$ = 9.8 (9.2-11) μM
Hill slope = 1.2
AVI-3651
$IC_{50}$ = 48 (44-52) μM
Hill slope = 1.2
AVI-5266
$IC_{50}$ = 32 (29-35) μM
Hill slope = 1.1
AVI-3670
$IC_{50}$ = 39 (36-41) μM
Hill slope = 1.2
AVI-3692
$IC_{50}$ = 49 (44-56) μM
Hill slope = 1.3
AVI-3697
$IC_{50}$ = 36 (32-40) μM
Hill slope = 1.2
AVI-3695
$IC_{50}$ = 40 (35-45) μM
Hill slope = 1.3
AVI-3758
$IC_{50}$ = 28 (25-31) μM
Hill slope = 1.0
AVI-3571
$IC_{50}$ = 15 (14-17) μM
Hill slope = 1.0
AVI-3760
$IC_{50}$ = 19 (18-21) μM
Hill slope = 1.1
AVI-3761
$IC_{50}$ = 13 (12-14) μM
Hill slope = 1.1
AVI-1444
$IC_{50}$ >1 mM
AVI-1443
$IC_{50}$ >1 mM
AVI-6217
$IC_{50}$ = 42 (38-46) μM
Hill slope = 1.1
AVI-4194
$IC_{50}$ = 160 (140-180) μM
Hill slope = 1.3
AVI-4314
$IC_{50}$ = 230 (210-250) μM
Hill slope = 1.2
AVI-4195
$IC_{50}$ = 52 (47-58) μM
Hill slope = 1.2
AVI-4319
$IC_{50}$ = 170 (150-180) μM
Hill slope = 1.2
AVI-4198
$IC_{50}$ >1 mM
AVI-4199
$IC_{50}$ >1 mM
AVI-4316
$IC_{50}$ >1 mM
AVI-4329
$IC_{50}$ >1 mM
AVI-6220
$IC_{50}$ = 260 (230-300) μM
Hill slope = 1.3
AVI-4331
$IC_{50}$ = 210 (190-230) μM
Hill slope = 1.2
% competition
0
50
100
0.01
0.1
1
10
100
1000
[Compound] (μM)

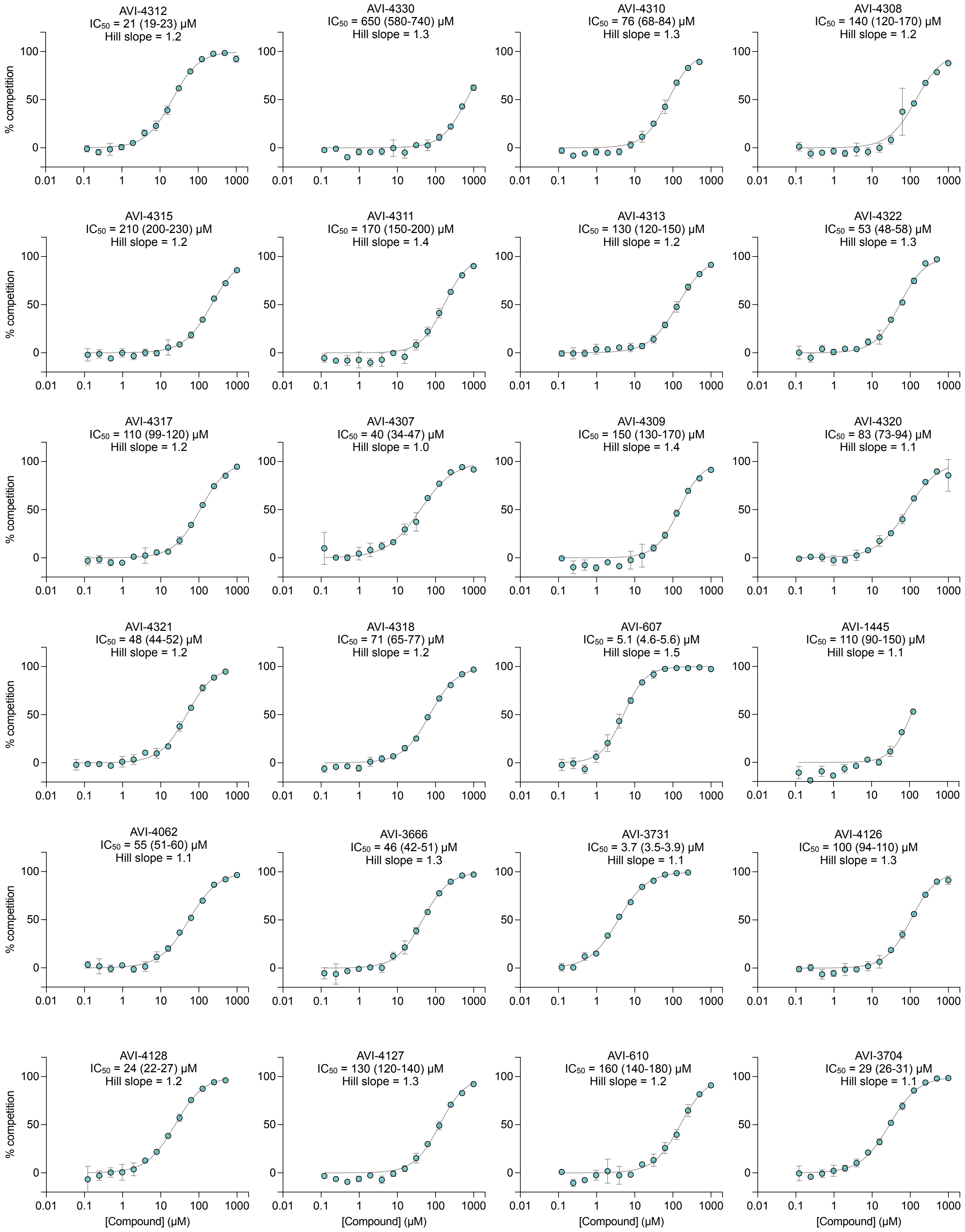
AVI-4312
$IC_{50}$ = 21 (19-23) µM
Hill slope = 1.2
AVI-4330
$IC_{50}$ = 650 (580-740) µM
Hill slope = 1.3
AVI-4310
$IC_{50}$ = 76 (68-84) µM
Hill slope = 1.3
AVI-4308
$IC_{50}$ = 140 (120-170) µM
Hill slope = 1.2
AVI-4315
$IC_{50}$ = 210 (200-230) µM
Hill slope = 1.2
AVI-4311
$IC_{50}$ = 170 (150-200) µM
Hill slope = 1.4
AVI-4313
$IC_{50}$ = 130 (120-150) µM
Hill slope = 1.2
AVI-4322
$IC_{50}$ = 53 (48-58) µM
Hill slope = 1.3
AVI-4317
$IC_{50}$ = 110 (99-120) µM
Hill slope = 1.2
AVI-4307
$IC_{50}$ = 40 (34-47) µM
Hill slope = 1.0
AVI-4309
$IC_{50}$ = 150 (130-170) µM
Hill slope = 1.4
AVI-4320
$IC_{50}$ = 83 (73-94) µM
Hill slope = 1.1
AVI-4321
$IC_{50}$ = 48 (44-52) µM
Hill slope = 1.2
AVI-4318
$IC_{50}$ = 71 (65-77) µM
Hill slope = 1.2
AVI-607
$IC_{50}$ = 5.1 (4.6-5.6) µM
Hill slope = 1.5
AVI-1445
$IC_{50}$ = 110 (90-150) µM
Hill slope = 1.1
AVI-4062
$IC_{50}$ = 55 (51-60) µM
Hill slope = 1.1
AVI-3666
$IC_{50}$ = 46 (42-51) µM
Hill slope = 1.3
AVI-3731
$IC_{50}$ = 3.7 (3.5-3.9) µM
Hill slope = 1.1
AVI-4126
$IC_{50}$ = 100 (94-110) µM
Hill slope = 1.3
AVI-4128
$IC_{50}$ = 24 (22-27) µM
Hill slope = 1.2
AVI-4127
$IC_{50}$ = 130 (120-140) µM
Hill slope = 1.3
AVI-610
$IC_{50}$ = 160 (140-180) µM
Hill slope = 1.2
AVI-3704
$IC_{50}$ = 29 (26-31) µM
Hill slope = 1.1
% competition
[Compound] (µM)
0
50
100
0.01
0.1
1
10
100
1000

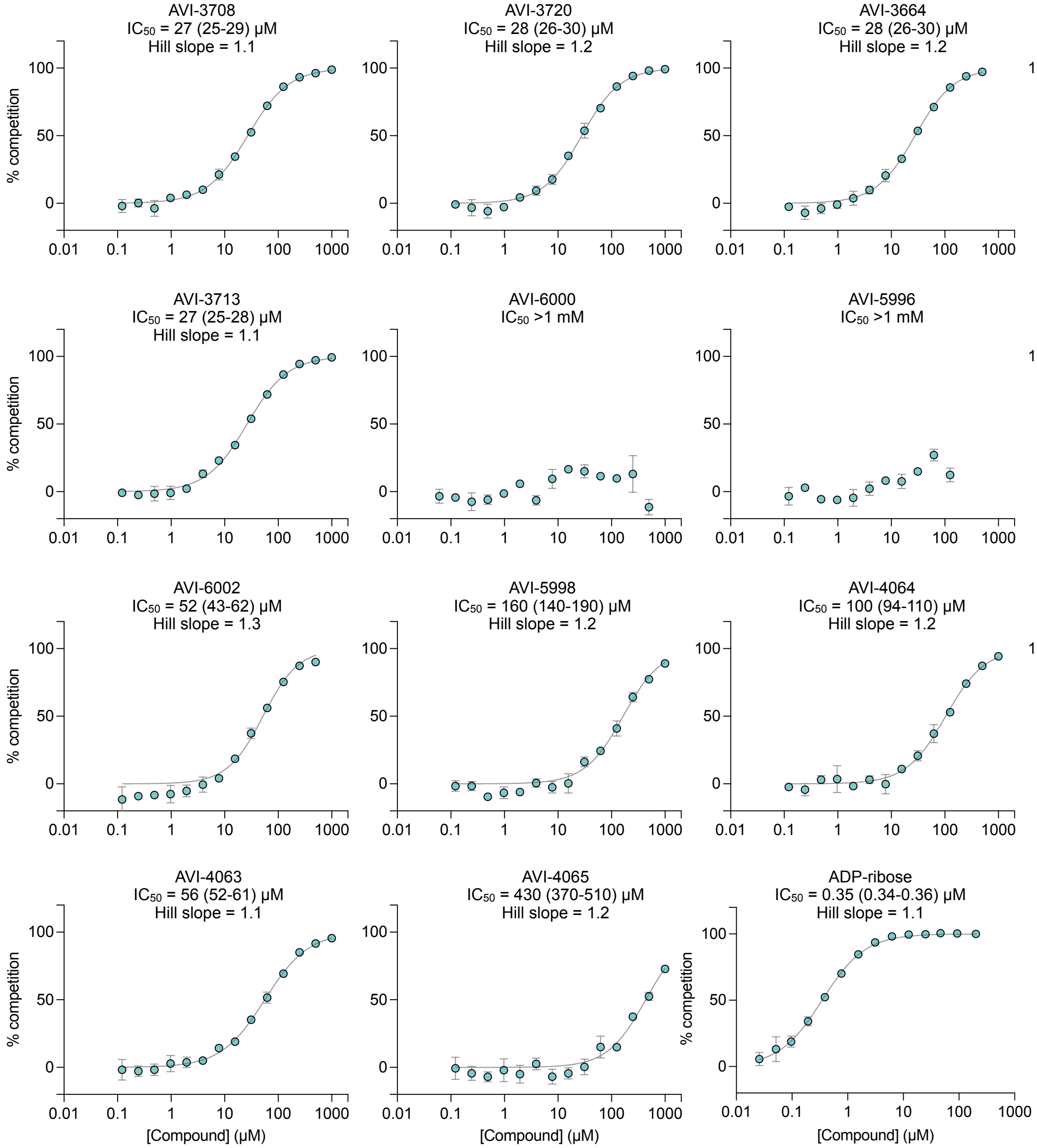
AVI-3708
IC50 = 27 (25-29) µM
Hill slope = 1.1
AVI-3720
IC50 = 28 (26-30) µM
Hill slope = 1.2
AVI-3664
IC50 = 28 (26-30) µM
Hill slope = 1.2
AVI-5997
IC50 = 37 (34-40) µM
Hill slope = 1.2
AVI-3713
IC50 = 27 (25-28) µM
Hill slope = 1.1
AVI-6000
IC50 >1 mM
AVI-5996
IC50 >1 mM
AVI-3716
IC50 = 4.7 (4.4-5.0) µM
Hill slope = 1.2
AVI-6002
IC50 = 52 (43-62) µM
Hill slope = 1.3
AVI-5998
IC50 = 160 (140-190) µM
Hill slope = 1.2
AVI-4064
IC50 = 100 (94-110) µM
Hill slope = 1.2
AVI-5994
IC50 = 68 (60-77) µM
Hill slope = 1.2
AVI-4063
IC50 = 56 (52-61) µM
Hill slope = 1.1
AVI-4065
IC50 = 430 (370-510) µM
Hill slope = 1.2
ADP-ribose
IC50 = 0.35 (0.34-0.36) µM
Hill slope = 1.1
% competition
[Compound] (µM)
0
50
100
0.01
0.1
1
10
100
1000

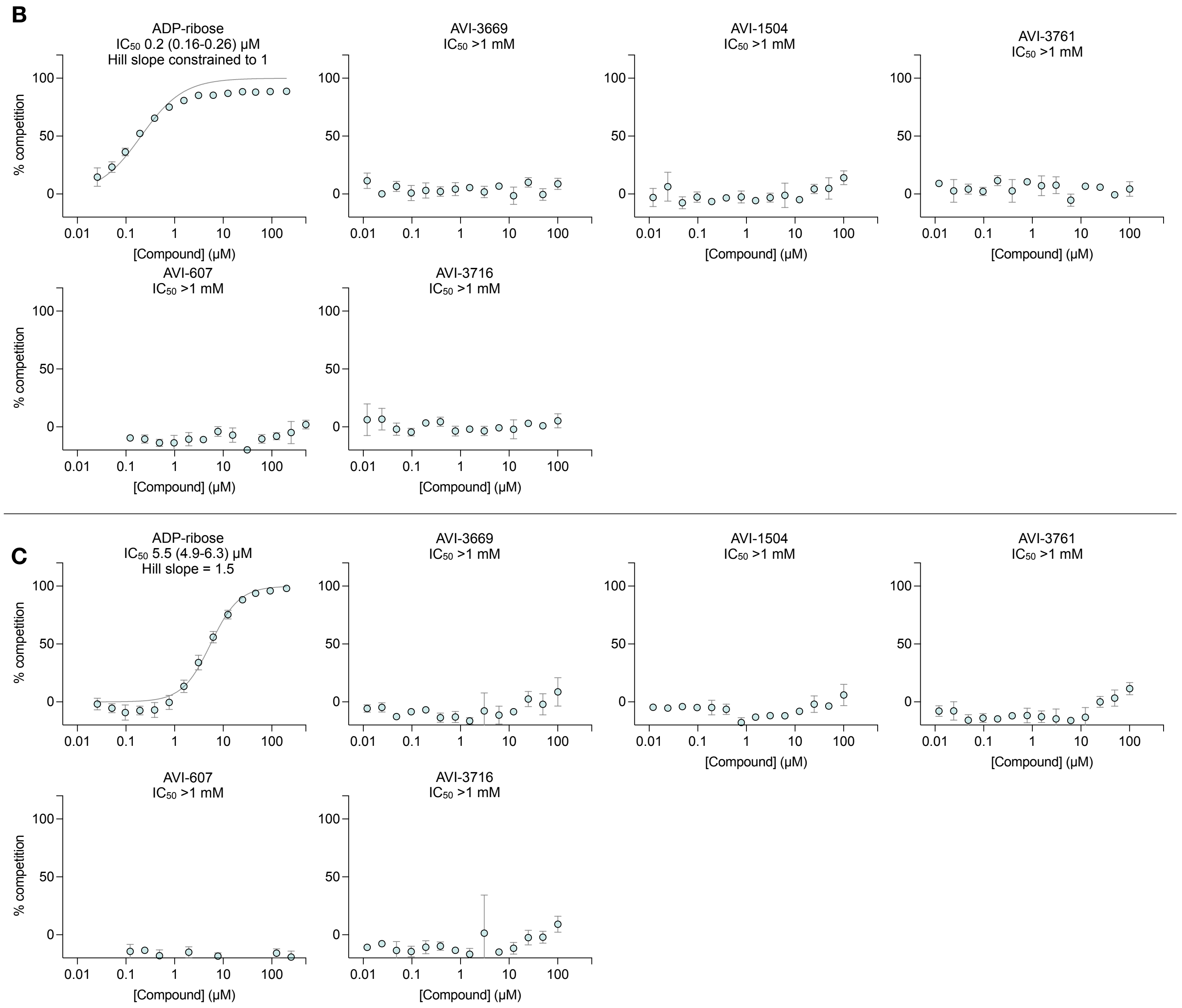


**Figure S3.** HTRF binding curves for all compounds tested. Curves are shown for SARS-Cov-2 Mac1 (A) and the human macrodomains MacroD2 (**B**) and Targ1 (**C**). Data points are mean ± SD for 3-4 technical replicates at each compound concentration, except for ADP-ribose with Mac1, where there were 12 technical replicates. The $IC_{50}$ value was determined by non-linear regression of a four-parameter sigmoidal dose-response equation in GraphPad Prism (version 10.1.1). The best fit value is indicated along with the 95% confidence interval and the Hill slope. The Hill slope was constrained to 1 for the MacroD2 ADP-ribose binding curve.

A

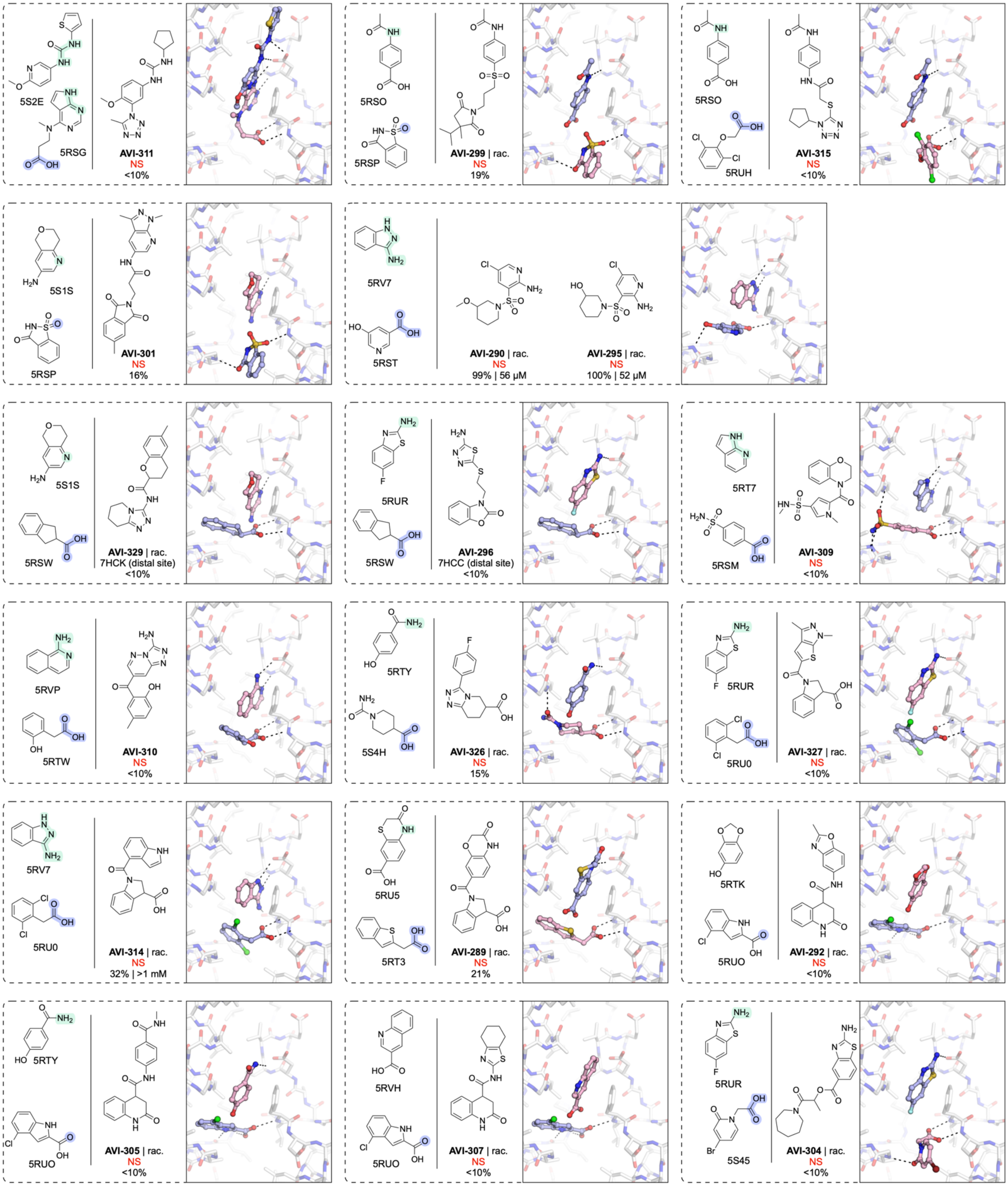
5S2E
5RSG
AVI-311
NS
<10%
5RSO
5RSP
AVI-299 | rac.
NS
19%
5RSO
5RUH
AVI-315
NS
<10%
5S1S
5RSP
AVI-301
NS
16%
5RV7
5RST
AVI-290 | rac.
NS
99% | 56 μM
AVI-295 | rac.
NS
100% | 52 μM
5S1S
5RSW
AVI-329 | rac.
7HCK (distal site)
<10%
5RUR
5RSW
AVI-296
7HCC (distal site)
<10%
5RT7
5RSM
AVI-309
NS
<10%
5RVP
5RTW
AVI-310
NS
<10%
5RTY
5S4H
AVI-326 | rac.
NS
15%
5RUR
5RU0
AVI-327 | rac.
NS
<10%
5RV7
5RU0
AVI-314 | rac.
NS
32% | >1 mM
5RU5
5RT3
AVI-289 | rac.
NS
21%
5RTK
5RUO
AVI-292 | rac.
NS
<10%
5RTY
5RUO
AVI-305 | rac.
NS
<10%
5RVH
5RUO
AVI-307 | rac.
NS
<10%
5RUR
5S45
AVI-304 | rac.
NS
<10%

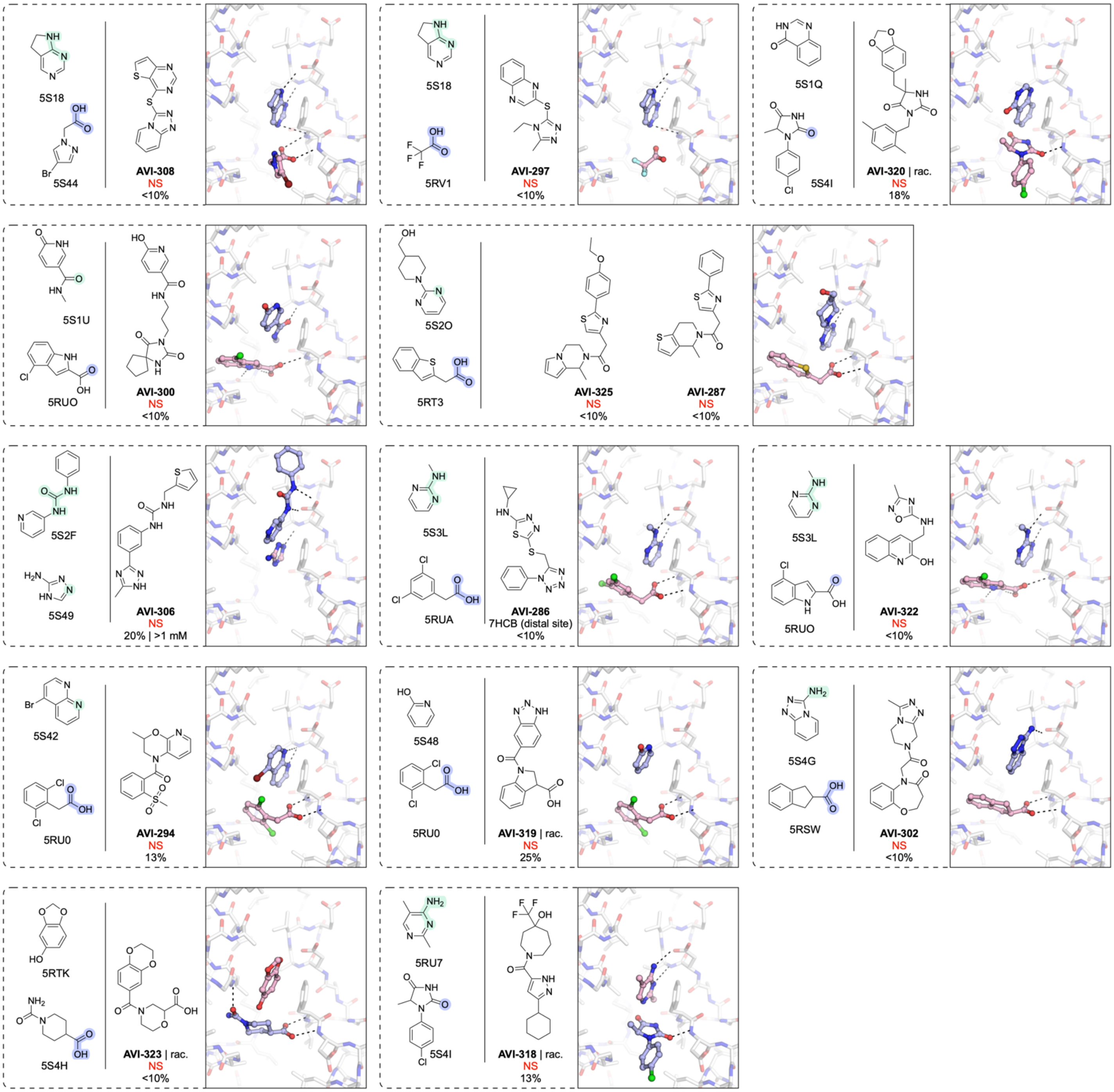
5S18
5S44
AVI-308
NS
<10%
5S18
5RV1
AVI-297
NS
<10%
5S1Q
5S4I
AVI-320 | rac.
NS
18%
5S1U
5RUO
AVI-300
NS
<10%
5S2O
5RT3
AVI-325
NS
<10%
AVI-287
NS
<10%
5S2F
5S49
AVI-306
NS
20% | >1 mM
5S3L
5RUA
AVI-286
7HCB (distal site)
<10%
5S3L
5RUO
AVI-322
NS
<10%
5S42
5RU0
AVI-294
NS
13%
5S48
5RU0
AVI-319 | rac.
NS
25%
5S4G
5RSW
AVI-302
NS
<10%
5RTK
5S4H
AVI-323 | rac.
NS
<10%
5RU7
5S4I
AVI-318 | rac.
NS
13%

**B**

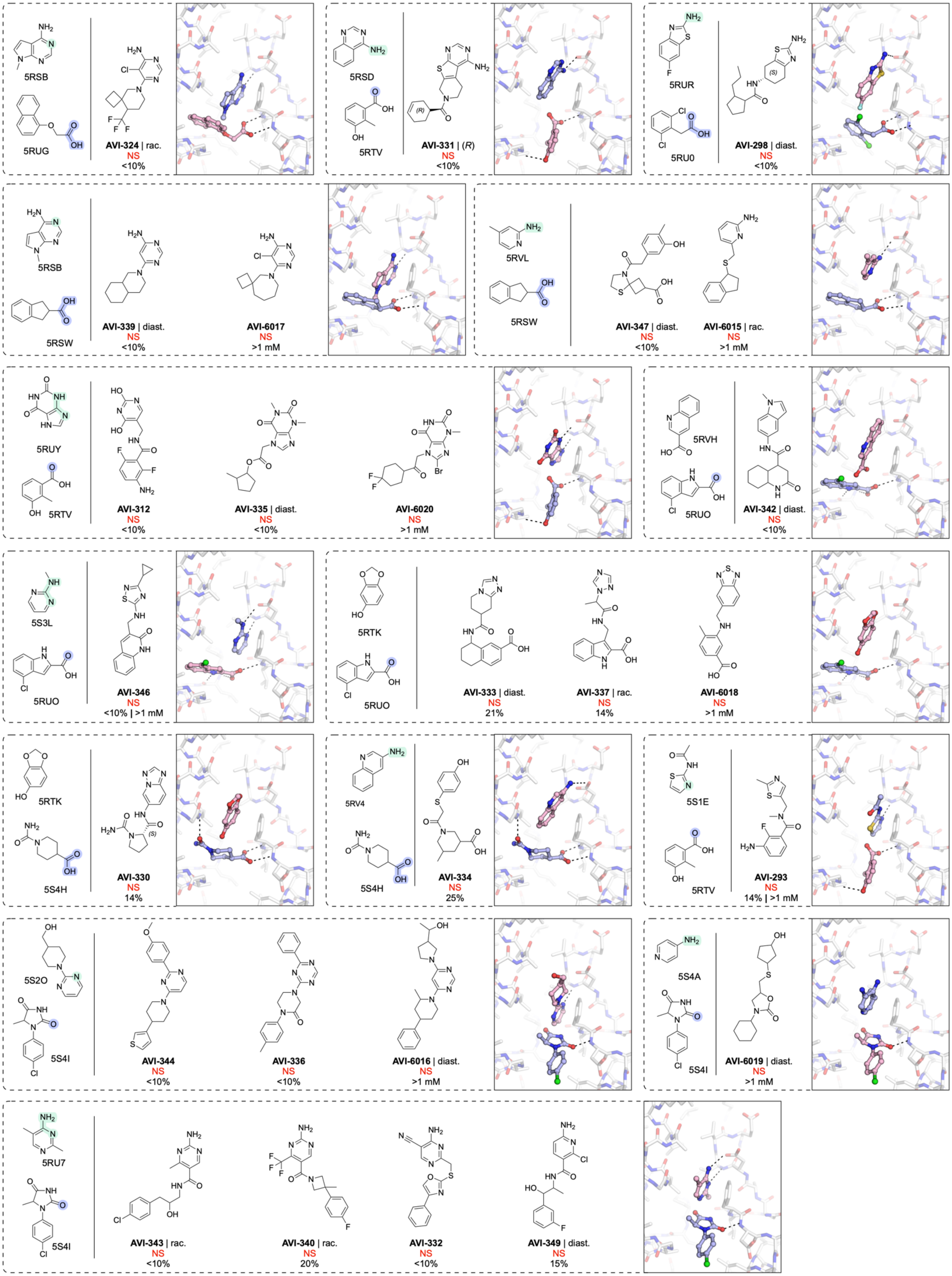

5RSB
5RUG
AVI-324 | rac.
NS
<10%
5RSD
5RTV
AVI-331 | (R)
NS
<10%
5RUR
5RU0
AVI-298 | diast.
NS
<10%
5RSB
5RSW
AVI-339 | diast.
NS
<10%
AVI-6017
NS
>1 mM
5RVL
5RSW
AVI-347 | diast.
NS
<10%
AVI-6015 | rac.
NS
>1 mM
5RUY
5RTV
AVI-312
NS
<10%
AVI-335 | diast.
NS
<10%
AVI-6020
NS
>1 mM
5RVH
5RUO
AVI-342 | diast.
NS
<10%
5S3L
5RUO
AVI-346
NS
<10% | >1 mM
5RTK
5RUO
AVI-333 | diast.
NS
21%
AVI-337 | rac.
NS
14%
AVI-6018
NS
>1 mM
5RTK
5S4H
AVI-330
NS
14%
5RV4
5S4H
AVI-334
NS
25%
5S1E
5RTV
AVI-293
NS
14% | >1 mM
5S2O
5S4I
AVI-344
NS
<10%
AVI-336
NS
<10%
AVI-6016 | diast.
NS
>1 mM
5S4A
5S4I
AVI-6019 | diast.
NS
>1 mM
5RU7
5S4I
AVI-343 | rac.
NS
<10%
AVI-340 | rac.
NS
20%
AVI-332
NS
<10%
AVI-349 | diast.
NS
15%

**Figure S4.** Compounds synthesized from the FrankenROCS (**A**) and the TS-FrankenROCS (**B**) pipelines where no X-ray crystal structure was determined. Compounds were purchased as pure isomers, as racamates (rac.) or mixtures of diastereomers (diast.). The X-ray crystal structures of the parent fragments (transparent blue/pink sticks) are shown. The fragment atoms making contact with the adenine and oxyanion subsites are shaded green and blue respectively. Hydrogen bonds between the fragments ligand and Mac1 are shown as dashed black lines. Results for the preliminary HTRF screen (% competition at 200 μM, mean of three technical replicates) and the confirmatory dose-response screen ($IC_{50}$, best fit value of 3-4 technical replicates) are indicated.

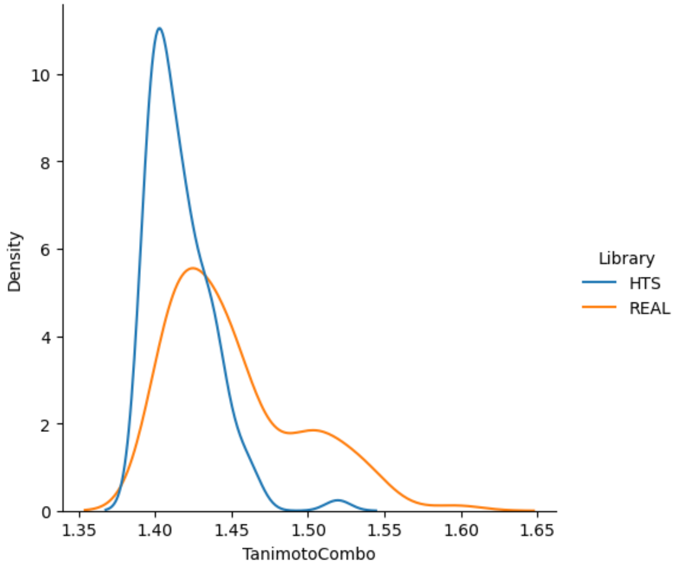


**Figure S5.** Comparison of the top 100 TanimotoCombo scores for the Enamine HTS collection versus the Enamine REAL database.

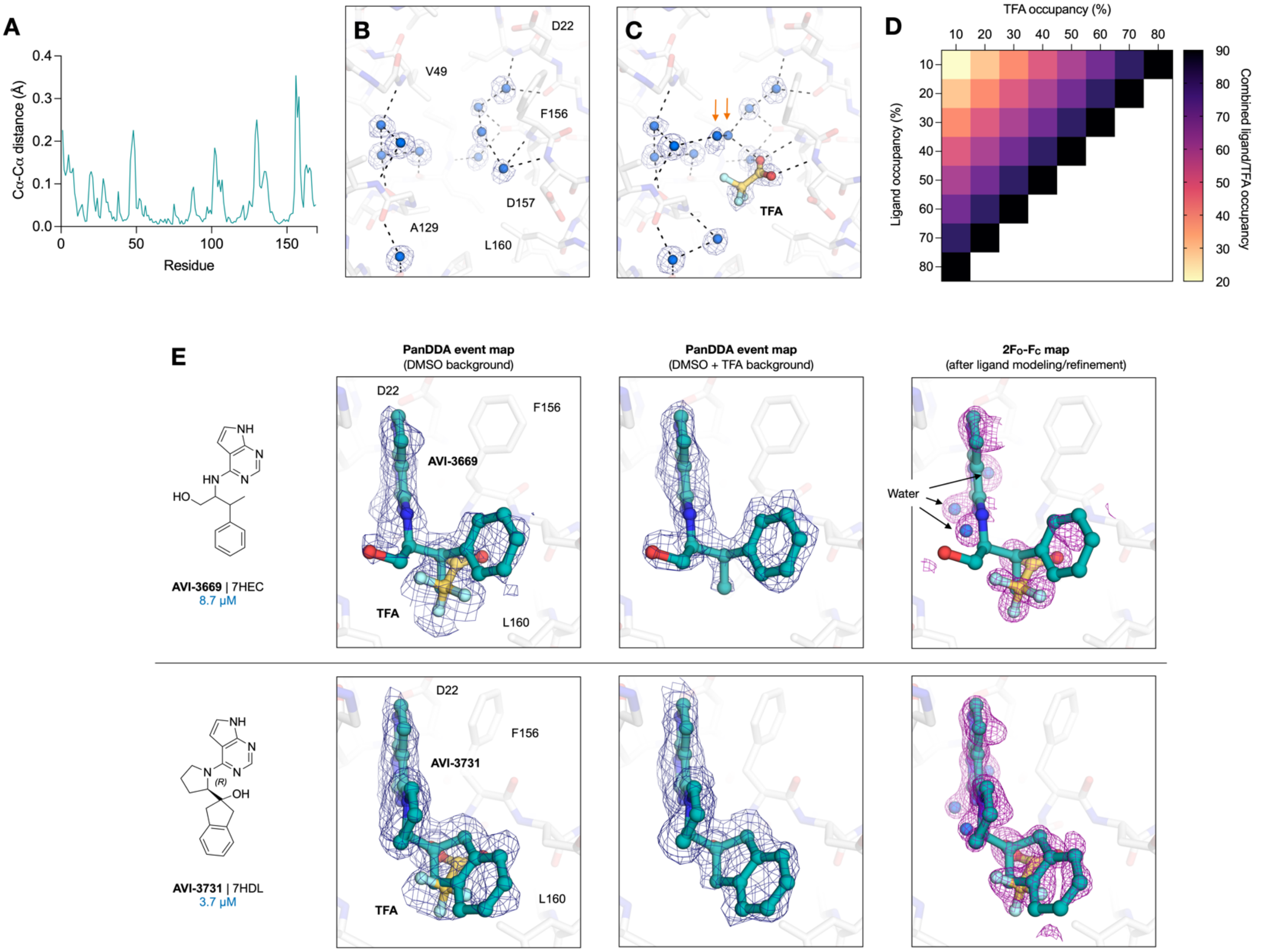


**Figure S6.** TFA subtraction improves the PanDDA map interpretation. (**A**) Pairwise Cα-Cα distances between the DMSO- and TFA-reference structures. (**B**) Average electron density maps calculated by PanDDA from 34 datasets collected from DMSO-soaked crystals with the DMSO reference structure shown (UCSF-P0075). Hydrogen bond networks with water molecules are shown with dashed black lines. (**C**) Average electron density maps calculated by PanDDA from 44 datasets collected from TFA/DMSO-soaked crystals with the TFA reference structure shown (UCSF-P3816). (**D**) Heat map showing the grid search performed to assess ligand/TFA occupancy. Each square represents a separate refinement run where the ligand/TFA occupancies were set to the indicated values. (**E**) Examples of ligands where the TFA density was removed in the PanDDA event maps (blue mesh, 2 σ) using the TFA/DMSO-background map. The $2F_O$-$F_C$ map (purple mesh, 1 σ) calculated with the final refined model are contoured around the ligands/TFA and overlapping water molecules.

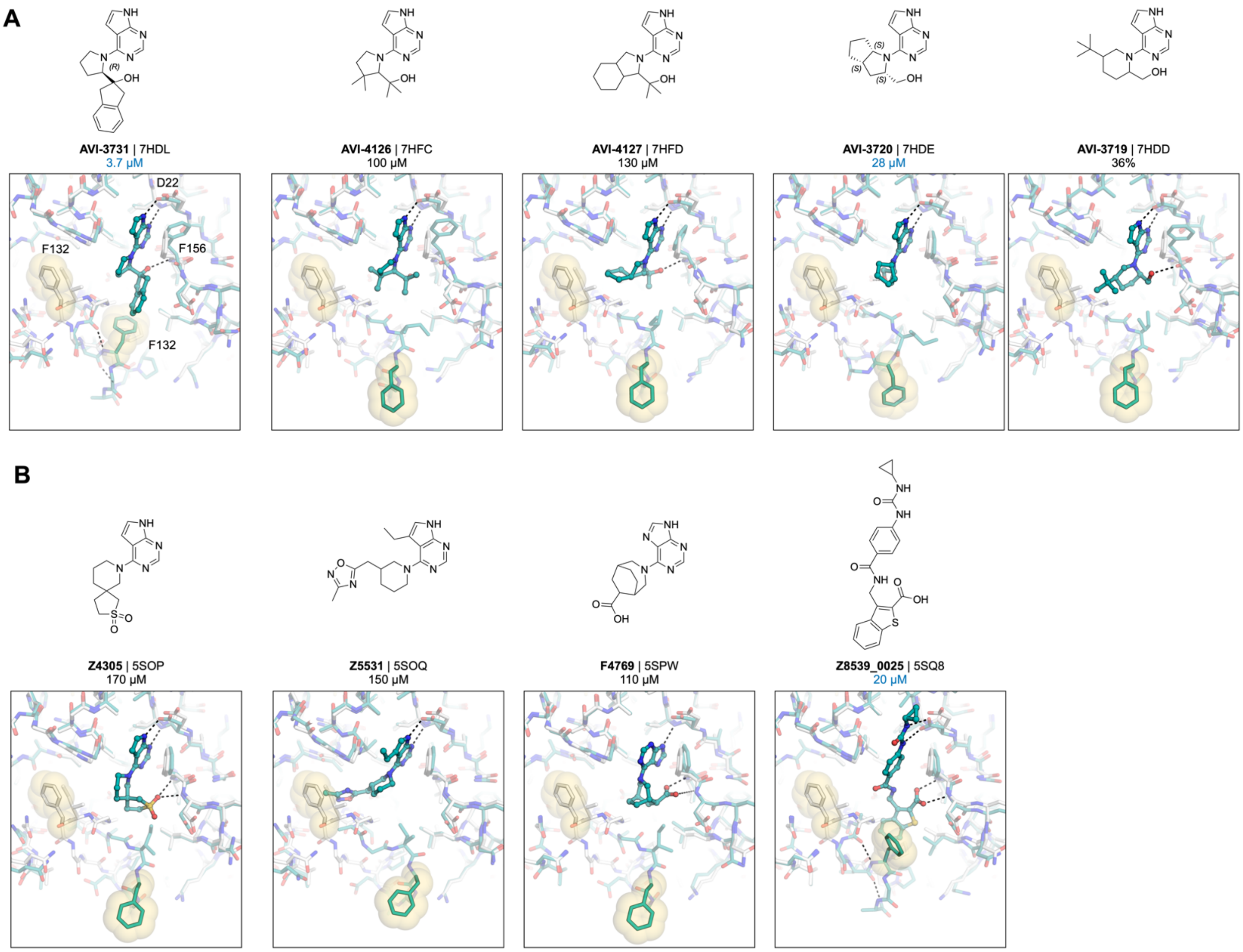


**Figure S7.** AVI-313 analogues stabilize an alternative, open state of the Phe132 loop. (**A**) Top: Chemical structures of AVI-131 ligands that stabilize the open state of Mac1 reported in this work. Bottom: alignment of the apo X-ray crystal structure in the $P4_3$ space group (PDB 7KQO, white sticks) with the open state crystal structure (teal sticks). The Phe132 side chain is shown with transparent yellow spheres. The Phe132 loop adopts a distinct conformation in the most potent open state binder (AVI-3731, $IC_{50}$ = 3.7 µM), where the Phe132 side-chain makes contact with the indane portion of the ligand and the Phe132 loop forms a short α-helix. (**B**) Same as (**A**) but showing open-state structures reported in (2). Although Z8539_0025 stabilizes a similar short α-helix compared to AVI-3731, the Phe132 side-chain is solvent exposed instead of making contact with the ligand.
